# Joint inference of discrete cell types and continuous type-specific variability in single-cell datasets with MMIDAS

**DOI:** 10.1101/2023.10.02.560574

**Authors:** Yeganeh Marghi, Rohan Gala, Fahimeh Baftizadeh, Uygar Sümbül

**Affiliations:** Allen Institute, 615 Westlake Ave N, Seattle, WA, USA; Paul G. Allen School of Computer Science & Engineering, University of Washington, Seattle, WA, USA

**Keywords:** neuronal diversity, discrete cell types, continuous type-specific variability, cell states, mixture model, variational inference

## Abstract

Reproducible definition and identification of cell types is essential to enable investigations into their biological function, and understanding their relevance in the context of development, disease and evolution. Current approaches model variability in data as continuous latent factors, followed by clustering as a separate step, or immediately apply clustering on the data. We show that such approaches can suffer from qualitative mistakes in identifying cell types robustly, particularly when the number of such cell types is in the hundreds or even thousands. Here, we propose an unsupervised method, MMIDAS, which combines a generalized mixture model with a multi-armed deep neural network, to jointly infer the discrete type and continuous type-specific variability. Using four recent datasets of brain cells spanning different technologies, species, and conditions, we demonstrate that MMIDAS can identify reproducible cell types and infer cell type-dependent continuous variability in both uni-modal and multi-modal datasets.

## Introduction

Understanding the nature and extent of cellular diversity in the brain is key to unraveling the complexity of neural circuits, their connectivity and roles in behavior, in health and disease [15, 47, 58]. While fine-grained cell typing helps to uncover stereotypy, the continuous variability around cell type representatives capture the flexibility and uniqueness of neuronal phenotypes and computation, in addition to their spatial organization [16, 50, 23]. However, a lack of methods to identify common aspects of within-cluster variability across cell types and difficulty in reproducible identification of neuronal types in high-dimensional datasets with large number of categories — a setting where clustering algorithms typically struggle — pose challenges towards dissection of the discrete and continuous factors governing the landscape of neuronal phenotypes [9, 43, 15, 58].

In the adult brain, this landscape contains stable wells of attraction, where the well that a cell belongs to represents the discrete and terminal cell type identity. In this analogy, the location within the well describes a continuous source of variability [23, 50, 58]. Among others, disease can perturb the landscape, including by changing the continuous factor around the discrete representative for multiple cell types. Since discrete and continuous factors jointly determine the phenotype, studying them sequentially may infer a qualitatively different characterization of the overall landscape in health and disease. Moreover, individual wells can have complicated shapes as manifested by within-type variability in molecular, physiological, and anatomical features, exacerbating the problem [23, 50, 56, 5]. Statistically, such a collection of discrete wells with non-canonical shapes can be modeled by a generalized mixture model.

Here, we introduce Mixture Model Inference with Discrete-coupled AutoencoderS (MMIDAS), a mixture model-based method designed to jointly infer interpretable discrete and continuous latent factors. We achieve this by utilizing discrete-coupled autoencoders, where the different autoencoding neural networks infer the parameters of their own mixture models from augmented copies of a given sample. We find that encouraging the networks to agree on their categorical assignments via the objective function avoids the notorious mode collapse problem [34] common in generative models involving discrete factors.

In single-cell analysis, dimensionality reduction typically precedes clustering. It is within this simplified representation that methods like community detection algorithms, e.g. Leiden and Louvain, are utilized to identify cell types by constructing graphs [8, 42, 22]. While various dimension reduction methods can infer meaningful continuous representations [45, 11, 37, 10, 53], clustering within these reduced spaces can be suboptimal because dimensionality reduction is not informed of the subsequent clustering objective. Similarly, analysis of within-cluster variability is commonly performed by studying the continuous diversity within each cluster separately, after clustering [50, 56]. Therefore, identifying factors of continuous variability that are common across multiple cell types can be laborious. Moreover, ignoring dependencies between the discrete identity and the remaining continuous variability can produce inaccurate characterizations of both of these factors.

MMIDAS addresses these challenges by jointly performing clustering and dimensionality reduction in an end-to-end differentiable variational formulation. This not only eliminates the need for separate clustering steps, but also enhances the stability of clusters, especially in the brain with its extreme cell type diversity (large number of clusters). Perhaps more importantly, MMIDAS offers a unified and interpretable solution by jointly inferring discrete and continuous variabilities and avoids the need for separate modeling for each cluster. Lastly, MMIDAS is not specialized for a specific data modality. It is naturally applicable to both uni-modal and multi-modal datasets.

MMIDAS is an unsupervised method and does not require any priors on the relative abundances of cell types, labelled data, or batch information [20, 38, 19, 55]. We benchmark MMIDAS on a diverse selection of datasets including two uni-modal single-cell RNA sequencing (scRNA-seq) datasets profiling multiple regions of the mouse cortex obtained by different platforms [51, 56], a multi-modal dataset with both electrophysiological and transcriptomic profiles of cortical neurons in mice [21, 49], and a recent scRNA-seq dataset from the human middle temporal gyrus (MTG) exploring changes in gene expression in disease, the Seattle Alzheimer’s Disease Brain Cell Atlas (SEA-AD) [17]. We demonstrate that the mixture models learned by MMIDAS discover accurate and interpretable discrete and continuous factors of cellular diversity. Furthermore, we show that MMIDAS offers novel insights into the molecular mechanisms governing the cellular landscape. By identifying features, e.g. genes, that control the continuous latent factors via traversal analyses, MMIDAS infers biological processes such as cell metabolism, physiological response characteristics, spatial gene expression gradients, and disease stages that are encoded by those continuous factors.

## Results

### A. Structured representation learning for cellular phenotypes

Consider (**c, s**) = ℰ (**x**), where **x** denotes the measurement vector for a cell. Here, **c** denotes a categorical variable, **s** captures the remaining continuous variability around the cluster representative, which depends on **c**, and E denotes a mapping function. Let 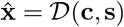 be an approximate reconstruction of the true measurement **x** via a mapping function 𝒟. In modern machine learning, this is known as an autoencoding architecture: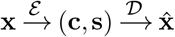, and ℰ and 𝒟 are referred to as the encoder and the decoder, respectively, whose parameters can be optimized by minimizing the discrepancy between **x** and 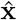.

Learning such interpretable dissections of variability, however, has been challenging because the objective of minimizing the discrepancy is not informative enough to guide the optimization to a meaningful parametrization in a highly non-convex landscape. This is known as the mode collapse problem [34]. While methods have been developed to circumvent this issue, they require prior knowledge of the underlying categorical abundances, commonly assumed to be uniform [12, 29] (Methods). However, this assumption is violated in single-cell datasets because cell types can have very different abundances. We reason that multiple autoencoders with coupled discrete variables can inform and regularize each other, removing the need for prior knowledge.

MMIDAS is a collective learning framework that uses multiple variational autoencoding neural networks (VAE) to jointly infer interpretable and accurate categorical and continuous factors of variability in the presence of a high dimensional discrete space (Fig. 1a). MMIDAS consists of a set of independent and architecturally identical VAEs. Each VAE “arm” parameterizes cell type identity (discrete diversity) and continuous type-dependent variability (continuous diversity) (Fig. 1b), and their parameters are tuned by efficient and scalable stochastic gradient descent.

**Figure 1.**
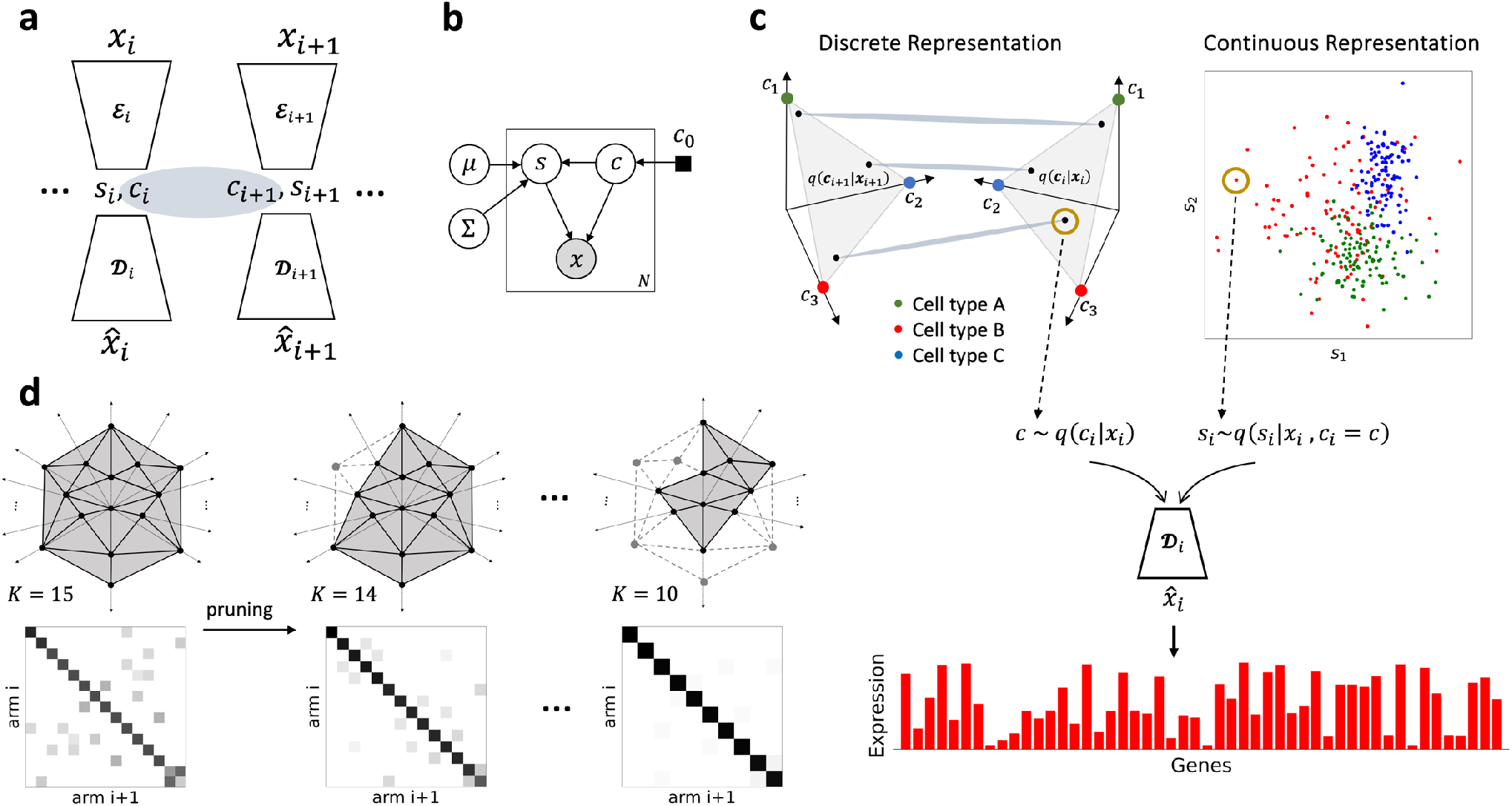
MMIDAS, a multi-arm autoencoder framework for mixture model inference. (a) Schematic of MMIDAS illustrating a pair of discrete-coupled VAE arms. Each arm comprises encoding (ℰ) and decoding (𝒟) functions that transform high-dimensional data (**x**) into a mixture latent representation and reconstruct it 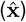. Individual arms receive non-identical noisy copies of a given sample **x**, i.e., {…, **x**_*i*_, **x**_*i*+1_,…}, belonging to a neighborhood of **x**, to learn mixture representations, i.e., {…, *q*(**c**_*i*_, **s**_*i*_), *q*(**c**_*i*+1_, **s**_*i*+1_), …}. VAE arms cooperate to learn categorical abundances, *p*(**c**). (b) Graphical model of each VAE, depicting categorical variable (type), **c**, and type dependence of the continuous variable, **s**. The continuous variable **s** is parameterized by a Gaussian distribution with trainable mean (***µ***) and covariance (**Σ**). **c**_0_ denotes the prior distribution of the categorical variable, provided by a different VAE arm in a self-supervised manner. (c) Discrete and continuous representations of cellular heterogeneity in the latent space. **c** is a categorical random variable that follows a categorical distribution, *q*(**c**_*i*_|**x**_*i*_), assigning probabilities to each cell within a *K*-simplex, with *K* possible categories. MMIDAS regularizes the categorical distribution, *q*(**c**_*i*_|**x**_*i*_), via a consensus constraint in the simplex. The continuous type-dependent variability of the cell is obtained by mapping the profile to a lower-dimensional continuous space, given the categorical identity. Accurate reconstruction is achieved by jointly decoding the contributions of **c** and **s**, e.g. transcriptomic profile (bar plot for a red cell) through the decoder of the *i*-th arm, *D*_*i*_. (d) Schematic of the pruning process in an over-parameterized simplex. During training, the model calculates the consensus measure between pairs of arms for each vertex. During pruning, vertices with the lowest consensus measures are pruned. For the sake of visualization, simplices are plotted in a 2-dimensional space.

In MMIDAS, arms receive non-identical observations, e.g. **x**_*i*_ and **x**_*i*+1_, that share the categorical variable (Fig. 1a). To enable this for unimodal datasets in an entirely unsupervised manner, we use “type-preserving” data augmentation (Methods and Supplementary Note 6, Fig. S21a). While each arm has its own mixture representation, all arms cooperate to learn a consensus category, **c**, for a given cell, representing the discrete diversity encoding the cell type. **c** is a random variable that governs the discrete probabilistic representation of each cell within a simplex whose dimensionality, *K*, denotes the expected number of cell types (Fig. 1c). The within-type deviation of the cell’s phenotype around the cell type representative is captured by the continuous latent factor, **s** (Fig. 1c). While the true number of types and their relative abundances are unknown, MMIDAS utilizes a pruning technique on an over-parameterized simplex to automatically identify the number of cell types in the dataset (Fig. 1d). The underlying principle is that it is harder to reach a consensus over spurious types (Methods). The use of a common probabilistic model, i.e. *p*(**s**|**c, x**) and *p*(**x**|**c, s**), as shown in Fig. 1, encourages consistent encoding of continuous variability **s**|**c** in the continuous space, across types.

We use the Gumbel-Softmax distribution [28], a continuous relaxation of the categorical distribution, which allows for a differentiable approximation of sampling from a categorical distribution in the simplex (Methods). The consensus constraint is defined based on a distance metric according to Aitchison geometry [1], among differentiable samples in the simplex (Eq. 9), which avoids the mode collapse problem (Methods and Supplementary Note 4). Theoretical analysis (Methods and Supplementary Note 3) and experimental results that we present in the next section show that this approach enhances accuracy, robustness, and interpretability of the inferred factors without requiring prior information on the relative abundances of the categories. Details of the network architecture and training configurations can be found in Figs. S21b,c, Table S7, and Supplementary Note 14. Throughout this manuscript, “discrete” and “categorical” are used interchangeably. The categorical variable refers to **c**, that follows a categorical distribution, *q*(**c**|**x**), in the simplex. For a given cell sample, **x**_*m*_, the discrete cell type or category, should be interpreted as 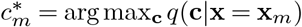.

Before demonstrating MMIDAS on single-cell datasets, we first study a benchmark machine learning dataset, the MNIST handwritten digits dataset (Supplementary Note 5 and Supplementary Note 7). Although MNIST is not as complex as single-cell datasets, we have included it in our evaluation to enable interpretability and facilitate the assessment of MMIDAS results with known ground truth. Our results indicate that MMIDAS outperforms earlier mixture model-based methods, and its computational cost remains comparable to that of the baselines (Fig. S1, Table S1, and Table S2). Supplementary Note 8 and Fig. S5 demonstrate that MMIDAS’s performance is minimally affected by variations in the consensus (coupling) factor. Thus, we do not fine-tune this hyperparameter in any of our studies.

### B. scRNA-seq analysis of mouse cortical regions

When analyzing uni-modal scRNA-seq datasets, each VAE arm receives a noisy copy of a cell’s profile, on which they try to infer a shared discrete factor through a variational optimization (Methods). In the unsupervised setting, the noisy copies are generated by “type-preserving” augmentation, which generates independent and identically distributed copies of data within a small neighborhood around the original sample, thus remaining in the well of attraction and preserving the cell’s type identity with high probability (Methods, Supplementary Note 6, Fig. S3b,c, and Fig. S4).

#### Transcriptomic cell type identification

We first study cellular diversity in a Smart-seq dataset of the mouse primary visual (VISp) and anterior lateral motor (ALM) cortices whose reference taxonomy includes 115 transcriptomic types (t-types) [51] (Fig. 2 and Fig. S14). These reference t-type clusters were obtained by the “scrattch.hicat” method [51, 56], which utilizes dimensionality reduction and graph-based clustering. Briefly, it iteratively performs principal component analysis (PCA) followed by Louvain graph-based clustering with clustering criteria specifically designed for scRNA-seq data analysis [51, 56].

**Figure 2.**
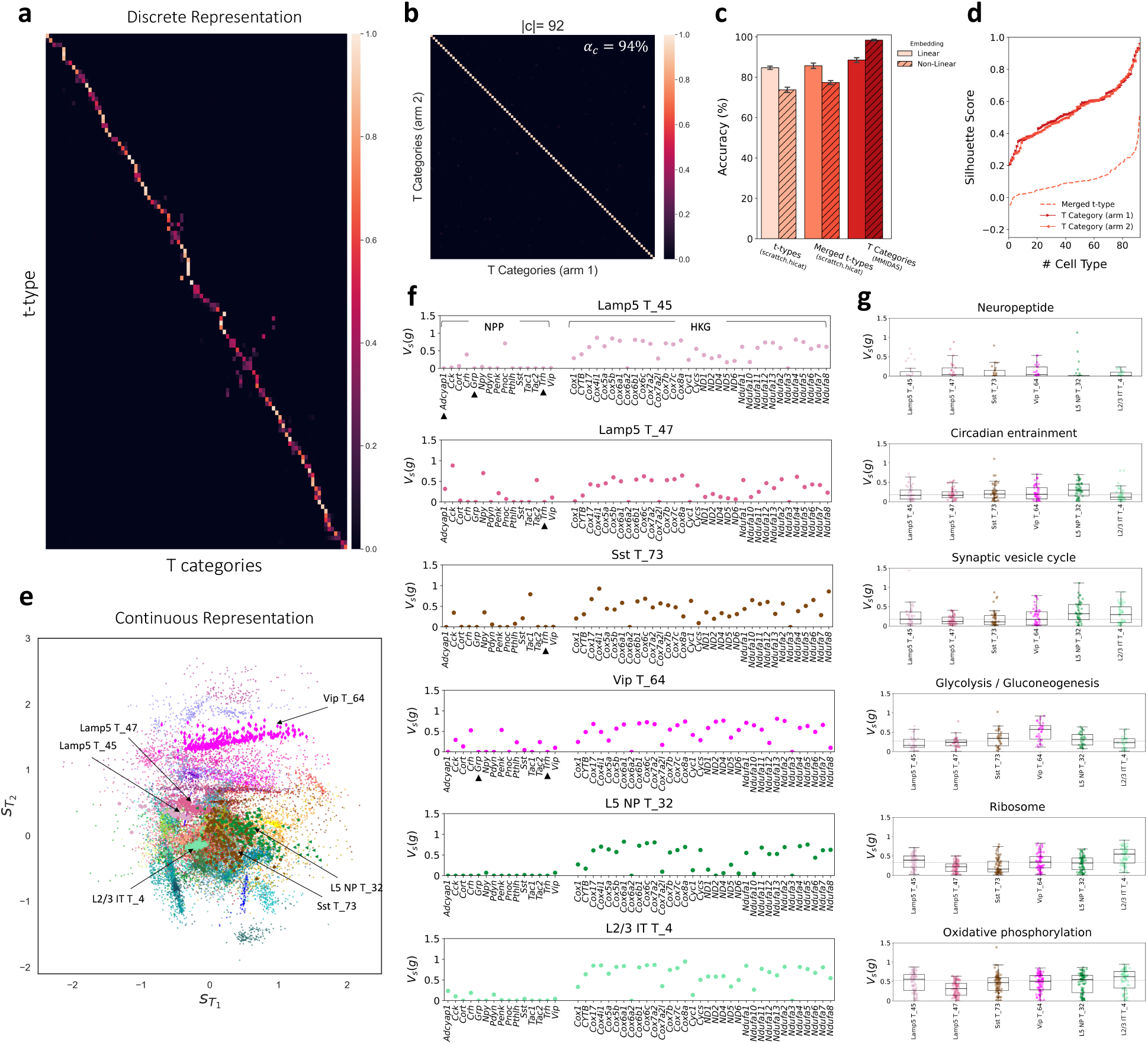
Characterizing discrete and continuous diversity in the adult mouse VISp and ALM neurons sequenced with the Smart-seq platform. (a) Mutual information scores between 115 t-types identified by scrattch.hicat [51] (y-axis) and 92 consensus categories inferred by MMIDAS (x-axis). (b) Confusion matrix between discrete representations of a pair of VAE arms. Consensus score, *α*_*c*_ (Methods), is 94%. (c) Mean balanced classification accuracy with standard deviation, obtained through 10-fold cross-validation, evaluated across two embeddings: linear (100-dimensional) and non-linear (10-dimensional). The analysis includes three groups of class labels: t-types denote the 115 reference types, merged t-types are t-type labels merged to 92 classes according to the reference hierarchy [51], MMIDAS categories are the 92 categories inferred by MMIDAS. (d) Average Silhouette scores per-cluster. Merged t-types use the linear embedding and MMIDAS T categories use the non-linear embedding. (e) 2-dimensional representation of within-cluster continuous diversity, color coded based on transcriptomic types inferred by MMIDAS, highlighting the distributions for 6 example T categories. (f) Variation in gene expression, *V*_*s*_(*g*) (Methods), based on the inferred continuous variable. For a given transcriptomic type, we alter the continuous variable **s** by traversing along the dominant principal component. Each data point illustrates the resulting variation for a subset of neuropeptide precursor (NPP) genes associated with neuropeptide signaling, and a subset of housekeeping genes (HKG) contributing to oxidative phosphorylation. Genes with zero expression are marked with▴. (g) Variation in gene expression across a subset of transcriptomic types for pathway-associated gene modules. Each subfigure illustrates box plots of overall variation in gene subsets contributing to five distinct pathways: Neuropeptide signaling (15 NPP genes and 10 NP-GPCR genes), circadian entrainment (50 genes), synaptic vesicle cycling (SynapCyc, 49 genes), Glycolysis / Gluconeogenesis (31 genes), Ribosome function (73 genes), and oxidative phosphorylation (OxPhos, 88 genes). The horizontal line within each box marks the median value and the dashed line represents the overall median across the indicated cell types. Subfigures are arranged from the lowest to the highest variation. Pathway-associated gene modules do not overlap, except for SynapCyc and OxPhos, which share 15 genes.

For this dataset, MMIDAS with a pair of VAE arms (Supplementary Note 11), uncovers 92 distinct transcriptomic categories (Fig. 2a). At the time of inference on a test set, the two VAEs agree with each other on 94% of their categorical assignments, which we define as the consensus score (Fig. 2b, Methods). These results are obtained by optimizing an initial, over-parameterized 120-simplex and a 2-dimensional continuous space (Supplementary Note 10). Mutual information analysis shows strong alignment between categories inferred by MMIDAS and the reference t-types (Fig. 2a). The MMIDAS categories, however, uncover a more reproducible and separable discrete representation of the underlying data as quantified by the classification performance on 10-fold cross-validation (Fig. 2c) and the average Silhouette score (Fig. 2d), which takes values between − 1 and 1, where 1 denotes perfect clusterability. We establish these results using both linear and nonlinear embeddings of the data (Methods). We also observe that scrattch.hicat’s performance degrades when the nonlinear embedding is used, reflecting its application of PCA to reduce dimensionality. To isolate the improvement due to MMIDAS, Fig. 2c and Fig. 2d also show the results due to merging of the t-types according to the reference hierarchy until 92 clusters are obtained.

#### Inferring genes contributing to continuous diversity

Having demonstrated that the reproducibility of the clusters inferred by MMIDAS outperforms that by sequential application of dimensionality reduction and clustering, we turn our attention to a unique advantage of MMIDAS. While the discrete representation uncovers more stable transcriptomic categories, the latent representation of continuous type-specific variability captures within-type diversity. Fig. 2e demonstrates the continuous representation, **s**, a 2-dimensional latent space of all cells in the Smart-seq dataset, highlighting the type-dependent variability for a few example types.

The fundamental difference between **c** and **s** is their discrete vs continuous nature. In our formulation, **s** depends on **c** to capture the continuous variability around the cluster representative. When such within-cluster variability is itself cell-type-dependent, e.g. some cell types displaying stronger spatial gradients than others, **s** can have (weak) information on the cell type (similar patterns in the MNIST dataset in Fig. S2). We quantified this by training a classifier on **s**: within-type variability predicts the inferred categories with an accuracy of 33%. While this is significantly lower than the classification ability of **c** (Table 1), it is also above the chance level (Fig. S15), which is 5% for this dataset.

**Table 1.**
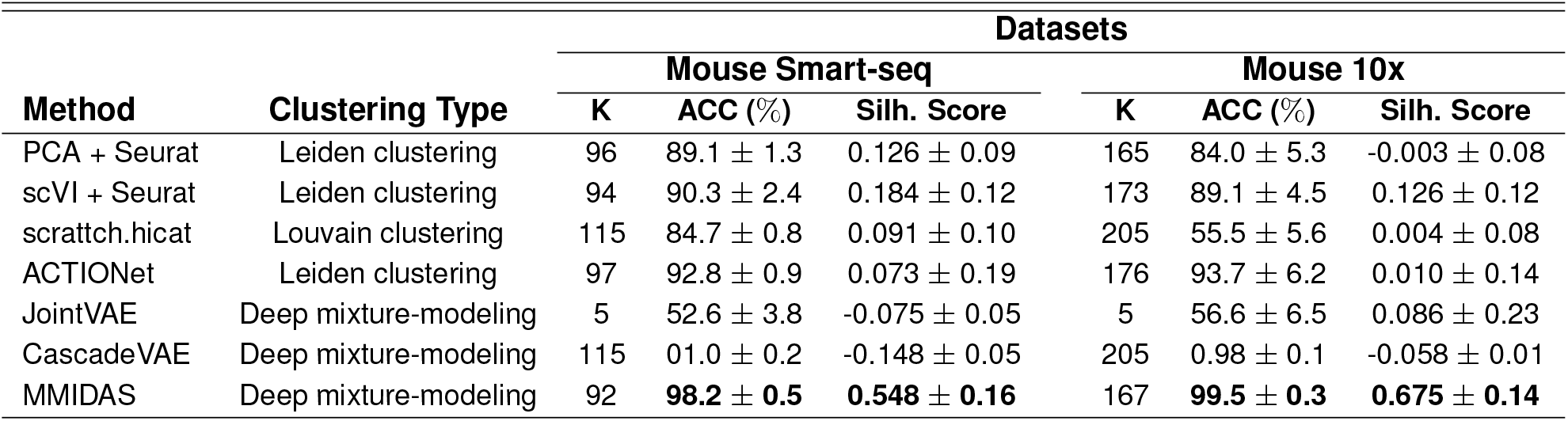
Evaluating scRNA-seq data clustering performance for three established scRNA-seq community detection-based algorithms, two state-of-the-art unsupervised deep mixture modeling approaches, and the proposed method, MMIDAS. *K* denotes the number of identified clusters (cell types) for each method. To quantify the stability of identified cell types, we present the balanced accuracy (ACC), determined through Random Forest classification on test samples in 10-fold cross-validation. We also report the mean and standard deviation of the Silhouette Scores (Silh. score) across all identified clusters. MMIDAS significantly outperforms in both metrics and across two datasets.

The continuous representation can help to characterize the contribution of each gene to both the discrete and continuous heterogeneity. To study correlates of the continuous representation with gene expression, we introduce a traversal analysis. We hold the categorical variable at a constant value (thereby fix the cell type), move along the span of the continuous representation, and evaluate the change in reconstructed expression values for each gene (Methods). To quantify this change and enable comparisons across cell types, we introduce *V*_*s*_(*g*), the normalized change in the expression of gene *g* triggered by traversing the continuous variable **s** along the primary principal component of the continuous representations of cells in the given type (Methods).

Fig. 2f and Fig. 2g showcase the results of this traversal analysis for six MMIDAS T categories within the *Lamp5, Sst, Vip*, L5 NP, and L2/3 IT subclasses. In Fig. 2f, the x-axis lists two gene groups: the first comprises 15 neuropeptide precursor (NPP) genes contributing to neuropeptide signaling pathways, and the second consists of a set of 35 housekeeping genes (HKGs) primarily involved in cellular oxygen consumption within the oxidative phosphorylation pathway. We observe that NPP genes, which exhibit highly specific expression across different neuronal types [48], display lower variability under the continuous variable traversal. In contrast, HKGs, integral to fundamental cellular maintenance functions [31], exhibit greater variation (Fig. 2f). Consistent with previous studies, this implies that NPP genes play a more substantial role in the discrete heterogeneity rather than the continuous one. HKGs from the *Cox, Ndufa*, and *ND* families, which are related to the electron transport chain and oxidative phosphorylation, are far more variable within cell types, suggesting there may be modules of genes related to a notion of cellular state that the continuous variable captures. Our results also suggest that the expression of activity-regulated genes (e.g. *Cox*) is cell type-dependent so that these genes respond differently to neuronal activation based on the underlying cell type (compare sub-figures in Fig. 2f).

To investigate this further, we selected gene modules implicated in a broad range of pathways [32], that *a priori* are rather generic cellular processes not expected to govern cellular identity (Fig. 2g). We observe that different pathways drive continuous variability across types. For example, variability in synaptic vesicle cycle pathway genes is the most in cells of type L5 NP T_32, whereas that in glycolysis/gluconeogenesis pathway genes is much higher in cell type Vip T_46 (among the types shown in Fig. 2g). A more comprehensive picture is provided in the Figs. S9, S10, and S11. The state variables inferred by MMIDAS can thus be viewed as a handle to explore and identify cell type specific gene modules that may be driving variability within populations. Further biological insights into the inferred discrete and continuous representations are discussed in Supplementary Note 13 and Fig. S8.

We next analyze the expression landscape of the mouse cortex as profiled by the 10x platform, where the reference taxonomy reports 113 glutamatergic and 97 GABAergic t-types [56] (Fig. 3, Figs. S16 and S17). In contrast, MMIDAS infers 97 glutamatergic and 70 GABAergic categories (Fig. 3a,b). As before, Fig. 3c,d,e and Fig. S18 quantify that MMIDAS offers a significantly more reproducible and identifiable characterization of the discrete neuronal diversity compared to the reference taxonomy. Indeed, the superiority of MMIDAS is even more evident in analyzing data obtained by the 10x platform, whose throughput, noise level, and gene dropout rate all tend to be higher [54].

**Figure 3.**
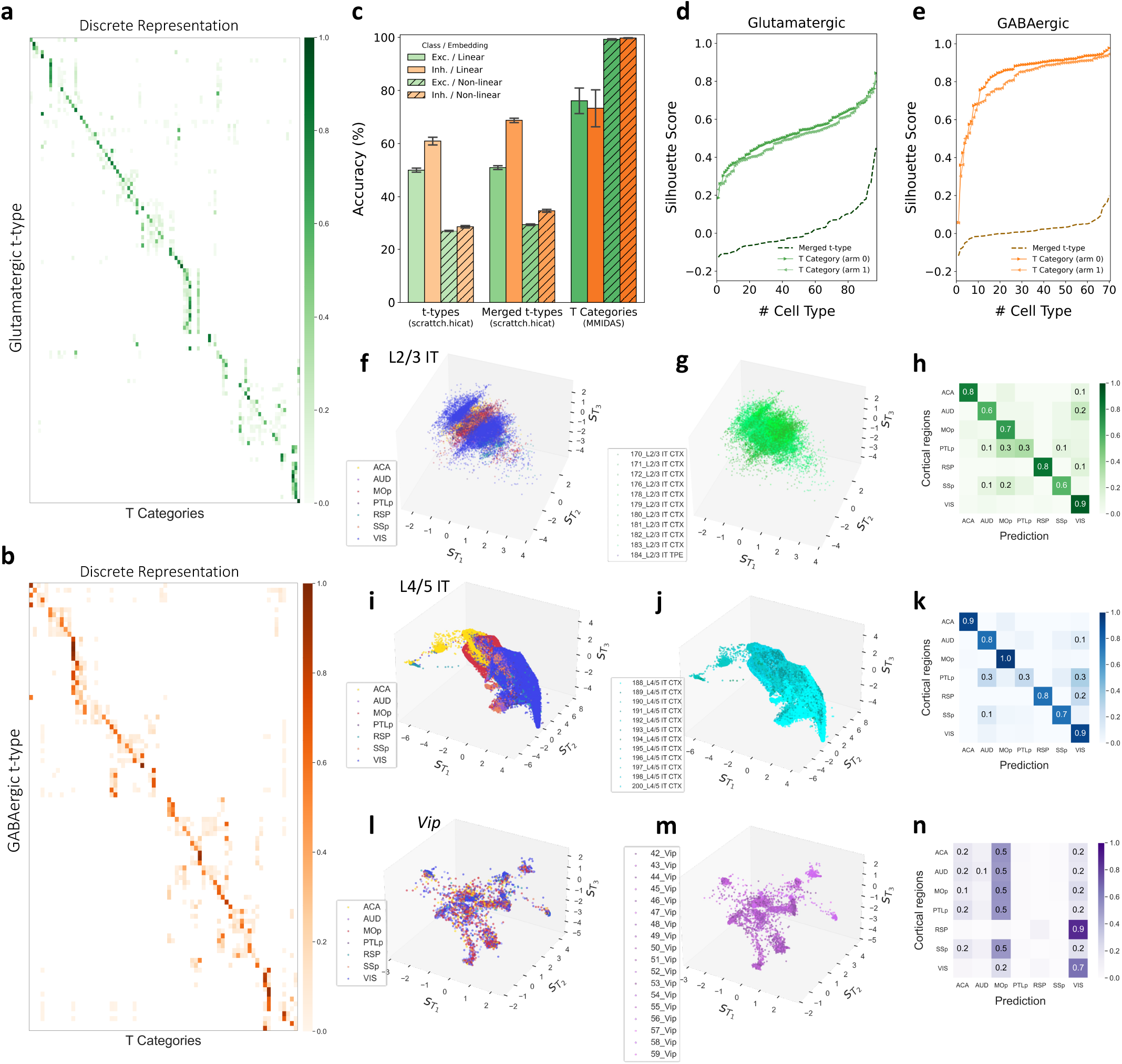
Characterizing discrete and continuous diversity in the adult mouse ACA, AUD, MOp, PTLp, RSP, SSp, VIS neurons sequenced with the 10xv2 platform. (a) Mutual information scores between 113 excitatory (glutamatergic) t-types identified by scrattch.hicat [56] (y-axis) and 97 consensus categories inferred by MMIDAS (x-axis). (b) Mutual information scores between 92 inhibitory (GABAergic) t-types [56](y-axis) and 70 consensus categories inferred by MMIDAS (x-axis). (c) Balanced classification accuracy based on 10-fold cross-validation (mean and standard deviation), evaluated across a 100-dimensional linear and a 10-dimensional non-linear embeddings for three groups of class labels: t-types in the reference taxonomy, merged t-types, and the transcriptomic categories inferred by MMIDAS. (d,e) Average Silhouette scores per-cluster, for cells in glutamatergic and GABAergic classes. Merged t-types use the linear embedding and MMIDAS T categories use the non-linear embedding. (f) Latent representation of the continuous diversity for L2/3 IT cells, color coded by cortical regions, compared to (g), which is color-coded by their t-types. (h) Confusion matrix for predicting cortical regions from the inferred continuous representation of L2/3 IT neurons. Classification accuracy: 68%. (i) Continuous latent representation of L4/5 IT cells, color coded by cortical regions and (j) color coded by their t-types. (k) Confusion matrix for predicting cortical regions, with accuracy 77%, using the inferred continuous representation of L4/5 IT neurons. (l,m,n) Same as f,g,h, respectively, but for *Vip* cells, showing that the latent representation encodes neither spatial gradients (accuracy 22%) nor cell type information (accuracy 32%, Fig. S19c) for *Vip* cells.

While MMIDAS is designed to infer a composite landscape of continuous and discrete cellular diversity, and not as a clustering method *per se*, we inspect the clusterability of the identified cell types in a comparative manner to elucidate MMIDAS’s stability and precision in the inference of categorical variables (cell types). Our investigation involves benchmarking against well-established scRNA-seq clustering methods, Seurat [22], scrattch.hicat [51, 56] and ACTIONet [42]. Additionally, we compare MMIDAS to state-of-the-art unsupervised VAE-based mixture models, namely JointVAE [12] and CascadeVAE [29]. Table 1 collates the performances of six competing models and MMIDAS in the clustering task. Among community detection-based algorithms, scrattch.hicat produces a larger number of cell types while Seurat and ACTIONet, utilizing the Leiden algorithm for clustering, capture a comparable number as MMIDAS. However, their Silhouette and accuracy scores are significantly lower than those of MMIDAS, suggesting poor clusterability and casting doubt on the discrete nature of the clusters identified by Seurat, ACTIONet, and scrattch.hicat. We observe that constructing the neighborhood graph from a linear embedding further degrades their performance (Fig. S6c,d). Among the deep mixture models, both JointVAE and CascadeVAE struggle to identify meaningful cell types, demonstrating mode collapse within the high-dimensional discrete space (Fig. S6a) and inferring an almost uniform distribution across types despite substantial differences in their relative abundances (Fig. S6b). Importantly, MMIDAS explicitly addresses both of those issues, producing high-quality cell type definitions as well as dissecting the continuous, type-dependent variability. The training details for models outlined in Table 1 and further results are reported in Supplementary Note 9 and Table S6.

#### Spatial gradient analysis of neuron types across regions of the isocortex

We next study whether the continuous type-dependent factor can inform about spatial organization. Fig. 3f and Fig. 3i demonstrate such encoding of a contiguous set of cortical regions – anterior cingulate area (ACA), auditory cortex (AUD), primary motor cortex (MOp), posterior parietal associative area (PTLp), retrosplenial cortex (RSP), primary somatosensory area (SSp), and visual cortex (VIS) – by the continuous latent factor, **s**, for L2/3 IT and L4/5 IT neurons in the mouse 10X dataset, consistent with previous literature [56]. The power of the continuous representation in predicting spatial location, compared to cell type information (Fig. 3g, Fig. 3j, and Fig. S19a,b), is more evident in Fig. 3h and Fig. 3k. These figures quantify the separability of cortical areas based on inferred **s** ∼ *q*(**s** | **c, x**). The PTLp region, being relatively small with fewer cells in this dataset, may contribute to its confusion with other regions.

This finding suggests the existence of more pronounced spatial gene expression gradients in the mouse cortex than previously established (refer to Fig. 7C in the study conducted by Yao et al., 2021 [56]). Across numerous regions, including MOp, RSP, and SSp, MMIDAS consistently demonstrates more precise encoding of the spatial gradient in the continuous factor. Additionally, we contrast these observations from Glutamatergic subpopulations with those from a GABAergic subclass, the *Vip* cells, for which MMIDAS does not identify such spatial gene gradients across the cortical sheet (Fig. 3l,n). Thus, the continuous type-dependent latent factor inferred by MMIDAS can effectively encode spatial gradients of gene expression, when they exist.

### C. Multimodal single-cell analysis

Simultaneous capture of a diverse range of cellular features from individual cells is key to attaining a definitive characterization of neuronal diversity [47, 6, 18, 21, 35, 46, 19, 58, 49]. A primary challenge in analyzing multimodal single-cell datasets is disentangling factors of variability that are shared among modalities, e.g. cell types, from those that are unique to specific modalities, without access to (and the potential bias of) labeled data. MMIDAS resolves this issue by enabling joint inference of both (shared) discrete and (modality-specific) continuous variabilities. In the context of multimodal data analysis, individual VAE arms learn mixture representations for their respective input modalities, i.e. one VAE arm per modality, while being constrained to learn similar discrete representations across modalities (Fig. 4a).

**Figure 4.**
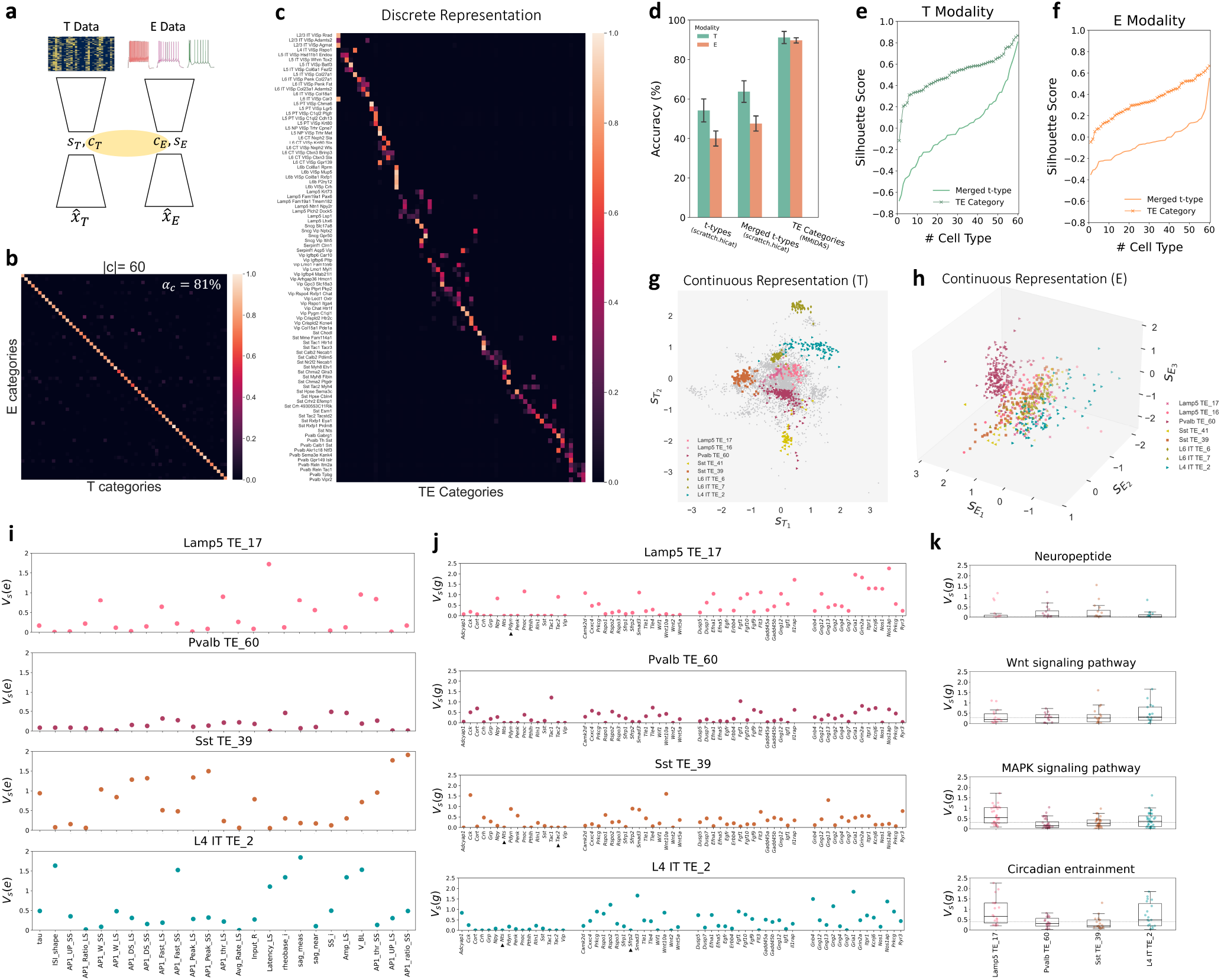
Multimodal characterization of discrete and continuous diversity in adult mouse VISp neurons. (a) Schematic of MMIDAS showing the discrete-coupled VAE architecture in the presence of two data modalities, i.e. transcriptomics (T) and electrophysiology (E), profiled with Patch-seq [21]. The consensus constraint encourages the T and E arms to achieve identical discrete representations, **c**_*T*_ and **c**_*E*_, while independently learning their own latent continuous factors, **s**_*T*_ and **s**_*E*_. (b) Consensus matrix between discrete representations of a pair of VAE-arms, at the time of inference with a consensus score (*α*_*c*_) of 81%. (c) Mutual information scores between the 92 reference t-types [21] (y-axis) and 60 transcriptomic-electrophysiology (TE) categories inferred by MMIDAS (x-axis). (d) Balanced classification accuracy, determined through 10-fold cross-validation, is presented with mean and standard deviation values. This analysis utilizes a 30-dimensional nonlinear embedding and categorizes data into three distinct groups: 92 t-types, 60 merged t-types according to the hierarchy in [51], and 60 TE categories inferred by MMIDAS. (e, f) Average per-cluster Silhouette scores based on transcriptomic and electrophysiology profiles for merged t-types and MMIDAS categories, respectively. (g,h) Latent continuous representations of within-cluster diversity for transcriptomic and electrophysiology modalities, outlining the continuous diversities of eight MMIDAS TE categories. (i,j) Continuous traversal analysis of electrophysiological features. For a given transcriptomic type, the continuous variable **s** is changed by traversing along the primary principal component. (i) Total variation of **s**_*E*_ for 25 IPFX features in the patch-seq dataset. (j) Gene expression variation for 16 NPP genes associated with neuropeptide signaling, as well as 45 genes suggested as contributors to Wnt signaling, MAPK signaling, and circadian entrainment pathways. (k) Gene expression variations across transcriptomic types for pathway-associated gene modules. The box plots display total variation of gene modules contributing to different pathways across four TE types. Here, we have neuropeptide signaling involving a set of 35 NPP and NP-GPCR genes, Wnt signaling with 17 genes, MAPK signaling with 36 genes, and circadian entrainment with 19 genes. The dashed line displays the overall median of gene expression variation across the indicated TE types. The gene module involved in the neuropeptide pathway exhibit the least variation, while one contributing to circadian entrainment display the highest variation.

We study transcriptomic and electrophysiological diversity in a large dataset of mouse visual cortical neurons [21, 49] (Fig. 4). These two observation modalities were profiled by the Patch-seq technique [6, 7, 21], which combines scRNA-seq (T data) with whole-cell patch clamp electrophysiology (E data). In this dataset, MMIDAS uncovers 60 shared TE categories (Fig. 4b,c) with a consensus score of 81% (Fig. 4b) between the T and E modalities. As in the previous datasets, the classification and Silhouette analyses for each modality indicate that consensus TE categories inferred by MMIDAS offer a more accurate and reproducible discrete representation than the t-type or the merged t-type labels (Fig. 4d,e,f). Noting that the reference transcriptomic taxonomy suggested 97 t-types (and MMIDAS inferred 92 categories in the analysis of the VISp Smart-seq dataset), we observe that the presence of another data modality, E data, influences cell type definitions and leads to fewer cell types [18, 21, 19, 49]. On the other hand, the pronounced alignment between TE categories and t-type labels(Fig. 4c) suggests that a diagonally-dominant, square-like confusion matrix is potentially attainable between transcriptomic and morpho-electric characterizations.

Fig. 4g and Fig. 4h illustrate the continuous representations corresponding to a subset of categories for each data modality. We examine the relationship between the features of each modality and their corresponding continuous representations through the continuous traversal analysis. Fig. 4i,j,k showcase the findings of this analysis for four MMIDAS TE categories within the *Lamp5, Pvalb, Sst*, and L4 IT subclasses. Each individual panel in Fig. 4i illustrates the variation of electrophysiological features resulting from traversing **s**_*E*_ along the primary principal axis (associated with the category) within the continuous representation of the E modality. Here, the x-axis corresponds to a set of 25 intrinsic physiological features (IPFX features – see the Datasets section in Methods). We observe that, unlike most other features, the average firing rate (Avg_Rate_LS) does not seem to be modulated within individual categories. Moreover, the intrinsic electrophysiology of some categories seem to be highly stereotyped (Pvalb TE_60), while others display much higher within-type variability (Sst TE_39, L4 IT TE_2).

Fig. 4j and Fig. 4k present the outcomes of continuous traversal (for the same TE types) obtained by traversing **s**_*T*_ along the principal axis for the T modality. In Fig. 4j, the x-axis shows four gene sets: 16 NPP genes contributing to neuropeptide signaling, 15 genes linked to Wnt signaling, 15 genes related to MAPK signaling, and 15 genes involved in circadian entrainment. Among their many functions, the Wnt- and MAPK signaling pathways are thought to regulate neuronal homeostasis and synaptic plasticity [26, 52]. Consistent with our earlier findings (Fig. 2f,g), NPP genes display lower variability under changes in the continuous latent factor relative to the remaining gene subsets (Fig. 4j). We further refine our focus on pathways comprising a minimum of 15 genes in this dataset (Fig. 4k). As in the mouse Smart-seq dataset, the level of variability is a function of both the pathways and the TE types. For instance, the variation in the gene set related to Wnt signaling is most pronounced in cell type L4 IT TE_2, whereas the variation in genes associated with MAPK signaling is notably higher in cell type Lamp5 TE_17 (among the considered types).

### D. Cellular heterogeneity in Alzheimer’s disease

Many neurological diseases, such as Alzheimer’s disease (AD), display progressive pathology and clinical symptoms. Such progression impacts cells differentially [40, 17], although imperfect understanding of phenotypic changes in cell types as a function of disease progression presents a challenge. As a further complication, disease progress may include changes to the relative abundance of cell types as well as within-type continuous transitions over the gene expression and pathology landscapes. We address these challenges by exploring the recent Seattle Alzheimer’s Disease Brain Cell Atlas (SEA-AD) dataset [17], which includes scRNA-seq profiles from the middle temporal gyrus (MTG) of 84 donors spanning the spectrum of AD pathology (Methods).

We train a MMIDAS model with two arms on three subclasses of neurons in this dataset; the IT, *Sst*, and *Pvalb* neurons. The original SEA-AD study [17] describes 42 transcriptomic categories for this set (13 IT, 16 *Sst*, 13 *Pvalb*), referred to as *“supertypes”* and obtained by mapping transcriptomic data to the BICCN neurotypical reference data[30] and annotating cells via scANVI [55]. In contrast, MMIDAS infers 30 categories (14 IT, 9 *Sst*, 7 *Pvalb*), explaining GABAergic variability with a smaller number of discrete types. Instead, the continuous variability around those (fewer) discrete types contributes to the observed transcriptomic diversity. Beyond this difference, the mutual information scores (Fig. 5a,b) suggest a reasonable correspondence between the inferred categories and the SEA-AD supertypes. We also observe more confusion between the categorical assignments of L2/3 IT neurons and some *Sst* neurons. Similar to our previous results, the classification and Silhouette analyses of transcriptomic types indicate that consensus T categories inferred by MMIDAS are more accurate and reproducible than the suptertypes or the merged supertypes (Fig. 4c,d).

**Figure 5.**
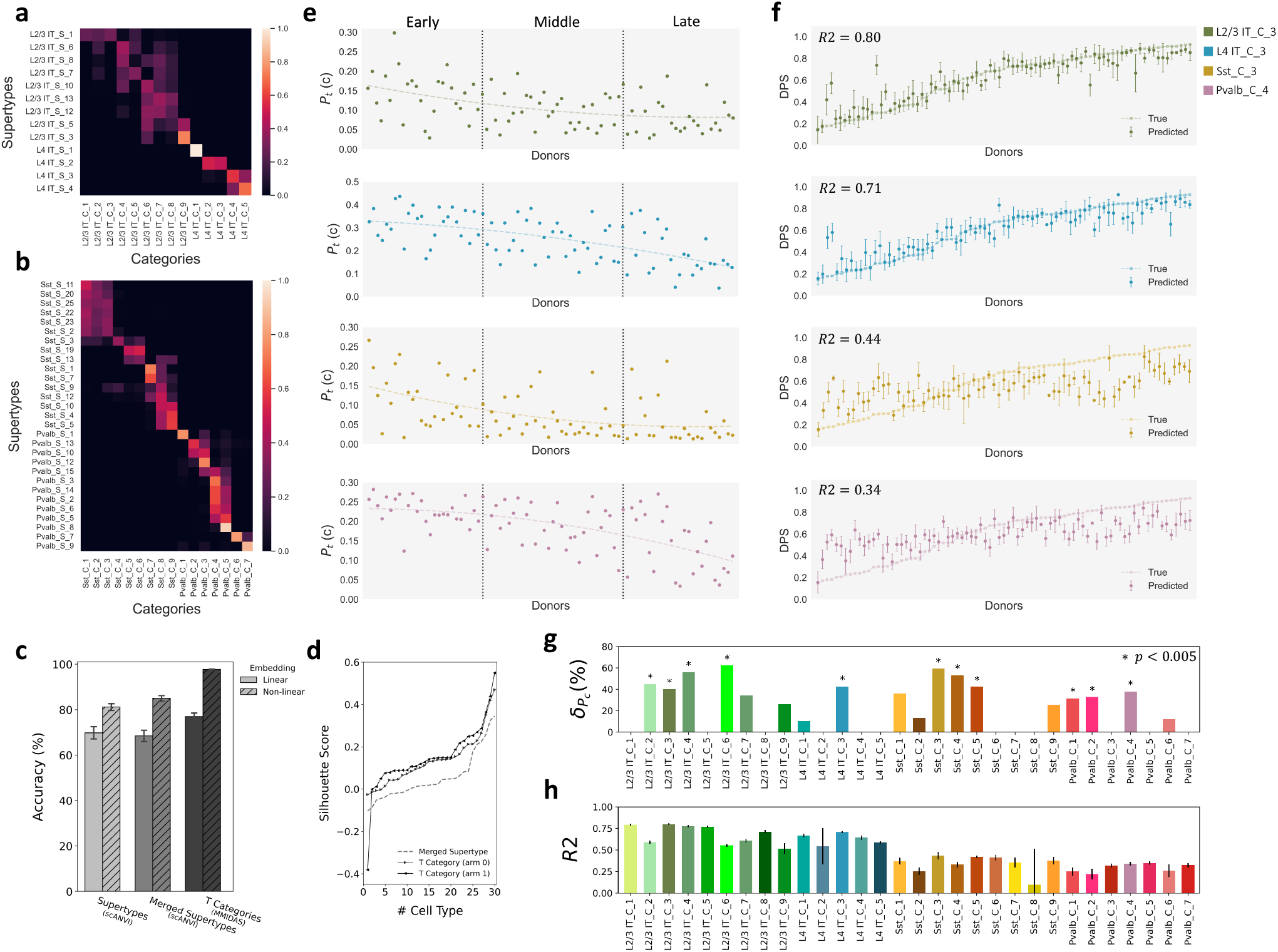
Characterizing discrete and continuous diversity in human MTG scRNA-seq data in Alzheimer’s disease. (a) Mutual information scores between 13 IT supertypes obtained with scANVI [55, 17] (y-axis) and 14 consensus categories inferred by MMIDAS (x-axis). (b) Mutual information scores between 29 *Sst* and *Pvalb* supertypes [17] (y-axis) and 16 consensus categories inferred by MMIDAS (x-axis). (c) Mean balanced classification accuracy with standard deviation, obtained through 10-fold cross-validation, evaluated across linear (100-dimensional) and non-linear (30-dimensional) embeddings. The analysis includes three groups of class labels: supertypes denote the 42 reference types, merged supertypes are supertype labels merged to 30 classes according to the reference hierarchy [17], MMIDAS categories are the 30 categories inferred by MMIDAS. (d) Average Silhouette scores per-cluster. Comparison between merged supertypes and MMIDAS categories, both employing the non-linear embedding. (e) Predicting relative abundance of cell types inferred by MMIDAS (*P*_*t*_(**c**) for category **c** at (pseudo) time *t*) from the pseudo-progression score of donors. Vertical lines separate early, middle, late stages of disease. Abundances of L2/3 IT, *Sst*, and *Pvalb* neurons decrease in the early stage while that of L4 IT neurons decrease in the late stage. Donors ordered by their pseudo-progression score obtained from MTG neuropathology [17] in the x-axis. (f) Predicting disease progression pseudotime (DPS) [17] from the continuous representation inferred by MMIDAS. Dots/bars denote mean/standard deviation across 10-fold cross validation. IT cells, but not *Sst* and *Pvalb* cells, have high coefficient of determination, *R*2, which provides a measure of how well the continuous representation can explain the variance in the pseudo disease progression. x-axis same as in (e). (g) Rate of cellular loss in AD. 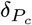 denotes the proportion of changes of *P*_*t*_(**c**), between early and late stages of the disease (Methods). About 35% of the transcriptomic types exhibit significant reduction as they progress from the early to the late stage of AD. *p*-values are obtained by Welch’s t-test. (h) The *R*2 scores between the continuous representation and DPS for all inferred cell types are obtained through 10-fold cross-validation (mean and standard deviation). Notably, the continuous representation of glutamatergic cells has *R*2 *>* 0.5, whereas GABAergic cells exhibit smaller *R*2 values.

We next study changes to the discrete and continuous factors of variability across the spectrum of AD pathology. For this analysis, we use the disease pseudo-progression score (DPS), which is a number between 0 and 1, increasing with disease progression and calculated per donor, capturing neuropathological changes in the MTG associated with AD [17]. By calculating the expected categorical assignment probabilities, *P*_*t*_(**c**), over cells belonging to a donor, we observe declines in the relative abundances of L2/3 IT and *Sst* neurons, i.e. their categorical probabilities, during early pathology (Fig. 5e, Fig. S20), suggesting selective death of those neurons during early stages of AD. In contrast, L4 IT and *Pvalb* neurons display reductions to their relative abundances during later stages of the disease (Fig. 5e, Fig. S20) [17]. The rate of cellular loss in Alzheimer’s disease can be summarized by the change in *P*_*t*_(**c**) between the early and late stages of the disease. Notably, a considerable fraction (35%) of the transcriptomic types demonstrate a significant reduction (*p <* 0.005, Welch’s t-test) as they transition from the early to the late stage of the disease.

Lastly, we conduct a non-linear regression analysis to explore the relationship between the continuous latent variables inferred by MMIDAS and the DPS (Methods, Fig. S20a). Fig 5f and Fig 5h reveal novel correlations between type-dependent continuous variables and the disease stage for IT neurons in the MTG. Similarly, Fig. S20c,d,e,f demonstrate that the within-type, continuous factor of variability, **s**, can also serve as an accurate predictor of the BRAAK score (staging of AD-associated neurofibrillary pathology) for IT neurons. In contrast, the continuous variability of GABAergic neurons does not exhibit significant predictive power over the DPS (*R*2 *<* 0.5, Fig 5f,h).

To assess the generalizability of the inferred mixture representation across donors, we omit data from a few randomly chosen donors during the training phase. Later, during testing, we use the trained model to predict outcomes for the excluded donors (Fig. S20b). Our results reveal meaningful predictions that are robust against donor-specific variations. We note that batch effects (e.g., due to the use of different technologies, protocols, or sites) should be removed before MMIDAS analysis and are not considered here.

## Discussion

While neurons in the adult brain presumably attain terminal discrete identities, their phenotypes nevertheless remain in flux to better serve the organism in an ever-changing environment. This exacerbates the challenge of analyzing cellular diversity, which is perhaps most confounding in the mammalian brain with hundreds, and perhaps thousands [57], of types of brain cells. In this work, we introduced a novel deep variational inference method, MMIDAS, for generalized mixture models that is especially suitable for high-dimensional data and large number of categories. It utilizes multiple autoencoders that are collaboratively trained to achieve a consensus discrete representation, enabling us to overcome various technical challenges associated with generalized mixture model inference. MMIDAS models cellular heterogeneity as a combination of discrete and continuous factors, where the continuous factors represent the variability around distinct cluster representatives.

We validated MMIDAS on four diverse datasets spanning multiple technologies, brain regions, conditions, and species. Our findings demonstrate that MMIDAS markedly enhances the identifiability and clusterability scores of the inferred cell types compared to reference clusters established via hierarchical application of dimensionality reduction and graph clustering or through its iterative probabilistic annotation. Importantly, we also observed that the numbers of cell types inferred by MMIDAS are smaller than that of the reference taxonomies. Instead, MMIDAS posits that some of the diversity previously attributed to discrete cell types is better explained by continuous, within-type variability, related to biological phenomena such as spatial gradients and disease progression. The role of mixture modeling in gaining biological insights from single-cell data is further explored in Supplementary Note 13.

Dissecting discrete and continuous factors of variability enables identification of key genes or other features associated with aspects of cellular identity or organism-level observables, such as disease progression. In multimodal settings, MMIDAS allows for the identification of biological factors that influence multiple modalities, as well as aspects that are specific to individual modalities. This integrated analysis provides a comprehensive and accurate description of cellular diversity. In both unimodal and multimodal analysis, if it is desirable to use existing cell type labels, one autoencoder in MMIDAS can be replaced to supply those available labels during training. This way, new cells can be analyzed into their discrete and continuous factors based on those available training labels.

The present approach has certain limitations for data without an underlying categorical basis. MMIDAS assumes that the underlying data manifold can be faithfully represented via a categorical variable that denotes the cluster identity (type), and a continuous variable that denotes the within-cluster variability. Consequently, datasets that deviate from this model are ill-suited for MMIDAS. Nevertheless, contemporary cell type analysis is based on the assumption of fundamental underlying types with variable state, as opposed to a continuum of states, and MMIDAS was developed supporting this approach. Another challenge is the choice of hyperparameters, which MMIDAS shares with other unsupervised approaches. Fig. S5 shows that MMIDAS’ performance is robust to the value of the coupling factor within a relatively large range. Supplementary Note 10 discusses selecting the dimensionalities of the latent factors. Supplementary Note 12 (Fig. S7) highlights the role of nonlinearity in learning precise and interpretable cell representations. Lastly, Table S3 shows that MMIDAS’ performance further improves with more VAE arms, although this produces diminishing returns when computational cost is taken into account.

Brain cells occupy a diverse set of locations in a complicated landscape of cellular phenotypes. As the signals they receive and the homeostatic needs change, cell states move within wells of attraction of that landscape to best help the organism they are a part of. However, identifying the wells of attraction and the shapes of those wells may become ambiguous or ill defined. We believe MMIDAS can help researchers to dissect such perplexing single-cell datasets in a principled and efficient manner.

## Methods

### Mixture model inference with single-arm autoencoder

For an observation **x** ∈ ℝ^*D*^, a VAE learns a generative model *p*_***θ***_ (**x**|**z**) and a variational distribution *q*_***ϕ***_ (**z**|**x**), where **z** ∈ ℝ^*M*^ is a latent variable with a parameterized distribution *p*(**z**) and *M* ≪ *D* [33]. Here, a mixture model structure is imposed on the latent variable by introducing a categorical latent variable **c**, denoting the class label, and a continuous latent variable **s**, which is conditioned on the discrete variable **c**. We refer to the continuous variable **s** as the *state* or *style* variable interchangeably. **c** and **s** together form the latent variables of each (mixture) autoencoder.

For completeness, we first derive the evidence lower bound (ELBO) [3] for a single-arm mixture variational autoencoders (VAE), in which an observation **x** can be described by one categorical random variable and one continuous random variable, with conditional dependency between **c** and **s**. The variational approach to inferring the latent variables corresponds to solving the optimization equation

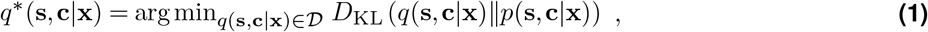

where 𝒟 is a family of density functions over the latent variables. Evaluating the objective function requires knowledge of *p*(**x**), which is usually unknown. Therefore, we rewrite the divergence term as

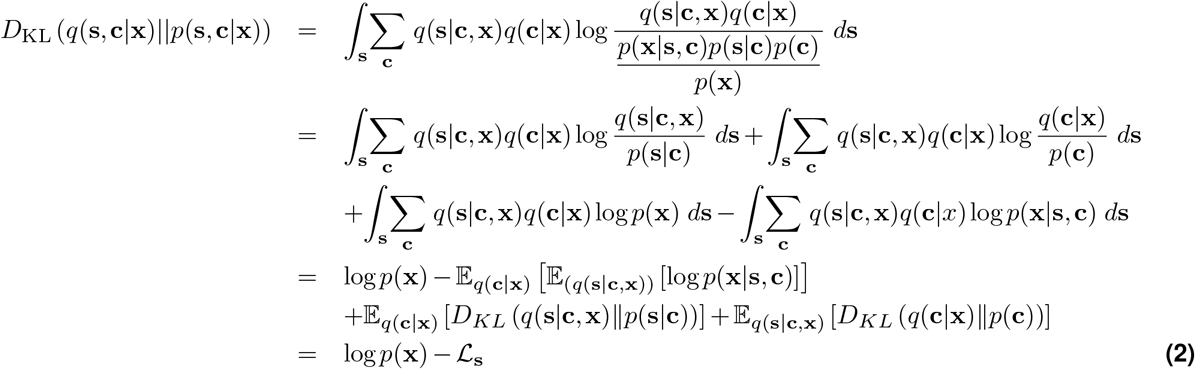

Since log*p*(**x**) is not function of the optimization parameters, instead of minimizing Eq. 2, the variational lower bound with the distributions parameterized by ***θ*** and ***ϕ***,

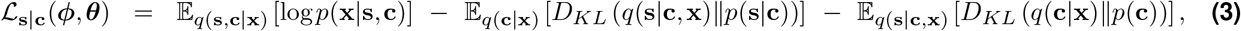

can be maximized. The first term in Eq. 3 is referred to as the (negative) “reconstruction loss” or ℒ_*rec*_. To generate a sample (conditioned on the observation **x**), first a discrete latent vector **c** is sampled from *q*(**c**|**x**). Given sample **c**, a continuous latent vector is sampled from *q*(**s**|**c, x**). The decoder outputs the parameters of *q*(**x**|**c, s**), from which we can sample an output vector.

Maximizing the ELBO in single-arm VAE as defined in Eq.3 imposes constraints on *q*(**s**|**c, x**) and *q*(**c**|**x**) that can lead to underestimation of posterior probabilities, known as the mode collapse problem, where the network disregards certain latent variables[41, 3]. Existing single VAE-based methods, such as JointVAE [12] and CascadeVAE [29], address this, primarily by enforcing a uniform structure on *p*(**c**). JointVAE [12] modifies the ELBO by assigning a pair of controlled information capacities for each variational factor, i.e. 𝒞_*s*_ ∈ ℝ^|**s**|^ and 𝒞_*c*_ ∈ ℝ^|**c**|^. Its performance heavily relies on the heuristic adjustment of these capacities, which scales poorly and may not be enough to prevent mode collapse in high-dimensional scenarios. Another alternative, CascadeVAE [29], tackles ELBO maximization through a semi-gradient-based approach, iteratively optimizing continuous and categorical variables separately. While this separation mitigates mode collapse concerns, it assumes uniform distribution of categorical variables. Hence, its efficacy depends on clusters exhibiting similar abundances within the dataset, which is typically not satisfied in single-cell analysis where the abundances of the clusters typically differ significantly. Thus, single-arm VAE fails to learn a precise and interpretable mixture representation for single-cell data (Fig. S6a,b).

### Mixture model inference with multi-arm autoencoders - MMIDAS

We introduce MMIDAS, a multi-arm VAE model designed to address the limitations of single-arm VAE models. Specifically, MMIDAS aims to overcome issues such as mode collapse of the categorical variable in high-dimensional settings without assuming uniform distribution on the categorical latent factor. MMIDAS consists of *A* mixture VAEs, i.e. *A*-tuple of independent and architecturally identical autoencoding arms.

The multi-arm framework of MMIDAS introduces a consensus constraint on the discrete factors of variability across the VAE models (arms) during the training process to address the aforementioned issues. In this framework, individual VAE arms receive a collection of non-identical copies, {**x**_*a*_, **x**_*b*_, …} of the given sample, **x**. These copies are sampled from a small neighborhood around **x** to remain within the same well of attraction and belong to the same category as **x**. (See Data augmentation below.) This novel approach serves to regularize the mixture representations by introducing variability during training, thus inferring a more robust and accurate model.

We choose *q*(**s**|**c, x**) to be a factorized Gaussian, parameterized using the reparametrization trick, and assume that the corresponding prior distribution is also a factorized Gaussian, **s**|**c** ∼ **𝒩** (0, **I**). For the categorical variable, instead of imposing a uniform structure on *p*(**c**) [12, 29], arms cooperate to learn *q*(**c**_*a*_ | **x**_*a*_), where **c**_*a*_ = **c**_*b*_ = ⃛, via a cost function at the time of training. Here, each arm has its own mixture representation with potentially non-identical parameters. Accordingly, the cost function of a set of VAEs with *A* arms can be formulated as a collection of constrained variational objectives as follows:

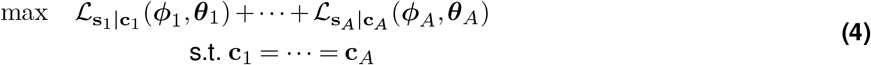

where 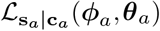 is the variational loss for a single arm *a* (Eq. 3). Propositions 1 and 2 (Supplementary Note 3) show that the shared categorical assignment inferred from *q*(**c**|**x**_1_, …, **x**_*A*_), under the **c** = **c**_1_ = … = **c**_*A*_ constraint of the multi-arm framework improves the accuracy of the categorical assignment on expectation, and having more arms increases the expected log posterior for the true categorical latent variable unless it is already at its maximum. Additionally, our theoretical results show that the required number of arms satisfying Eq. S35 is a function of the categorical distribution and the likelihood (Eq. S39, Supplementary Note 3). In the particular case of uniformly abundant categories, one pair of coupled arms is enough to satisfy Eq. S35 (Corollary 1, Supplementary Note 3).

We note that MMIDAS is an unsupervised method. As such, it does not require weak supervision, unlike [55, 4]. Instead, it relies on representations that are invariant under non-identical copies of observations. Moreover, unlike [37, 19], the multi-arm framework is not restricted to the continuous space.

### Sampler in variational inference

Variational inference approximates the posterior probability by using a computationally tractable sampler distribution. We utilize two distinct sampling techniques tailored to handle different types of latent variables. For continuous variables, we employ the widely used reparameterization trick [33], which enables efficient optimization through stochastic gradient descent. This method ensures differentiability of the sampling process with respect to the parameters of the distribution. For categorical variables, we employ the Gumbel-Softmax distribution [28], which similarly renders the sampling process differentiable with respect to the distribution’s parameters.

### Gumbel Distribution

A categorical distribution (CD) is a discrete probability distribution that describes the probability of a random variable that can take on one of *K* possible categories (classes). A categorical distribution is a generalization of the Bernoulli distribution and a special case of the multinomial distribution with *n* = 1, where category *c* is drawn with probability *π*_*c*_ from the finite probability set {*π*_1_, *π*_2_, …, *π*_*K*_}. The Gumbel distribution represents the distribution of extreme values (either maximum or minimum) of samples from various distributions. The cumulative distribution function (CDF) and the probability density function (PDF) of the Gumbel distribution for the largest values are defined as:

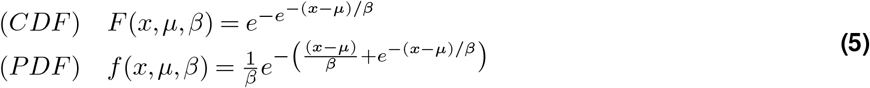

where *x, µ*, and *β* are the random variable, the location and the scale parameters, respectively.

#### Gumbel-Max Trick

A key trick in working with discrete distributions is to work with log(·) or logit(·) transformed data to avoid numerical issues due to small probability values.

The *Gumbel-Max trick* enables sampling from a CD given log(*π*_*i*_) or logit(*π*_*i*_). Unnormalized {log *π*_1_, …, log *π*_*K*_} values are first transformed to proper probabilities using the *softmax* function:

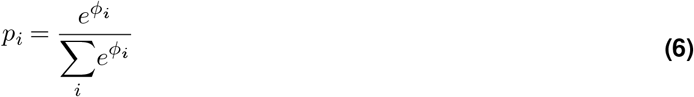

where *ϕ*_*i*_ = log *π*_*i*_ and 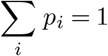.

Let *G*(*µ*) denotes the Gumbel distribution with parameters *µ* and *β* = 1. If *g*_1:*K*_ ∼ *G*(0) are independent samples taken from the standard Gumbel distribution parameterized by location *µ* = 0, we have

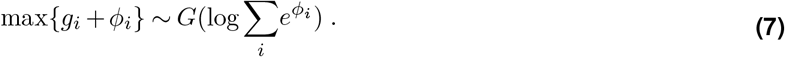

Accordingly, the maximum of the Gumbel distribution also comes from a Gumbel distribution as follows.

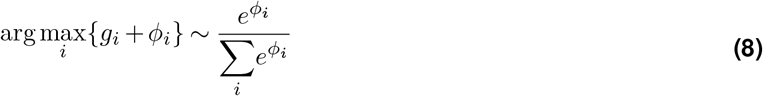

#### Gumbel-Softmax

To sample from a CD, similar to the reparameterization trick for continuous variables, we can consider the linear combination of the categorical variable with additive Gumbel noise and then take the `argmax`, which comes from the Gumbel distribution. Instead of using the *one-hot* operation for sampling, we can replace `argmax` with the softmax function and have,

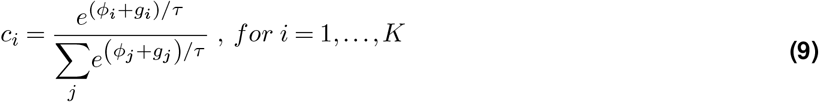

where its joint distribution is

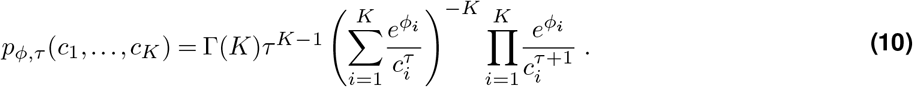

The probability representation in Eq. 10 is called the Gumbel-SoftMax distribution [28]. The parameter *τ* is a temperature parameter that allows us to control how closely samples from the Gumbel-Softmax distribution approximate those from the categorical distribution. As *τ* → 0, the softmax converges to the `argmax` function and the Gumbel-Softmax distribution converges to the categorical distribution. On the other hand, having *τ* ≫ 0 will provide a smooth approximation to the discrete uniform distribution.

### Reconstruction error

MMIDAS uses two different noise models for handling single-cell data in this paper:

i. Gaussian Noise Model: For log-transformed scRNA-seq data obtained via the Smart-seq technique and electrophysiological data, a multivariate normal distribution with equal variances across features can effectively approximate the single-cell measurements. To achieve this, gene expression values are first normalized to counts per million (CPM) and then transformed using the formula log (*CPM* + 1). In this case, the corresponding objective that minimizes the reconstruction error in Eq. 4 is the mean squared error (MSE).
ii. Zero-Inflated Negative Binomial (ZINB) Model: For scRNA-seq data obtained through the 10x platform, we use the ZINB model [14]:

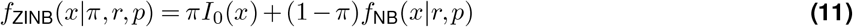

With

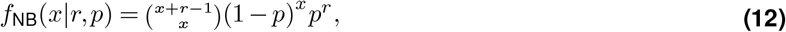

where *f*_ZINB_ and *f*_NB_ are probability mass functions for the count variable *x. I*_0_(.) is an indicator function, which equals 1 when *x* = 0, and *π* approximates the probability of gene dropout in 10x data. *r* and *p* denote the number and probability of success (not being a zero) in the NB distribution, respectively. In this case, MMIDAS uses the negative log likelihood of the ZINB distribution as the reconstruction loss in Eq. 4.

### Data augmentation

In MMIDAS, VAE arms receive non-identical observations that share the discrete variational factor. To achieve this in a fully unsupervised setting, we use *type-preserving* data augmentation that generates independent and identically distributed copies of data while preserving its categorical identity.

Augmentation can be considered as a generative process. Hence, we seek a generative model that learns the transformations representing within-class variability in an unsupervised manner. Learning such transformations is not straightforward: while conventional transformations such as rotation, scaling, or translation can serve as type-preserving augmentations for image data, they may not capture the richness of the underlying process. Moreover, such transformations cannot be used when within-class variability is unknown. Alternatives in the literature either rely on the availability of the class label, or are specific to image data [24, 27]. To this end, inspired by DAGAN [2], we propose an unsupervised augmentation scheme using a VAE-GAN-like architecture [36], parametrized by a neural network 𝒢, which implicitly learns the underlying conditional data distributions. 𝒢 should generate type-preserving samples (samples that remain within the well of attraction of the original observation) **x**_**a**_ conditioned on the given sample, **x**. This is achieved by concatenating the low dimensional representation of **x** with Gaussian noise **n** in 𝒢. To prevent 𝒢 from disregarding the noise, we formulate the training procedure as the following minmax optimization problem which uses a discriminator network 𝒟 as a regularizer:

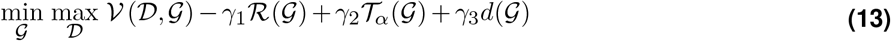

where,

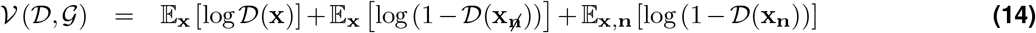

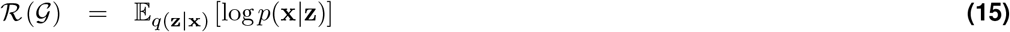

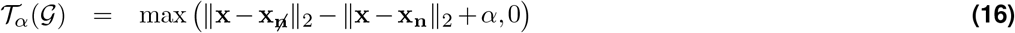

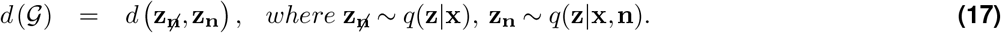

While training, 𝒢 generates two samples: **x**_**n**_ and 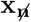. The former denotes **x**_**a**_, and the latter is a sample generated in the absence of noise. In Eq. 13, 𝒱 is the value function for the joint training of the discriminator and generator; ℛ is the reconstruction loss, which operates only over 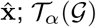 is the triplet loss that prevents network 𝒢 from disregarding noise and generating identical samples; and *d* (𝒢) is the Euclidean distance between the latent variables in the absence and presence of noise. *d* (𝒢) is a regularizer to encourage original and noisy samples to be located close to one another in the latent space. *γ*_1_, *γ*_2_, and *γ*_3_ are hyperparameters, each constrained to the range of 0 to 1. **z** denotes the latent representation of network 𝒢, not to be confused with the structured latent variables of the mixture VAE arms.

Importantly, the augmenter learns to generate samples in the vicinity of a given sample in the latent space, which is independent of **s** and **c**. The augmenter does not use any label information in any way. We call this a type-preserving augmenter not because label information was utilized during training or label preservation is guaranteed, but because the augmenter generates samples ‘similar’ to the original **x** that typically remain in the same well of attraction, i.e. belong to the same cluster.

In Supplementary Note 3, Remark 1 further discusses an under-exploration scenario in data augmentation, in which the augmented samples are not conditionally independently distributed and are concentrated around the given sample.

### Pairwise coupling

In MMIDAS, the mixture representation is obtained through the optimization in Eq. 4. Not only is it challenging to solve the maximization in Eq. 4 due to the equality constraint, but the objective remains a function of *p*(**c**) which is unknown, and typically non-uniform. To overcome this, we begin with an equivalent formulation for Eq. 4 by applying a pairwise coupling paradigm as follows (details of derivation in Supplementary Note 1):

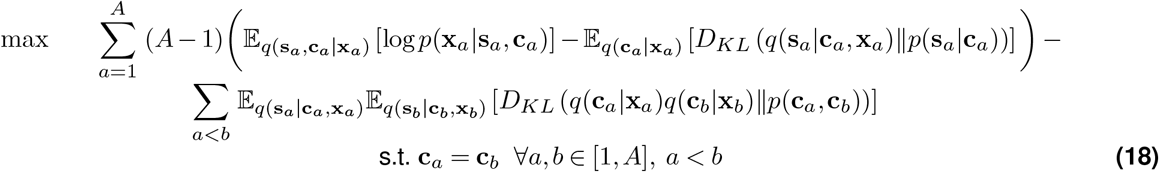

We next relax the optimization in Eq. 18 into an unconstrained problem by marginalizing the joint distribution over a mismatch measure between categorical variables (full derivation in Supplementary Note 2):

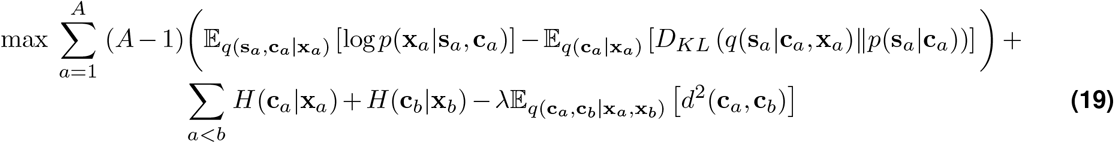

In Eq. 19, in addition to entropy-based confidence penalties known as mode collapse regularizers [44], the distance measure *d*(**c**_*a*_, **c**_*b*_) encourages a consensus on the categorical assignment controlled by *λ* ≥ 0, the coupling hyperparameter.

We refer to the model in Eq. 19 as MMIDAS. In this model, VAE arms try to achieve identical categorical assignments while independently learning their own style variables. In experiments, we set *λ* = 1 universally. While the bottleneck architecture already encourages interpretable continuous variables, this formulation can be easily extended to include an additional hyperparameter to promote disentanglement of continuous variables as in *β*-VAE [25]. Additional analyses to assess the sensitivity of the performance of MMIDAS to its coupling factor can be found in Supplementary Note 8.

In summary, we can cast the optimization in Eq. 19 in an equivalent a collection of constrained variational models as follows:

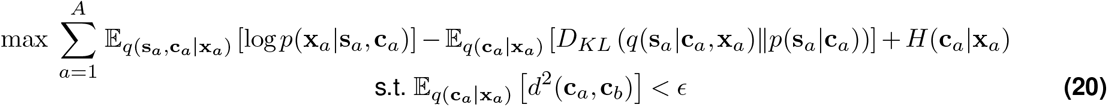

where *ϵ* denotes the strength of the consensus constraint. Here, **c**_*b*_ indicates the assigned category by any one of the arms, *b* ∈ {1,…, *A*}, imposing structure on the discrete variable to approximate its prior distribution.

### Distance between categorical variables

*d*(**c**_*a*_, **c**_*b*_) denotes the distance between a pair of |**c**| -dimensional un-ordered categorical variables, which are associated with probability vectors with non-negative entries and sum-to-one constraint that form a *K*-dimensional simplex, where *K* = |**c**|. In the real space, a typical choice to compute the distance between two vectors is using Euclidean geometry. However, this geometry is not suitable for probability vectors. Here, we utilize *Aitchison geometry* [1, 13], which defines a vector space on the simplex. Accordingly, the distance in the simplex, i.e. 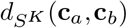 is defined as 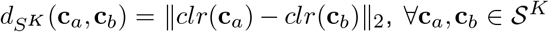, where *clr*(·) denotes the *isometric centered-log-ratio* transformation in the simplex. This categorical distance satisfies the conditions of a mathematical metric.

### Seeking consensus in the simplex

An instance of the mode collapse problem [39] manifests itself in the minimization of 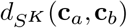 (Eq. 19): its trivial local optima encourages the network to abuse the discrete latent factor by ignoring many of the available categories. In the extreme case, the representations can collapse onto a single category; **c**_*a*_ = **c**_*b*_ = **c**_0_. In this scenario, the continuous variable is compelled to act as a primary latent factor. The model fails to deliver an interpretable mixture representation despite achieving an overall low loss value.

To avoid such undesirable local equilibria while training, we add perturbations to the categorical representation of each arm. If posterior probabilities in the simplex have small dispersions, the perturbed distance calculation overstates the discrepancies. Thus, instead of minimizing 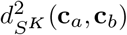), we minimize a perturbed distance 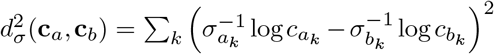. This corresponds to the distance between additively perturbed **c**_*a*_ and **c**_*b*_ vectors in Aitchison geometry. Here, 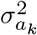 and 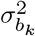 indicate the mini-batch variances of the *k*-th category, for arms *a* and *b*. We showed that the perturbed distance *d*_*σ*_(·) is bounded by 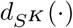 and non-negative values *ρ*_*u*_, *ρ*_*l*_ (Proposition 6, Supplementary Note 4). Accordingly, when **c**_*a*_ and **c**_*b*_ are similar and their spread is not small within the mini-batch, *d*_*σ*_(**c**_*a*_, **c**_*b*_) closely approximates 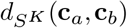. Otherwise, it diverges from 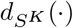 to avoid mode collapse.

#### Consensus score

To report the overall agreement between the categorical representations learned by each VAE arm, at the time of inference, we define the consensus score, *α*_*c*_ as follows.

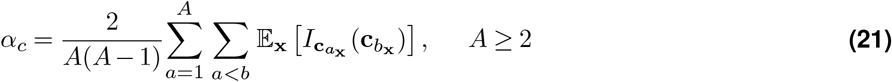

Where 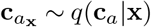, and 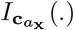 is an indicator function, which equals 1 when 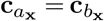. An ideal consensus score of 1 indicates that the VAE arms exhibit identical categorical representations for the discrete diversity present in the data.

### Clustering as mapping to a simplex

Clustering methods have been widely used to identify cell types. In MMIDAS, we define clustering as membership probability of each cell in a simplex. Vertices of the simplex, inferred as the discrete latent representation, formalize the notion of clusters (cell types). This clustering approach can be considered as a non-parametric technique, which offers flexible modeling of the data distribution for complex data structures.

### Over-parameterized simplex and network pruning to determine the number of discrete classes

Clustering is a classical ill-defined problem, popularly summarized as the lumpers vs splitters problem. Therefore, clustering methods, including non-parametric techniques, cannot determine the number of clusters truly automatically. Instead, the user either needs to provide an estimate of the count or have some prior knowledge about the underlying data structure, forming parameters or hyperparameters of the method. In the same vein, we propose an over-parameterization technique to determine the number of categories: at the beginning of the training, we initialize the dimensionality of the discrete latent space with a number larger than the expected number of cell types, e.g. if one expects around 100 types, set for instance 120 as an upper bound of the number of clusters. To choose the number of clusters, the model evaluates the importance or contribution of each vertex based on the consensus measure among arms. Those vertices which do not contribute with similar probability across arms are pruned. This iterative process continues until the model reaches a minimum level of consensus (a pre-defined hyperparameter) for all vertices in the simplex.

#### Algorithm 1

Mixture Model Inference with Discrete-coupled Autoencoders

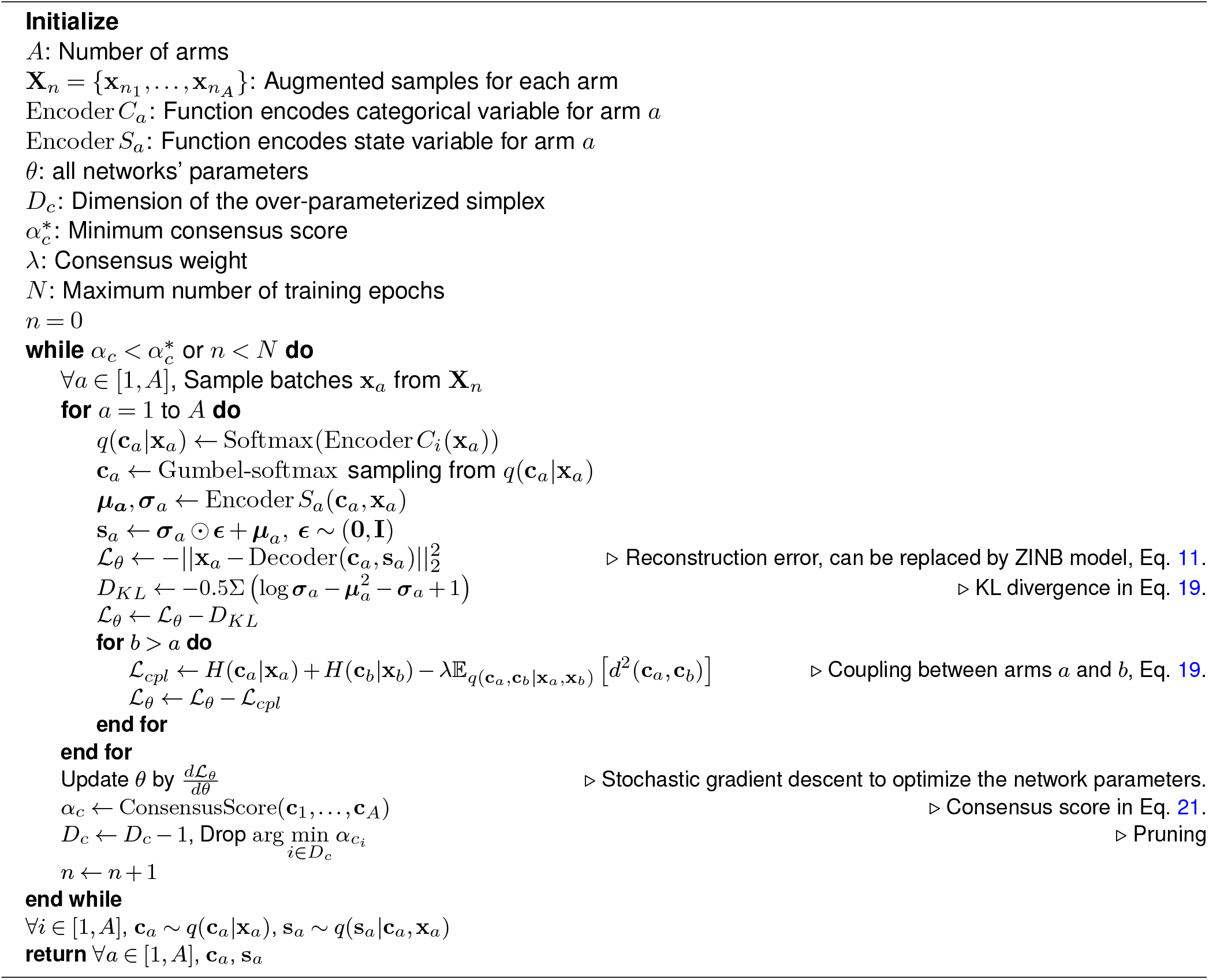

### Comparative analysis on cell types

Evaluating clustering performance without ground truth is inherently challenging. To address this, we conduct two comprehensive analyses to quantitatively assess stability and reproducibility of identified cell types within each dataset.

High-dimensional single-cell data presents a challenge due to the “curse of dimensionality”. This necessitates assessing cluster labels in a lower-dimensional space, as distance metrics become less informative for clustering in the original gene expression space (Fig. S13). We apply two distinct latent space embeddings: (i) a linear embedding via principal component analysis (PCA), akin to the method used in scrattch.hicat for mouse datasets; (ii) a nonlinear embedding, i.e. the latent space obtained by VAE, similar to the approach utilized in scVI / scANVI for defining Supertypes in SEA-AD.

Our first analysis involves a classification task to measure the post hoc identifiability of cluster labels through a 10-fold random train-test sampling approach. We train a Random Forest classifier on both linear and nonlinear embeddings and evaluate their performance on a test set. The average balanced (adjusted) accuracy score of the classifier is reported, which accounts for cluster imbalance. Both embeddings were obtained from the entire dataset. Notably, we did not observe data leakage from the test set to the training set (Table S5).

The second analysis involves a Silhouette analysis for each set of cell type labels. The Silhouette score provides a measure of how similar a given sample is to other samples in the same cluster as compared to samples in other clusters. The Silhouette score is a real value between − 1 and 1. A Silhouette score of 1 corresponds to ideal clusterability where samples in a cluster are identical to each other and different from samples in other clusters. For each set of cell type labels, we report the average Silhouette score per cell type, for the entire dataset.

For the analysis of the mouse Smart-seq and 10x datasets, we selected the first 100 principal components (PCs) to serve as our linear embedding. This choice is supported by testing across a range of 5 to 500 PCs, where 100 PCs provided the optimal performance (Fig. S12). The 10-dimensional latent space (denoted as *z* in Fig. S21b,c) used a non-linear embedding. For the Patch-seq dataset, the existing electrophysiological features are captured through PCA including 108 principal components, which summarize the time-series data, alongside 25 IPFX features (detailed further in the Dataset section). Accordingly, the performance metrics are reported based on a 30-dimensional nonlinear embedding for each modality. For the human Alzheimer’s disease dataset, again we utilized 100 PCs for the linear embedding and a 30-dimensional latent space for the nonlinear embedding.

In the Silhouette analysis, for each group of cell type labels, i.e. merged scrattch.hicat labels and MMIDAS labels, we report the results due to the embedding (linear and nonlinear) that produces the highest average performance. That is, we report the Silhouette scores based on the linear embedding for the merged scrattch.hicat labels, and based on the nonlinear embedding for the MMIDAS labels.

### Exploring continuous latent factors

To interpret and investigate the continuous representation obtained from the data, we conduct two sets of analysis, namely traversal analysis and regression analysis.

i. Traversal analysis: Here, we aim to understand the contribution of the continuous latent factors to the observed profiles, while holding the discrete factor fixed. We achieve this by modifying the continuous variables according to the approximate posterior, *q*(**s** | **c, x**). We explore the latent space, quantify how changes in the continuous variables affect the data, while keeping the discrete factor constant. For a given cell **x** = [*x*_1_, *x*_2_, …, *x*_*M*_]^*T*^ characterized by categorical type **c**, we change **s** along the direction that accounts for the maximum amount of variation. Subsequently, we calculate the normalized variation for each feature using the following formulation.

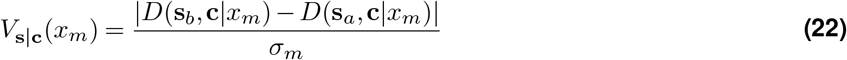

where *x*_*m*_ denotes the *m*-th feature of **x** (e.g., *m*-th gene), **s**_*a*_ = *µ*_*s* | *c*_ − *σ*_*s* | *c*_, **s**_*b*_ = *µ*_*s* | *c*_ + *σ*_*s* | *c*_, *µ*_*s* | *c*_ and *σ*_*s* | *c*_ denote the mean and standard deviation of of the continuous variable, respectively. *D* stands for the decoder function, such that 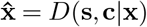. *σ*_*m*_ denotes the standard deviation of the *m*-th feature across all cells. We use the notation *V*_**s**_(*x*_*m*_) in the paper for ease of reference.
ii. Regression analysis: The regression analysis serves to explore the intricate relationships between the continuous latent factors and the meta-information associated with the data, such as brain regions or disease progression. In the mouse 10x dataset, we use a Random Forest regression model to quantify the predictability of the cortical regions from the continuous representation. Thus, this study explores the hypothesis that gene expression within a cell type can form spatial gradients in the brain.

For the SEA-AD dataset, we opt for a deep regression approach to better capture the intricate and flexible patterns that might exist between the continuous latent factors and the associated meta-data (Fig. S20a). In order to optimize the regression parameters, we use mean squared error (MSE) minimization, as follows:

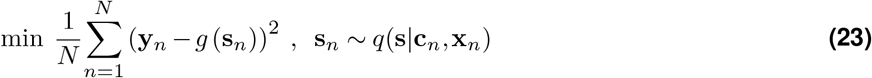

where the variable **y**_*n*_ represents the metadata associated with the *n*-th cell, such as the disease pseudo-progression score or the BRAAK score. *g* is a neural network parametrizing the nonlinear fit between **s** and **y**.

We quantify the quality of the fit between the latent variable and the meta-information with the coefficient of determination, *R*^2^.

### Datasets

#### Mouse Smart-seq dataset

This single-cell dataset was released as part of a transcriptomic cell types study [51]. RNA sequencing of 22,365 neurons from the anterolateral motor cortex (ALM) and primary visual cortex (VISp) regions of adult mice was performed with the Smart-seq (SSv4) platform. The cell isolation technique utilized for Smart-seq involved fluorescence-activated cell sorting (FACS). The reference transcriptomic taxonomy proposed for this dataset has 115 cortical neuronal types [51].

To focus on genes relevant to neuronal cell types, genes associated with sex or mitochondria, as well as those showing high expression in non-neuronal cells were removed. Subsequently, a subset of 5,032 highly variable genes (HVGs) was selected.

#### Mouse 10x isocortex dataset

This single-cell dataset was made available as part of a comprehensive transcriptomic study of the mouse cortex [56]. The dataset was generated using the 10xv2 RNA sequencing platform and includes 519,093 glutamatergic (excitatory) neurons and 94,493 GABAergic (inhibitory) neurons collected from seven cortical regions: anterior cingulate area (ACA), auditory cortex (AUD), primary motor cortex (MOp), retrosplenial cortex (RSP), the posterior parietal associative area (PTLp), primary somatosensory area (SSp), and visual cortex (VIS). The proposed transcriptomic cell type taxonomy for this dataset encompasses 113 excitatory and 92 inhibitory cortical neurons. Similar to the gene selection approach used for the mouse Smart-seq dataset, a subset of 10,000 HVGs was selected as input for MMIDAS.

#### Patch-seq dataset

This multimodal dataset profiles both transcriptomic and electrophysiological features of neurons in the adult mouse visual cortex. Profiles of inhibitory neurons have already been released by [21], and those of the excitatory neurons will be available on the Brain map portal (https://portal.brain-map.org/). Following patch-clamp recordings to measure intrinsic electrophysiological features in individual cells, scRNA-seq was performed on the same cells using the SSv4 platform.

The reference taxonomy for this dataset includes 92 cell types [21], which was established by mapping the transcriptomic profiles of neurons to the taxonomy proposed in [51]. The dataset comprises data from a total of 5,165 cortical neurons, each characterized by both a gene expression vector and a set of electrophysiological features. For gene selection, a two-step filtering procedure was followed. In the first step, genes associated with sex or mitochondria, as well as those exhibiting high expression levels in non-neuronal cells were excluded. In the second step, we removed genes that showed strong associations with the experimental platform, thereby reducing any potential platform-related bias. After applying these steps, we obtained a subset of 1,252 HVGs for our analysis.

For the electrophysiological modality, following previous studies [21, 19], we applied PCA to each time series. We retained principal components sufficient to explain 99% of the data, resulting in a total of 108 PCs to summarize the electrophysiological time series. We also included an additional 25 measurements of intrinsic physiological features that were computed by the IPFX library (https://ipfx.readthedocs.io/). To ensure consistency across experiments, the PCs were scaled to have unit variance, while the remaining IPFX features were individually normalized to have zero mean and unit norm. The following abbreviations are used for labeling electrophysiological features: action potential (AP), first action potential (AP1), inter-spike interval (ISI), amplitude (Amp), membrane potential (V), current (i), long square (LS), and short square (SS). A more detailed description of each electrophysiology feature can be found in the “Electrophysiology Overview” document available at https://help.brain-map.org/display/celltypes/Documentation.

#### Seattle Alzheimer’s disease dataset (SEA-AD)

This dataset is derived from the recently published [17] Seattle Alzheimer’s Disease Cell Atlas (SEA-AD) data, which is publicly accessible through the SEA-AD portal (https://SEA-AD.org/). The dataset consists of scRNA-seq data collected via 10xv3 from the Middle Temporal Gyrus (MTG) tissue of the human brain. Additionally, it includes essential meta-information such as cognitive score, BRAAK score, and demographic details.

The cohort consists of 84 donors covering a broad spectrum of Alzheimer’s disease (AD) pathology. The dataset also includes a continuous disease pseudo-progression score (DPS) per donor, ranging from low (score 0) to high (score 1), which was derived using a quantitative image-based neuropathology approach, enabling the ordered arrangement of donors based on their burden of AD pathology.

Here, our focus was on investigating four neuronal subclasses that are vulnerable to AD pathology: 299,563 L2/3 Intratelencephalic (IT) glutamatergic neurons, 152,052 L4 IT glutamatergic neurons, 45,872 Somatostatin-expressing (Sst) GABAergic neurons, and 79,976 Parvalbumin-expressing (Pvalb) GABAergic neurons. The dataset comprises 42 transcriptional types, referred to as “supertypes.” These supertypes were identified through an iterative projection of MTG data from young neurotypical reference donors [30] to itself, and by using scANVI [55] to predict the class label, as explained in [17].

In line with previous datasets, we performed a gene selection process to identify genes relevant to neuronal types. Genes associated with sex or mitochondria, as well as those exhibiting high expression in non-neuronal cells, were excluded. To focus on tissue-specific biology while reducing tissue-specific or technical variability, we independently selected the top 2000 HVGs for each donor. The union of these gene subsets was then utilized for our single-cell analysis. Both the gene selection and single-cell studies were carried out separately for each subclass in this dataset.

### KEGG Pathway and gene module database

In the continuous traversal study, we explored total variation across 18 pathways identified in the KEGG database [32]. Each pathway encompasses a gene module formed by the intersection of KEGG-suggested genes and those available in our datasets.

## Data and code availability

The mouse Smart-seq single-cell transcriptomic dataset is available at the NCBI Gene Expression Omnibus (GEO) under accession id GSE115746. The raw and processed mouse 10x isocortex dataset is deposited in the NeMO Archive for the BRAIN Initiative Cell Census Network, which is available at https://assets.nemoarchive.org/dat-jb2f34y. The inhibitory Patch-seq transcriptomic dataset is available at https://dandiarchive.org/dandiset/000020/. The SEA-AD dataset is available at https://portal.brain-map.org/explore/seattle-alzheimers-disease/seattle-alzheimers-disease-brain-cell-atlas-download?edit&language=en.

Code for MMIDAS implementation and analysis are available at https://github.com/AllenInstitute/MMIDAS. All taxonomies, list of genes, and additional metadata are available at https://github.com/AllenInstitute/MMIDAS/data. Collection of KEGG pathways from the KEGG database [32] along with the corresponding genes can be found at https://github.com/AllenInstitute/MMIDAS/blob/main/KEGG.

## Acknowledgements

We thank Polina Kosillo for help with cellular pathway analysis, Michael Hawrylycz and Bosiljka Tasic for helpful feedback on the manuscript, Zizhen Yao and Cindy van Velthoven for help with reference mouse cell type taxonomies, and Mariano Gabitto for help with the SEA-AD metadata. We wish to thank the Allen Institute for Brain Science founder, P. G. Allen, for his vision, encouragement and support. This work was supported by the NIH grants 1RF1MH128778-01, 1RF1MH125317-01, and 1U01NS132267-01.

