## Supplementary Information for "Joint inference of discrete cell types and continuous type-specific variability in single-cell datasets with MMIDAS"

#### Supplementary Note 1

##### Inference in MMIDAS

As discussed in the main text, the collective decision making for an A-arm VAE network can be formulated as an equality constrained optimization as follows:

$$\begin{aligned} \max \quad & \mathcal{L}(\boldsymbol{\phi}_1, \boldsymbol{\theta}_1, \mathbf{x}_1, \mathbf{s}_1, \mathbf{c}_1) + \cdots + \mathcal{L}(\boldsymbol{\phi}_A, \boldsymbol{\theta}_A, \mathbf{x}_A, \mathbf{s}_A, \mathbf{c}_A) \\ & \text{s.t. } \mathbf{c}_1 = \cdots = \mathbf{c}_A \end{aligned} \tag{S1}$$

Without loss of generality, the optimization in Eq. S1 can be rephrased as follows:

$$\begin{aligned} \max \quad & \mathcal{L}(\boldsymbol{\phi}_1, \boldsymbol{\theta}_1, \mathbf{s}_1, \mathbf{c}_1) + \mathcal{L}(\boldsymbol{\phi}_2, \boldsymbol{\theta}_2, \mathbf{s}_2, \mathbf{c}_2) + \cdots + \mathcal{L}(\boldsymbol{\phi}_A, \boldsymbol{\theta}_A, \mathbf{s}_A, \mathbf{c}_A) \\ & \text{s.t. } \mathbf{c}_1 = \mathbf{c}_2 \\ & \quad\;\; \mathbf{c}_1 = \mathbf{c}_3 \\ & \quad\;\; \cdots \\ & \quad\;\; \mathbf{c}_1 = \mathbf{c}_A \\ & \quad\;\; \cdots \\ & \quad\;\; \mathbf{c}_{A-1} = \mathbf{c}_A \end{aligned} \tag{S2}$$

where the equality constraint is represented as $\binom{A}{2}$ pairs of categorical agreements. Multiplying the objective term in Eq. S1 by a constant value, $A-1$, we obtain,

$$\begin{aligned} \max \quad & (A-1)\left(\mathcal{L}\left(\boldsymbol{\phi}_1, \boldsymbol{\theta}_1, \mathbf{s}_1, \mathbf{c}_1\right) + \mathcal{L}\left(\boldsymbol{\phi}_2, \boldsymbol{\theta}_2, \mathbf{s}_2, \mathbf{c}_2\right) + \cdots + \mathcal{L}\left(\boldsymbol{\phi}_A, \boldsymbol{\theta}_A, \mathbf{s}_A, \mathbf{c}_A\right)\right) \\ & \text{s.t. } \quad \mathbf{c}_a = \mathbf{c}_b \quad \forall a, b \in [1, A], a < b \end{aligned} \tag{S3}$$

Consider one pair of $\mathcal{L}$ objectives for two arms $a$ and $b$:

$$\begin{aligned} \mathcal{L}(\boldsymbol{\phi}_a, \boldsymbol{\theta}_a, \mathbf{s}_a, \mathbf{c}_a) + \mathcal{L}(\boldsymbol{\phi}_b, \boldsymbol{\theta}_b, \mathbf{s}_b, \mathbf{c}_b) &= \mathbb{E}_{q_{\boldsymbol{\phi}_a}(\mathbf{s}_a, \mathbf{c}_a|\mathbf{x}_a)}\left[\log p_{\boldsymbol{\theta}_a}(\mathbf{x}_a|\mathbf{s}_a, \mathbf{c}_a)\right] + \mathbb{E}_{q_{\boldsymbol{\phi}_b}(\mathbf{s}_b, \mathbf{c}_b|\mathbf{x}_b)}\left[\log p_{\boldsymbol{\theta}_b}(\mathbf{x}_b|\mathbf{s}_b, \mathbf{c}_b)\right] \\ & - \mathbb{E}_{q_{\boldsymbol{\phi}_a}(\mathbf{c}_a|\mathbf{x}_a)}\left[D_{KL}\left(q_{\boldsymbol{\phi}_a}(\mathbf{s}_a|\mathbf{c}_a, \mathbf{x}_a)\|p(\mathbf{s}_a|\mathbf{c}_a)\right)\right] - \mathbb{E}_{q_{\boldsymbol{\phi}_b}(\mathbf{c}_b|\mathbf{x}_b)}\left[D_{KL}\left(q_{\boldsymbol{\phi}_b}(\mathbf{s}_b|\mathbf{c}_b, \mathbf{x}_b)\|p(\mathbf{s}_b|\mathbf{c}_b)\right)\right] \\ & - \mathbb{E}_{q_{\boldsymbol{\phi}_a}(\mathbf{s}_a|\mathbf{c}_a, \mathbf{x}_a)}\left[D_{KL}\left(q_{\boldsymbol{\phi}_a}(\mathbf{c}_a|\mathbf{x}_a)\|p(\mathbf{c}_a)\right)\right] - \mathbb{E}_{q_{\boldsymbol{\phi}_b}(\mathbf{s}_b|\mathbf{c}_b, \mathbf{x}_b)}\left[D_{KL}\left(q_{\boldsymbol{\phi}_b}(\mathbf{c}_b|\mathbf{x}_b)\|p(\mathbf{c}_b)\right)\right] \end{aligned} \tag{S4}$$

When all arms receive augmented samples from the same original distribution, we have $p(\mathbf{c}_a) = p(\mathbf{c}_b) = p(\mathbf{c})$. Using the simplified notation, $q_a = q_{\boldsymbol{\phi}_a}(\mathbf{c}_a|\mathbf{x}_a)$, the last two KL divergence terms can be expressed as

$$\begin{aligned} D_{KL}\left(q_a\|p(\mathbf{c})\right) + D_{KL}\left(q_b\|p(\mathbf{c})\right) &= \sum_{\mathbf{c}_a} q_a \log \frac{q_a}{p(\mathbf{c})} + \sum_{\mathbf{c}_b} q_b \log \frac{q_b}{p(\mathbf{c})} \\ &= \sum_{\mathbf{c}_a}\sum_{\mathbf{c}_b} q_a q_b \log \frac{q_a}{p(\mathbf{c})} + \sum_{\mathbf{c}_a}\sum_{\mathbf{c}_b} q_a q_b \log \frac{q_b}{p(\mathbf{c})} \\ &= \sum_{\mathbf{c}_a}\sum_{\mathbf{c}_b} q_a q_b \log \frac{q_a q_b}{p(\mathbf{c})} \end{aligned} \tag{S5}$$

Now, if we marginalize $p(\mathbf{c})$ over the joint distribution $p(\mathbf{c}_a, \mathbf{c}_b)$, we can represent the categorical prior distribution as follows:

$$p(\mathbf{c}) = \sum_{\mathbf{c}_a, \mathbf{c}_b} p(\mathbf{c}|\mathbf{c}_a, \mathbf{c}_b)p(\mathbf{c}_a, \mathbf{c}_b) \tag{S6}$$

Since there is a categorical agreement condition i.e., $\mathbf{c}_a = \mathbf{c}_b$, $p(\mathbf{c})$ can be expressed as

$$p(\mathbf{c}) = \sum_{\mathbf{m}} p(\mathbf{c}|\mathbf{c}_a = \mathbf{c}_b = \mathbf{m}) p(\mathbf{c}_a = \mathbf{c}_b = \mathbf{m}) \tag{S7}$$

where

$$p(\mathbf{c}|\mathbf{c}_a = \mathbf{c}_b = \mathbf{m}) = \begin{cases} 1 & \mathbf{m} = \mathbf{c} \\ 0 & \text{otherwise} \end{cases} \tag{S8}$$

Accordingly, under the $\mathbf{c}_a = \mathbf{c}_b$ constraint, we merge those KL divergence terms as follows:

$$\begin{aligned} D_{KL}\left(q_a \| p(\mathbf{c})\right) + D_{KL}\left(q_b \| p(\mathbf{c})\right) &= \sum_{\mathbf{c}_a} \sum_{\mathbf{c}_b} q_a q_b \log \frac{q_a q_b}{p(\mathbf{c}_a, \mathbf{c}_b)} \\ &= D_{KL}(q_a q_b \| p(\mathbf{c}_a, \mathbf{c}_b)) \end{aligned} \tag{S9}$$

Finally, the optimization in Eq. S3 can be expressed as

$$\begin{aligned} \max \quad & \sum_{a=1}^{A} (A-1) \left( \mathbb{E}_{q(\mathbf{s}_a, \mathbf{c}_a | \mathbf{x}_a)} \left[ \log p(\mathbf{x}_a | \mathbf{s}_a, \mathbf{c}_a) \right] - \mathbb{E}_{q(\mathbf{c}_a | \mathbf{x}_a)} \left[ D_{KL}\left( q(\mathbf{s}_a | \mathbf{c}_a, \mathbf{x}_a) \| p(\mathbf{s}_a | \mathbf{c}_a) \right) \right] \right) - \\ & \sum_{a<b} \mathbb{E}_{q(\mathbf{s}_a, \mathbf{s}_b | \mathbf{c}_a, \mathbf{c}_b, \mathbf{x}_a, \mathbf{x}_b)} \left[ D_{KL}\left( q(\mathbf{c}_a | \mathbf{x}_a) q(\mathbf{c}_b | \mathbf{x}_b) \| p(\mathbf{c}_a, \mathbf{c}_b) \right) \right] \\ & \text{s.t.} \quad \mathbf{c}_a = \mathbf{c}_b \quad \forall a, b \in [1, A], a < b \end{aligned} \tag{S10}$$

### Supplementary Note 2

#### ELBO in MMIDAS

We first derive the evidence lower bound (ELBO) for the multi-arm VAE and show that this is equivalent to Eq. S10. Then, we derive a relaxation for the equality constrained optimization.

Given input data $\mathbf{x}_a$, an arm approximates two models $q(\mathbf{c}_a|\mathbf{x}_a)$ and $q(\mathbf{s}_a|\mathbf{x}_a, \mathbf{c}_a)$. If we use pairwise coupling to allow interactions between the arms, then, for a pair of VAE arms, $a$ and $b$, the variational lower bound obtained from the KL divergence can be generalized as

$$\begin{aligned}\Delta(a,b) &\triangleq D_{\mathrm{KL}}\left(q(\mathbf{s}_a,\mathbf{s}_b,\mathbf{c}_a,\mathbf{c}_b|\mathbf{x}_a,\mathbf{x}_b)\|p(\mathbf{s}_a,\mathbf{s}_b,\mathbf{c}_a,\mathbf{c}_b|\mathbf{x}_a,\mathbf{x}_b)\right)\\ &= \int_{\mathbf{s}_a}\int_{\mathbf{s}_b}\sum_{\mathbf{c}_a}\sum_{\mathbf{c}_b} q(\mathbf{s}_a,\mathbf{s}_b|\mathbf{c}_a,\mathbf{c}_b,\mathbf{x}_a,\mathbf{x}_b)q(\mathbf{c}_a,\mathbf{c}_b|\mathbf{x}_a,\mathbf{x}_b)\\ &\qquad \times \log\frac{q(\mathbf{s}_a,\mathbf{s}_b|\mathbf{c}_a,\mathbf{c}_b,\mathbf{x}_a,\mathbf{x}_b)q(\mathbf{c}_a,\mathbf{c}_b|\mathbf{x}_a,\mathbf{x}_b)}{\left(\frac{p(\mathbf{x}_a,\mathbf{x}_b|\mathbf{s}_a,\mathbf{s}_b,\mathbf{c}_a,\mathbf{c}_b)p(\mathbf{s}_a,\mathbf{s}_b|\mathbf{c}_a,\mathbf{c}_b)p(\mathbf{c}_a,\mathbf{c}_b)}{p(\mathbf{x}_a,\mathbf{x}_b)}\right)}\, d\mathbf{s}_a d\mathbf{s}_b \end{aligned} \tag{S11}$$

When each arm learns the continuous factor independent of other arms, we have $q(\mathbf{s}_a,\mathbf{s}_b|\mathbf{c}_a,\mathbf{c}_b,\mathbf{x}_a,\mathbf{x}_b) = q(\mathbf{s}_a|\mathbf{c}_a,\mathbf{x}_a)q(\mathbf{s}_b|\mathbf{c}_b,\mathbf{x}_b)$. Equivalently, for independent samples $\mathbf{x}_a$ and $\mathbf{x}_b$, we have $q(\mathbf{c}_a,\mathbf{c}_b|\mathbf{x}_a,\mathbf{x}_b) = q(\mathbf{c}_a|\mathbf{x}_a)q(\mathbf{c}_b|\mathbf{x}_b)$. Hence,

$$\begin{aligned}\Delta(a,b) = \log p(\mathbf{x}_a,\mathbf{x}_b) &+ \int_{\mathbf{s}_a}\int_{\mathbf{s}_b}\sum_{\mathbf{c}_a}\sum_{\mathbf{c}_b} q(\mathbf{s}_a|\mathbf{c}_a,\mathbf{x}_a)q(\mathbf{s}_b|\mathbf{c}_b,\mathbf{x}_b)q(\mathbf{c}_a|\mathbf{x}_a)q(\mathbf{c}_b|\mathbf{x}_b)\log\frac{q(\mathbf{c}_a|\mathbf{x}_a)q(\mathbf{c}_b|\mathbf{x}_b)}{p(\mathbf{c}_a,\mathbf{c}_b)}\, d\mathbf{s}_a d\mathbf{s}_b\\ &+\int_{\mathbf{s}_a}\sum_{\mathbf{c}_a} q(\mathbf{s}_a|\mathbf{c}_a,\mathbf{x}_a)q(\mathbf{c}_a|\mathbf{x}_a)\log\frac{q(\mathbf{s}_a|\mathbf{c}_a,\mathbf{x}_a)}{p(\mathbf{s}_a|\mathbf{c}_a)}\, d\mathbf{s}_a + \int_{\mathbf{s}_b}\sum_{\mathbf{c}_b} q(\mathbf{s}_b|\mathbf{c}_b,\mathbf{x}_b)q(\mathbf{c}_b|\mathbf{x}_b)\log\frac{q(\mathbf{s}_b|\mathbf{c}_b,\mathbf{x}_b)}{p(\mathbf{s}_b|\mathbf{c}_b)}\, d\mathbf{s}_b\\ &-\int_{\mathbf{s}_a}\sum_{\mathbf{c}_a} q(\mathbf{s}_a|\mathbf{c}_a,\mathbf{x}_a)q(\mathbf{c}_a|\mathbf{x}_a)\log p(\mathbf{x}_a|\mathbf{s}_a,\mathbf{c}_a)\, d\mathbf{s}_a - \int_{\mathbf{s}_b}\sum_{\mathbf{c}_b} q(\mathbf{s}_b|\mathbf{c}_b,\mathbf{x}_b)q(\mathbf{c}_b|\mathbf{x}_b)\log p(\mathbf{x}_b|\mathbf{s}_b,\mathbf{c}_b)\, d\mathbf{s}_b\end{aligned} \tag{S12}$$

$$\begin{aligned}\Delta(a,b) = &-\mathbb{E}_{q(\mathbf{c}_a|\mathbf{x}_a)}\left[\mathbb{E}_{q(\mathbf{s}_a|\mathbf{c}_a,\mathbf{x}_a)}\left[\log p(\mathbf{x}_a|\mathbf{s}_a,\mathbf{c}_a)\right]\right] - \mathbb{E}_{q(\mathbf{c}_b|\mathbf{x}_b)}\left[\mathbb{E}_{q(\mathbf{s}_b|\mathbf{c}_b,\mathbf{x}_b)}\left[\log p(\mathbf{x}_b|\mathbf{s}_b,\mathbf{c}_b)\right]\right]\\ &+\mathbb{E}_{q(\mathbf{c}_a|\mathbf{x}_a)}\left[D_{KL}\left(q(\mathbf{s}_a|\mathbf{c}_a,\mathbf{x}_a)\|p(\mathbf{s}_a|\mathbf{c}_a)\right)\right] + \mathbb{E}_{q(\mathbf{c}_b|\mathbf{x}_b)}\left[D_{KL}\left(q(\mathbf{s}_b|\mathbf{c}_b,\mathbf{x}_b)\|p(\mathbf{s}_b|\mathbf{c}_b)\right)\right]\\ &+\mathbb{E}_{q(\mathbf{s}_a|\mathbf{c}_a,\mathbf{x}_a)}\left[\mathbb{E}_{q(\mathbf{s}_b|\mathbf{c}_b,\mathbf{x}_b)}\left[D_{KL}\left(q(\mathbf{c}_a|\mathbf{x}_a)q(\mathbf{c}_b|\mathbf{x}_b)\|p(\mathbf{c}_a,\mathbf{c}_b)\right)\right]\right] + \log p(\mathbf{x}_a,\mathbf{x}_b)\end{aligned} \tag{S13}$$

Therefore, the variational lower bound for a pair of coupled VAE arms can be expressed as,

$$\begin{aligned}\mathcal{L}_{\mathrm{pair}}(a,b) = &\ \mathbb{E}_{q(\mathbf{s}_a,\mathbf{c}_a|\mathbf{x}_a)}\left[\log p(\mathbf{x}_a|\mathbf{s}_a,\mathbf{c}_a)\right] + \mathbb{E}_{q(\mathbf{s}_b,\mathbf{c}_b|\mathbf{x}_b)}\left[\log p(\mathbf{x}_b|\mathbf{s}_b,\mathbf{c}_b)\right]\\ &-\mathbb{E}_{q(\mathbf{c}_a|\mathbf{x}_a)}\left[D_{KL}\left(q(\mathbf{s}_a|\mathbf{c}_a,\mathbf{x}_a)\|p(\mathbf{s}_a|\mathbf{c}_a)\right)\right] - \mathbb{E}_{q(\mathbf{c}_b|\mathbf{x}_b)}\left[D_{KL}\left(q(\mathbf{s}_b|\mathbf{c}_b,\mathbf{x}_b)\|p(\mathbf{s}_b|\mathbf{c}_b)\right)\right]\\ &-\mathbb{E}_{q(\mathbf{s}_a|\mathbf{c}_a,\mathbf{x}_a)}\left[\mathbb{E}_{q(\mathbf{s}_b|\mathbf{c}_b,\mathbf{x}_b)}\left[D_{KL}\left(q(\mathbf{c}_a|\mathbf{x}_a)q(\mathbf{c}_b|\mathbf{x}_b)\|p(\mathbf{c}_a,\mathbf{c}_b)\right)\right]\right]\end{aligned} \tag{S14}$$

which is equivalent to the loss function in Eq. S10, for $A = 2$.

To compute the joint distribution $p(\mathbf{c}_a,\mathbf{c}_b)$, we first introduce an approximation by allowing $\mathbf{c}_a$ to take values in the probability simplex. We define an auxiliary continuous random variable $e$ representing the mismatch (error) between $\mathbf{c}_a$ and $\mathbf{c}_b$ such that $\forall \mathbf{c}_a,\mathbf{c}_b \in \mathcal{S}^K$, and $0 < \epsilon \ll 1$,

$$p(\mathbf{c}_a,\mathbf{c}_b|e) = \begin{cases}1 & |e - d^2(\mathbf{c}_a,\mathbf{c}_b)| < \epsilon/2\\ 0 & \text{otherwise}\end{cases} \tag{S15}$$

Here, $d(\mathbf{c}_a, \mathbf{c}_b)$ denotes the distance between $\mathbf{c}_a$ and $\mathbf{c}_b$ in the simplex $\mathcal{S}^K$, as a measure of mismatch between categorical variables. The random variable $e$ is distributed according to an exponential probability density function with parameter $\lambda$ i.e., $\forall e \geq 0,\ f(e,\lambda) = \lambda \exp(-\lambda e)$, where $\lambda > 0$. Accordingly, the joint categorical distribution can be represented as,

$$
\begin{aligned}
p(\mathbf{c}_a, \mathbf{c}_b) &= \int p(\mathbf{c}_a, \mathbf{c}_b | e) p(e) de && \text{(S16)}\\
&= \int_{-\epsilon/2+d^2(\mathbf{c}_a,\mathbf{c}_b)}^{\epsilon/2+d^2(\mathbf{c}_a,\mathbf{c}_b)} f(e,\lambda) de = \epsilon f\left(d^2\left(\mathbf{c}_a, \mathbf{c}_b\right), \lambda\right) + E && \text{(S17)}
\end{aligned}
$$

where $E$ is the error bound of the Midpoint integral rule. For given exponential function $f(e, \lambda)$, since $|f''(e,\lambda)| \leq \lambda^3$, $\forall e > 0$, the Midpoint approximation error is bounded by,

$$
|E| \leq \frac{(\lambda\epsilon)^3}{24}. \tag{S18}
$$

Subsequently, the joint probability distribution is equivalent to:

$$
p(\mathbf{c}_a, \mathbf{c}_b) = \epsilon\lambda \exp\left(-\lambda d^2\left(\mathbf{c}_a, \mathbf{c}_b\right)\right) + E \tag{S19}
$$

where $\epsilon$ and $\lambda$ are arbitrary constants. Therefore, we approximate the joint distribution as follows:

$$
p(\mathbf{c}_a, \mathbf{c}_b) \approx \epsilon\lambda \exp\left(-\lambda d^2\left(\mathbf{c}_a, \mathbf{c}_b\right)\right) \tag{S20}
$$

Thus, the last KL divergence in Eq. S14 can be approximated as

$$
\begin{aligned}
D_{\mathrm{KL}}\left(q(\mathbf{c}_a|\mathbf{x}_a)q(\mathbf{c}_a|\mathbf{x}_b)\|p(\mathbf{c}_a,\mathbf{c}_b)\right) &= \sum_{\mathbf{c}_a}\sum_{\mathbf{c}_b} q(\mathbf{c}_a|\mathbf{x}_a)q(\mathbf{c}_b|\mathbf{x}_b)\log\frac{q(\mathbf{c}_a|\mathbf{x}_a)q(\mathbf{c}_b|\mathbf{x}_b)}{p(\mathbf{c}_a,\mathbf{c}_b)} && \text{(S21)}\\
&= -H(\mathbf{c}_a|\mathbf{x}_a) - H(\mathbf{c}_b|\mathbf{x}_b) - \sum_{\mathbf{c}_a}\sum_{\mathbf{c}_b} q(\mathbf{c}_a|\mathbf{x}_a)q(\mathbf{c}_b|\mathbf{x}_b)\log p(\mathbf{c}_a,\mathbf{c}_b)\\
&\approx -H(\mathbf{c}_a|\mathbf{x}_a) - H(\mathbf{c}_b|\mathbf{x}_b) + \lambda\mathbb{E}_{q(\mathbf{c}_a|\mathbf{x}_a)}\mathbb{E}_{q(\mathbf{c}_b|\mathbf{x}_b)}\left[d^2\left(\mathbf{c}_a,\mathbf{c}_b\right)\right] - \log\epsilon\lambda && \text{(S22)}
\end{aligned}
$$

Therefore, the approximated variational cost for a pair of VAE arms can be written as follows:

$$
\begin{aligned}
\mathcal{L}_{\mathrm{pair}}(a,b) &= \mathbb{E}_{q(\mathbf{s}_a,\mathbf{c}_a|\mathbf{x}_a)}\left[\log p(\mathbf{x}_a|\mathbf{s}_a,\mathbf{c}_a)\right] + \mathbb{E}_{q(\mathbf{s}_b,\mathbf{c}_b|\mathbf{x}_b)}\left[\log p(\mathbf{x}_b|\mathbf{s}_b,\mathbf{c}_b)\right]\\
&\quad -\mathbb{E}_{q(\mathbf{c}_a|\mathbf{x}_a)}\left[D_{KL}\left(q(\mathbf{s}_a|\mathbf{c}_a,\mathbf{x}_a)\|p(\mathbf{s}_a|\mathbf{c}_a)\right)\right] - \mathbb{E}_{q(\mathbf{c}_b|\mathbf{x}_b)}\left[D_{KL}\left(q(\mathbf{s}_b|\mathbf{c}_b,\mathbf{x}_b)\|p(\mathbf{s}_b|\mathbf{c}_b)\right)\right]\\
&\quad +H(\mathbf{c}_a|\mathbf{x}_a) + H(\mathbf{c}_b|\mathbf{x}_b) - \lambda\mathbb{E}_{q(\mathbf{c}_a|\mathbf{x}_a)}\mathbb{E}_{q(\mathbf{c}_b|\mathbf{x}_b)}\left[d^2\left(\mathbf{c}_a,\mathbf{c}_b\right)\right] && \text{(S23)}
\end{aligned}
$$

Now, by extending $\mathcal{L}_{pair}$ from two arms to $A$ arms, in which there are $\binom{A}{2}$ paired networks, the total loss function for $A$ arms can be written as

$$
\begin{aligned}
\mathcal{L}_{\mathrm{cpl}} &= \sum_{a=1}^{A-1}\sum_{b=a+1}^{A}\mathcal{L}_{\mathrm{pair}}(a,b)\\
&= \sum_{a=1}^{A}(A-1)\mathbb{E}_{q(\mathbf{s}_a,\mathbf{c}_a|\mathbf{x}_a)}\left[\log p(\mathbf{x}_a|\mathbf{s}_a,\mathbf{c}_a)\right] - (A-1)\mathbb{E}_{q(\mathbf{c}_a|\mathbf{x}_a)}\left[D_{KL}\left(q(\mathbf{s}_a|\mathbf{c}_a,\mathbf{x}_a)\|p(\mathbf{s}_a|\mathbf{c}_a)\right)\right]\\
&\quad +\sum_{a<b} H(\mathbf{c}_a|\mathbf{x}_a) + H(\mathbf{c}_b|\mathbf{x}_b) - \lambda\mathbb{E}_{q(\mathbf{c}_a|\mathbf{x}_a)}\mathbb{E}_{q(\mathbf{c}_b|\mathbf{x}_b)}\left[d^2\left(\mathbf{c}_a,\mathbf{c}_b\right)\right]. && \text{(S24)}
\end{aligned}
$$

### Supplementary Note 3

#### Collective learning with multi-arm mixture VAEs

**Proposition 1.** *Consider the problem of mixture representation learning in the multi-arm VAE framework. For independent samples from category* $\mathbf{m}$*, i.e.* $\mathbf{x}_i \sim p(\mathbf{x}|\mathbf{m})$*,*

$$\begin{aligned} \mathbb{E}_{q(\mathbf{x}|\mathbf{m})}\left[\log q(\mathbf{c}=\mathbf{m}|\{\mathbf{x}_i\}_{1:A})\right] \quad &> \quad \mathbb{E}_{q(\mathbf{x}|\mathbf{m})}\left[\log q(\mathbf{c}=\mathbf{m}|\{\mathbf{x}_i\}_{1:B})\right] \\ s.t.\ \mathbf{c}=\mathbf{c}_1=\cdots=\mathbf{c}_A \qquad & \qquad s.t.\ \mathbf{c}=\mathbf{c}_1=\cdots=\mathbf{c}_B \end{aligned} \tag{S25}$$

*if* $q(\mathbf{m}|\mathbf{x}_i) < 1$ *and* $A > B \geq 1$ *denote the number of arms.*

*Proof.* Given sample $\mathbf{x}$ with categorical variable $\mathbf{m}$, in the multi-arm framework, each arm receives a noisy copy $\mathbf{x}_i$ with the same categorical factor $\mathbf{c}=\mathbf{c}_1=\cdots=\mathbf{c}_A$ for an $A$-arm VAE. Defining $\mathbf{X}_A := \{\mathbf{x}_i\}_{1:A}$ the approximated categorical log posterior can be expressed as

$$\begin{aligned} \log q(\mathbf{c}_1=\cdots=\mathbf{c}_A=\mathbf{c}|\mathbf{X}_A) &= \log\frac{q(\mathbf{x}_1|\mathbf{X}_A\setminus\{\mathbf{x}_1\},\mathbf{c})q(\mathbf{x}_2|\mathbf{X}_A\setminus\{\mathbf{x}_1,\mathbf{x}_2\},\mathbf{c})\dots q(\mathbf{x}_A|\mathbf{c})q(\mathbf{c})}{q(\mathbf{X}_A)} \\ &= \log\frac{q(\mathbf{x}_1|\mathbf{X}_A\setminus\{\mathbf{x}_1\},\mathbf{c})\dots q(\mathbf{x}_A|\mathbf{c})}{q(\mathbf{X}_A)} + \log q(\mathbf{c}) \end{aligned} \tag{S26}$$

where we used $q(\mathbf{c}) = q(\mathbf{c}_1=\cdots=\mathbf{c}_A=\mathbf{c})$ to simplify the notation. Since all samples are independently generated, i.e. $q(\mathbf{x}_i|\mathbf{c},\mathbf{x}_j) = q(\mathbf{x}_i|\mathbf{c})$, the categorical log probability can be simplified as

$$\begin{aligned} \log q(\mathbf{c}|\mathbf{X}_A) &= \log\prod_{i=1}^{A}\frac{q(\mathbf{x}_i|\mathbf{c})}{q(\mathbf{x}_i)} + \log q(\mathbf{c}) \\ &= \sum_{i=1}^{A}\log\frac{q(\mathbf{x}_i|\mathbf{c})}{q(\mathbf{x}_i)} + \log q(\mathbf{c}) \end{aligned} \tag{S27}$$

By computing the expectation of the log posterior with respect to the empirical distribution of noisy samples with categorical factor $\mathbf{m}$, we have

$$\mathbb{E}_{q(\mathbf{x}|\mathbf{m})}\left[\log q(\mathbf{c}|\mathbf{X}_A)\right] = \sum_{i=1}^{A}\mathbb{E}_{q(\mathbf{x}|\mathbf{m})}\left[\log\frac{q(\mathbf{x}_i|\mathbf{c})}{q(\mathbf{x}_i)}\right] + \log q(\mathbf{c}) \tag{S28}$$

Since all of the data are independently sampled, the expected log-likelihood values can be expressed as follows:

$$\sum_{i=1}^{A}\mathbb{E}_{q(\mathbf{x}|\mathbf{c})}\left[\log q(\mathbf{x}_i|\mathbf{c})\right] = A\mathbb{E}_{q(\mathbf{x}|\mathbf{c})}\left[\log q(\mathbf{x}|\mathbf{c})\right] \tag{S29}$$

Therefore, in the $A$-arm framework, the approximated expected log posterior probability for the joint categorical factor $\mathbf{m}$ is defined as,

$$\mathbb{E}_{q(\mathbf{x}|\mathbf{m})}\left[\log q(\mathbf{c}=\mathbf{m}|\mathbf{X}_A)\right] = A\mathbb{E}_{q(\mathbf{x}|\mathbf{m})}\left[\log\frac{q(\mathbf{x}|\mathbf{c}=\mathbf{m})}{q(\mathbf{x})}\right] + \log q(\mathbf{c}=\mathbf{m}). \tag{S30}$$

According to Eq. S30, for a single arm (1-arm), we have

$$\begin{aligned} \mathbb{E}_{q(\mathbf{x}|\mathbf{m})}\left[\log q(\mathbf{c}=\mathbf{m}|\mathbf{x})\right] &= \mathbb{E}_{q(\mathbf{x}|\mathbf{m})}\left[\log\frac{q(\mathbf{x}|\mathbf{c}=\mathbf{m})}{q(\mathbf{x})}\right] + \log q(\mathbf{c}=\mathbf{m}), \\ &= D_{KL}\left(q(\mathbf{x}|\mathbf{c}=\mathbf{m})\|q(\mathbf{x})\right) + \log q(\mathbf{c}=\mathbf{m}) \end{aligned} \tag{S31}$$

and for a $B$-arm framework, we have

$$\mathbb{E}_{q(\mathbf{x}|\mathbf{m})}\left[\log q(\mathbf{c}=\mathbf{m}|\mathbf{X}_B)\right] \;=\; B\, D_{KL}\left(q(\mathbf{x}|\mathbf{c}=\mathbf{m})\|q(\mathbf{x})\right)+\log q(\mathbf{c}=\mathbf{m}). \tag{S32}$$

Since $D_{KL}\left(q(\mathbf{x}|\mathbf{m})\|q(\mathbf{x})\right) > 0$ (for more than one category), given $\mathbf{m}$, when for each individual arm $q(\mathbf{c}=\mathbf{m}|\mathbf{x}_i) < 1$ and $A > B \geq 1$, we have

$$A\, D_{KL}\left(q(\mathbf{x}|\mathbf{c}=\mathbf{m})\|q(\mathbf{x})\right) \;>\; B\, D_{KL}\left(q(\mathbf{x}|\mathbf{c}=\mathbf{m})\|q(\mathbf{x})\right), \tag{S33}$$

$$\mathbb{E}_{q(\mathbf{x}|\mathbf{m})}\left[\log q(\mathbf{c}=\mathbf{m}|\mathbf{x}_1,\cdots,\mathbf{x}_A)\right] \;>\; \mathbb{E}_{q(\mathbf{x}|\mathbf{m})}\left[\log q(\mathbf{c}=\mathbf{m}|\mathbf{x}_1,\cdots,\mathbf{x}_B)\right]. \tag{S34}$$

□

**Proposition 2.** *In the A-arm VAE framework, there exists an A that guarantees a true categorical assignment on expectation. That is,*

$$\mathbf{m}=\arg\max_{\mathbf{c}}\, \mathbb{E}_{q(\mathbf{x}|\mathbf{m})}\left[\log q(\mathbf{c}|\{\mathbf{x}_i\}_{1:A})\right], \quad s.t.\ \mathbf{c}=\mathbf{c}_1=\cdots=\mathbf{c}_A\ . \tag{S35}$$

*Proof.* In the $A$-arm framework, an accurate categorical assignment for all samples with the same categorical factor, e.g. $\mathbf{m}$, can be obtained on expectation, if and only if,

$$\mathbb{E}_{q(\mathbf{x}|\mathbf{m})}\left[\log q(\mathbf{c}_1=\cdots=\mathbf{c}_A=\mathbf{m}|\{\mathbf{x}_i\}_{1:A})\right] > \mathbb{E}_{q(\mathbf{x}|\mathbf{m})}\left[\log q(\mathbf{c}_1=\cdots=\mathbf{c}_A=\mathbf{n}|\{\mathbf{x}_i\}_{1:A})\right], \quad \forall \mathbf{n}\neq\mathbf{m},$$

where $\mathbf{m}$ is the ground-truth category. In case of the 1-arm framework, the correct categorical assignment receives the highest log posterior probability on expectation, if and only if,

$$\mathbb{E}_{q(\mathbf{x}|\mathbf{m})}\left[\log\frac{q(\mathbf{x}|\mathbf{c}=\mathbf{m})}{q(\mathbf{x}|\mathbf{c}=\mathbf{n})}\right] \;>\; \log\frac{q(\mathbf{c}=\mathbf{n})}{q(\mathbf{c}=\mathbf{m})}, \quad \forall \mathbf{n}\neq\mathbf{m} \tag{S36}$$

which is a function of categorical distributions and is not always satisfied for any arbitrary discrete distribution. According to Eq. S30, in the presence of $A$ VAE arms, we have

$$\mathbb{E}_{q(\mathbf{x}|\mathbf{m})}\left[\log q(\mathbf{c}_1=\cdots=\mathbf{c}_A=\mathbf{c}|\{\mathbf{x}_i\}_{1:A})\right] = A\mathbb{E}_{q(\mathbf{x}|\mathbf{m})}\left[\log q(\mathbf{x}|\mathbf{c})\right]+\log q(\mathbf{c})-A\mathbb{E}_{q(\mathbf{x}|\mathbf{m})}\left[\log q(\mathbf{x})\right]. \tag{S37}$$

In this framework, the accurate categorical assignment is obtained on expectation, if and only if,

$$\begin{aligned}
\mathbb{E}_{q(\mathbf{x}|\mathbf{m})}\left[\log q(\mathbf{c}_1=\cdots=\mathbf{c}_A=\mathbf{m}|\{\mathbf{x}_i\}_{1:A})\right] \;&>\; \mathbb{E}_{q(\mathbf{x}|\mathbf{m})}\left[\log q(\mathbf{c}_1=\cdots=\mathbf{c}_A=\mathbf{n}|\{\mathbf{x}_i\}_{1:A})\right],\\
A\mathbb{E}_{q(\mathbf{x}|\mathbf{m})}\left[\log q(\mathbf{x}|\mathbf{m})\right]+\log q(\mathbf{m})-A\mathbb{E}_{q(\mathbf{x}|\mathbf{m})}\left[\log q(\mathbf{x})\right] \;&>\; A\mathbb{E}_{q(\mathbf{x}|\mathbf{m})}\left[\log q(\mathbf{x}|\mathbf{n})\right]+\log q(\mathbf{n})-A\mathbb{E}_{q(\mathbf{x}|\mathbf{m})}\left[\log q(\mathbf{x})\right],\\
A\mathbb{E}_{q(\mathbf{x}|\mathbf{m})}\left[\log\frac{q(\mathbf{x}|\mathbf{c}=\mathbf{m})}{q(\mathbf{x}|\mathbf{c}=\mathbf{n})}\right] \;&>\; \log\frac{q(\mathbf{c}=\mathbf{n})}{q(\mathbf{c}=\mathbf{m})}.
\end{aligned} \tag{S38}$$

Thus, when the number of arms, $A$, satisfies

$$A \;>\; \max\left(\rho(\mathbf{m})D^{-1}(\mathbf{m}),1\right) \tag{S39}$$

where $\rho(\mathbf{m})=\max_{\mathbf{n}\neq\mathbf{m}}\log\frac{q(\mathbf{c}=\mathbf{n})}{q(\mathbf{c}=\mathbf{m})}$ and $D(\mathbf{m})=\min_{\mathbf{n}\neq\mathbf{m}}\ D_{KL}(q(\mathbf{x}|\mathbf{m})\|q(\mathbf{x}|\mathbf{n})))$, maximum of the expected log posterior probability belongs to the true categorical factor. □

**Corollary 1.** *For a uniform prior on the discrete factor, one pair of VAE arms ($A=2$) is sufficient to satisfy Eq. S35.*

*Proof.* For uniformly distributed clusters, $\forall\mathbf{m}$, $\rho(\mathbf{m})=0$. According to Eq. S39, for any $A\geq 2$, the accurate categorical assignment is satisfied. □

**Under-exploration**

We further study the under-exploration scenario in data augmentation, in which the noisy samples are concentrated around the given sample. Under this scenario, the proof of earlier Propositions follow in the same way except the augmented samples are no longer conditionally independent. This means that each augmented sample adds less than before to the expected log posterior. Yet, the same argument shows that there will be an $A$ for which the claims of the proposition are satisfied. This issue is discussed in Remark 1 as follows.

**Remark 1.** *When the augmentation is type-preserving, by definition,* $q(\mathbf{x}_i|\mathbf{x}_j, \mathbf{c}) = q(\mathbf{x}_i|\mathbf{c})$, *where* $\mathbf{x}_j$ *could be either the given training sample or another noisy copy. If the augmented samples concentrate around* $\mathbf{x}_j$, *i.e. the augmenter under-explores the category-conditioned distribution, the earlier proofs should be adapted by keeping the conditioning on* $\mathbf{x}_j$ *explicit. Conditionally independent terms used in Eq. S26 should be replaced by* $q(\mathbf{x}_i|\mathbf{x}_j, \mathbf{c})$ *as follows.*

$$\log q(\mathbf{c}|\mathbf{X}_A) = \log \frac{q(\mathbf{x}_1|\mathbf{x}_j, \mathbf{X} \setminus \{\mathbf{x}_1, \mathbf{x}_j\}, \mathbf{c}) \dots q(\mathbf{x}_A|\mathbf{x}_j, \mathbf{c}) q(\mathbf{x}_j|\mathbf{c}) q(\mathbf{c})}{q(\mathbf{X}_A)}. \tag{S40}$$

*Since all augmented samples are generated from sample* $\mathbf{x}_j$, *the conditional probability distribution can be simplified as follows.*

$$q(\mathbf{x}_i|\mathbf{x}_j, \mathbf{X}_A \setminus \{\mathbf{x}_i, \mathbf{x}_j\}, \mathbf{c}) = q(\mathbf{x}_i|\mathbf{x}_j, \mathbf{c}), \quad \textit{for } i \neq j \tag{S41}$$

*By computing the expectation of the log posterior with respect to the empirical distribution of noisy samples,* $q\left(\mathbf{X}_A|\mathbf{m}\right)$ *(samples are not independent anymore), the expected categorical log posterior probability for the true categorical factor can be defined as,*

$$\begin{aligned} \mathbb{E}_{q(\mathbf{X}_A|\mathbf{m})}\left[\log q(\mathbf{c}=\mathbf{m}|\mathbf{X}_A)\right] &= \mathbb{E}_{q(\mathbf{x}_i|\mathbf{x}_j,\mathbf{m})}\left[\log \prod_{i=1}^{A-1} \frac{q(\mathbf{x}_i|\mathbf{x}_j, \mathbf{c}=\mathbf{m})}{q(\mathbf{x}_i|\mathbf{x}_j)}\right] + \mathbb{E}_{q(\mathbf{x}_j|\mathbf{m})}\left[\log \frac{q(\mathbf{x}_j|\mathbf{m})}{q(\mathbf{x}_j)}\right] + \log q(\mathbf{c}=\mathbf{m}) \\ &= (A-1)\mathbb{E}_{q(\mathbf{x}_i|\mathbf{x}_j,\mathbf{m})}\left[\log \frac{q(\mathbf{x}_i|\mathbf{x}_j, \mathbf{c}=\mathbf{m})}{q(\mathbf{x}_i|\mathbf{x}_j)}\right] + \mathbb{E}_{q(\mathbf{x}_j|\mathbf{m})}\left[\log q(\mathbf{c}=\mathbf{m}|\mathbf{x}_j)\right] \end{aligned} \tag{S42}$$

*Based on Eq. S42, if the data augmenter only regenerates given sample, the expected log posterior probability in the A-arm framework is equal to the expected log posterior probability in the single framework,* $\mathbb{E}_{q(\mathbf{X}_A|\mathbf{m})}\left[\log q(\mathbf{c}=\mathbf{m}|\mathbf{X}_A)\right] = \mathbb{E}_{q(\mathbf{x}|\mathbf{m})}\left[\log q(\mathbf{c}=\mathbf{m}|\mathbf{x})\right]$.

### Supplementary Note 4

#### Aitchison geometry

In this section, we first briefly review some critical definitions in *Aitchison geometry.* Then, to support the proof of Proposition 6, here we introduce Lemma 1 and Propositions 4 and 5.

According to Aitchison geometry, a simplex of $K$ parts can be considered as a vector space $(\mathcal{S}^K, \oplus, \otimes)$, in which $\oplus$ and $\otimes$ corresponds to *perturbation* and *power* operations, respectively, as follows:

$$Perturbation: \ \forall \mathbf{x}, \mathbf{y} \in \mathcal{S}^K, \ \mathbf{x} \oplus \mathbf{y} = \mathcal{C}\left(x_1 y_1, \ldots, x_K y_K\right)$$

$$Power: \ \forall \mathbf{x} \in \mathcal{S}^K \ and \ \forall \alpha \in \mathbb{R}, \ \alpha \otimes \mathbf{x} = \mathcal{C}\left(x_1^\alpha, \ldots, x_K^\alpha\right)$$

where $\mathcal{C}$ denotes the closure operation as follows:

$$\mathcal{C}(\mathbf{x}) = \left( \frac{c x_1}{\sum_{k=1}^{K} x_k}, \ldots, \frac{c x_K}{\sum_{k=1}^{K} x_k} \right).$$

In the simplex vector space, for any $\mathbf{x}, \mathbf{y} \in \mathcal{S}^K$, the distance is defined as

$$d_{\mathcal{S}^K}\left(\mathbf{x}, \mathbf{y}\right) = \left( \frac{1}{K} \sum_{i<j} \left( \log \frac{x_i}{x_j} - \log \frac{y_i}{y_j} \right)^2 \right)^{1/2}. \tag{S43}$$

Furthermore, Aitchison has introduced *centered-logratio* transformation (CLR), which is an isometric transformation from a simplex to a $K-$dimensional real space, $clr(\mathbf{x}) \in \mathbb{R}^K$. The CLR transformation involves the logratio of each $x_k$ over geometric means in the simplex as follows.

$$clr(\mathbf{x}) = \left( \log \frac{x_1}{g(\mathbf{x})}, \ldots, \log \frac{x_K}{g(\mathbf{x})} \right). \tag{S44}$$

where $g(\mathbf{x}) = \left( \prod_{k=1}^{K} x_k \right)^{1/K}$ and $\sum_{k=1}^{K} \log \frac{x_k}{g(\mathbf{x})} = 0$.

Since CLR is an isometric transformation, we have

$$\begin{aligned} d_{\mathcal{S}^K}\left(\mathbf{x}, \mathbf{y}\right) &= d_{\mathbb{R}^K}\left(clr(\mathbf{x}), clr(\mathbf{y})\right) \\ &= \|clr(\mathbf{x}) - clr(\mathbf{y})\|_2 . \end{aligned}$$

The algebraic-geometric definition of $\mathcal{S}^K$ satisfies standard properties, such as

$$d_{\mathcal{S}^K}\left(\mathbf{x} \oplus \mathbf{v}, \mathbf{y} \oplus \mathbf{v}\right) = d_{\mathcal{S}^K}\left(\mathbf{x} \ominus \mathbf{v}, \mathbf{y} \ominus \mathbf{v}\right) = d_{\mathcal{S}^K}\left(\mathbf{x}, \mathbf{y}\right) \tag{S45}$$

where $\mathbf{v} \in \mathcal{S}^K$ could be any arbitrary vector in the simplex.

**Lemma 1.** *Given a set of vectors* $\{\mathbf{x}_1, \ldots, \mathbf{x}_N\} \in \mathcal{S}^K$ *where* $\mathcal{S}^K$ *is a simplex of* $K$ *parts,*

$$clr(\mathbf{x}_1 \oplus \mathbf{x}_2 \oplus \cdots \oplus \mathbf{x}_N) = clr(\mathbf{x}_1) + clr(\mathbf{x}_2) + \cdots + clr(\mathbf{x}_N).$$

*Proof.* According to Aitchison geometry, addition of vectors in the simplex is defined as,

$$\mathbf{x}_1 \oplus \cdots \oplus \mathbf{x}_N = \left( \frac{\prod_{n=1}^{N} x_{n_1}}{c_N}, \ldots, \frac{\prod_{n=1}^{N} x_{n_K}}{c_N} \right) \tag{S46}$$

where $c_N = \sum_{k=1}^{K} \prod_{n=1}^{N} x_{n_k}$.

By applying the centered-logratio transformation, we have

$$clr\left(\mathbf{x}_1 \oplus \cdots \oplus \mathbf{x}_N\right) = \left( \log \frac{\prod_{n=1}^{N} x_{n_1}}{\delta_{K,N}}, \ldots, \frac{\prod_{n=1}^{N} x_{n_K}}{\delta_{K,N}} \right) \tag{S47}$$

where $\delta_{K,N} = c_N \left( \prod_{k=1}^{K} \frac{\prod_{n=1}^{N} x_{n_k}}{c_N} \right)^{1/K} = \left( \prod_{k=1}^{K} \prod_{n=1}^{N} x_{n_k} \right)^{1/K}$.

Now, we can rewrite Eq. S47 as,

$$\begin{aligned} clr\left(\mathbf{x}_1 \oplus \cdots \oplus \mathbf{x}_N\right) &= \left( \log \frac{x_{1_1} \ldots x_{N_1}}{\left(\prod_k x_{1_k}\right)^{1/K} \cdots \left(\prod_k x_{N_k}\right)^{1/K}}, \ldots, \log \frac{x_{1_K} \ldots x_{N_K}}{\left(\prod_k x_{1_k}\right)^{1/K} \cdots \left(\prod_k x_{N_k}\right)^{1/K}} \right) \\ &= \left( \sum_n \log \frac{x_{n_1}}{\left(\prod_k x_{n_k}\right)^{1/K}}, \ldots, \sum_n \log \frac{x_{n_K}}{\left(\prod_k x_{n_k}\right)^{1/K}} \right) \\ &= clr(x_1) + \cdots + clr(x_N) \end{aligned} \tag{S48}$$

□

**Proposition 4.** *Given vectors $\mathbf{x}, \mathbf{y}, \mathbf{v}_x, \mathbf{v}_y \in \mathcal{S}^K$ where $\mathcal{S}^K$ is a simplex of $K > 0$ parts,*

$$d^2_{\mathcal{S}^K}\left(\mathbf{x}, \mathbf{y}\right) - \Gamma_l \leq d^2_{\mathcal{S}^K}\left(\mathbf{x} \oplus \mathbf{v}_x, \mathbf{y} \oplus \mathbf{v}_y\right) \leq d^2_{\mathcal{S}^K}\left(\mathbf{x}, \mathbf{y}\right) + \Gamma_u$$

*where $\Gamma_u, \Gamma_l \geq 0$, $\Gamma_u = K\tau_u^2 - \frac{\Delta^2}{K}$, $\Gamma_l = \frac{\Delta^2}{K} - K\tau_l^2$, $\tau_u = \max_k\{\log \frac{v_{x_k}}{v_{y_k}}\}$, $\tau_l = \max_k\{\log \frac{v_{y_k}}{v_{x_k}}\}$, and $\Delta = \sum_k \left( \log \frac{v_{x_k}}{v_{y_k}} \right)$.*

*Proof.* According to Aitchison geometry, the distance between two vectors $\mathbf{x}$, $\mathbf{y} \in \mathcal{S}^K$ is defined as

$$d^2_{\mathcal{S}^K}(\mathbf{x},\mathbf{y}) = \|clr(\mathbf{x}) - clr(\mathbf{y})\|_2^2$$

If we perturb vectors $\mathbf{x}$ and $\mathbf{y}$ by $\mathbf{v}_x$ and $\mathbf{v}_y$, the distance between the perturbed vectors in the simplex can be expressed as

$$d^2_{\mathcal{S}^K}\left(\mathbf{x}\oplus\mathbf{v}_x, \mathbf{y}\oplus\mathbf{v}_y\right) = \|clr(\mathbf{x}\oplus\mathbf{v}_x) - clr(\mathbf{y}\oplus\mathbf{v}_y)\|_2^2 \ .$$

According to Lemma 1,

$$\begin{aligned}
d^2_{\mathcal{S}^K}\left(\mathbf{x}\oplus\mathbf{v}_x, \mathbf{y}\oplus\mathbf{v}_y\right) &= \|\left(clr(\mathbf{x}) - clr(\mathbf{y})\right) + \Big(clr(\mathbf{v}_x) - clr\left(\mathbf{v}_y\right)\Big)\|_2^2 \\
&= \|clr(\mathbf{x}) - clr(\mathbf{y})\|_2^2 + \|clr(\mathbf{v}_x) - clr\left(\mathbf{v}_y\right)\|_2^2 + \\
&\quad \left(clr(\mathbf{x}) - clr(\mathbf{y})\right)^T\Big(clr(\mathbf{v}_x) - clr\left(\mathbf{v}_y\right)\Big) + \Big(clr(\mathbf{v}_x) - clr\left(\mathbf{v}_y\right)\Big)^T\left(clr(\mathbf{x}) - clr(\mathbf{y})\right) \\
&= d^2_{\mathcal{S}^K}(\mathbf{x},\mathbf{y}) + d^2_{\mathcal{S}^K}\left(\mathbf{v}_x,\mathbf{v}_y\right) + 2\sum_{k=1}^{K}\left(\log\frac{x_k}{g(\mathbf{x})} - \log\frac{y_k}{g(\mathbf{y})}\right)\left(\log\frac{v_{x_k}}{g(\mathbf{v}_x)} - \log\frac{v_{y_k}}{g(\mathbf{v}_y)}\right)
\end{aligned} \tag{S49}$$

For simplicity, define $d_1^2 = d^2_{\mathcal{S}^K}(\mathbf{x},\mathbf{y})$ and $d_2^2 = d^2_{\mathcal{S}^K}\left(\mathbf{x}\oplus\mathbf{v}_x, \mathbf{y}\oplus\mathbf{v}_y\right)$. Then,

$$\begin{aligned}
d_2^2 &= d_1^2 + d^2_{\mathcal{S}^K}\left(\mathbf{v}_x,\mathbf{v}_y\right) + 2\sum_{k=1}^{K}\left(\log\frac{x_k}{g(\mathbf{x})} - \log\frac{y_k}{g(\mathbf{y})}\right)\left(\log\frac{v_{x_k}}{g(\mathbf{v}_x)} - \log\frac{v_{y_k}}{g(\mathbf{v}_y)}\right) \\
&= d_1^2 + d^2_{\mathcal{S}^K}\left(\mathbf{v}_x,\mathbf{v}_y\right) + 2\sum_{k=1}^{K}\log\frac{x_k}{g(\mathbf{x})}\left(\log\frac{v_{x_k}}{v_{y_k}} - \log\frac{g(\mathbf{v}_x)}{g(\mathbf{v}_y)}\right) - 2\sum_{k=1}^{K}\log\frac{y_k}{g(\mathbf{y})}\left(\log\frac{v_{x_k}}{v_{y_k}} - \log\frac{g(\mathbf{v}_x)}{g(\mathbf{v}_y)}\right)
\end{aligned} \tag{S50}$$

Define $\log\dfrac{g(\mathbf{v}_x)}{g(\mathbf{v}_y)} = \log\dfrac{\left(\prod_k v_{x_k}\right)^{1/K}}{\left(\prod_k v_{y_k}\right)^{1/K}} = \dfrac{1}{K}\sum_k \log\dfrac{v_{x_k}}{v_{y_k}} = \dfrac{\Delta}{K}$. Then,

$$d_2^2 = d_1^2 + d^2_{\mathcal{S}^K}\left(\mathbf{v}_x,\mathbf{v}_y\right) + 2\sum_{k=1}^{K}\log\frac{v_{x_k}}{v_{y_k}}\left(\log\frac{x_k}{g(\mathbf{x})} - \log\frac{y_k}{g(\mathbf{y})}\right) - \frac{2\Delta}{K}\sum_{k=1}^{K}\left(\log\frac{x_k}{g(\mathbf{x})} - \log\frac{y_k}{g(\mathbf{y})}\right) \tag{S51}$$

Since CLR is a zero-mean transformation, $\sum_k \log\dfrac{x_k}{g(\mathbf{x})} = 0$ and $\sum_k \log\dfrac{y_k}{g(\mathbf{y})} = 0$. Therefore,

$$d_2^2 = d_1^2 + d^2_{\mathcal{S}^K}\left(\mathbf{v}_x,\mathbf{v}_y\right) + 2\sum_{k=1}^{K}\log\frac{v_{x_k}}{v_{y_k}}\left(\log\frac{x_k}{g(\mathbf{x})} - \log\frac{y_k}{g(\mathbf{y})}\right) \tag{S52}$$

Additionally, $d^2_{\mathcal{S}^K}\left(\mathbf{v}_x,\mathbf{v}_y\right) \geq 0$ can be expressed as,

$$\begin{aligned}
d^2_{\mathcal{S}^K}\left(\mathbf{v}_x,\mathbf{v}_y\right) &= \sum_{k=1}^{K}\left(\log\frac{v_{x_k}}{v_{y_k}} - \log\frac{g(\mathbf{v}_x)}{g(\mathbf{v}_y)}\right)^2 \\
&= \sum_{k=1}^{K}\left(\log\frac{v_{x_k}}{v_{y_k}}\right)^2 + \sum_{k=1}^{K}\left(\log\frac{g(\mathbf{v}_x)}{g(\mathbf{v}_y)}\right)^2 - 2\log\frac{g(\mathbf{v}_x)}{g(\mathbf{v}_y)}\sum_{k=1}^{K}\log\frac{v_{x_k}}{v_{y_k}} \\
&= \sum_{k=1}^{K}\left(\log\frac{v_{x_k}}{v_{y_k}}\right)^2 - \frac{\Delta^2}{K}
\end{aligned} \tag{S53}$$

Therefore,

$$d_2^2 \quad = d_1^2 + \sum_{k=1}^{K}\left(\log\frac{v_{x_k}}{v_{y_k}}\right)^2 - \frac{\Delta^2}{K} + 2\sum_{k=1}^{K}\log\frac{v_{x_k}}{v_{y_k}}\left(\log\frac{x_k}{g(\mathbf{x})} - \log\frac{y_k}{g(\mathbf{y})}\right) \tag{S54}$$

Now, consider $\tau_u = \max_k\{\log\frac{v_{x_k}}{v_{y_k}}\}$ and $\tau_l = \max_k\{\log\frac{v_{y_k}}{v_{x_k}}\} = -\min_k\{\log\frac{v_{x_k}}{v_{y_k}}\}$. Then,

$$\begin{aligned} d_2^2 &\le d_1^2 + K\tau_u^2 - \frac{\Delta^2}{K} + 2\tau_u\left(\sum_k\log\frac{x_k}{g(\mathbf{x})} - \sum_k\log\frac{y_k}{g(\mathbf{y})}\right) \\ d_2^2 &\ge d_1^2 + K\tau_l^2 - \frac{\Delta^2}{K} - 2\tau_l\left(\sum_k\log\frac{x_k}{g(\mathbf{x})} - \sum_k\log\frac{y_k}{g(\mathbf{y})}\right) \end{aligned} \tag{S55}$$

Again, because of $\sum_k\log\frac{x_k}{g(\mathbf{x})} = 0$ and $\sum_k\log\frac{y_k}{g(\mathbf{y})} = 0$, we conclude that,

$$d_1^2 - \frac{\Delta^2}{K} + K\tau_l^2 \le d_2^2 \le d_1^2 - \frac{\Delta^2}{K} + K\tau_u^2 \tag{S56}$$

$$d_1^2 - \Gamma_l \le d_2^2 \le d_1^2 + \Gamma_u \tag{S57}$$

Since $K\tau_u \ge \Delta \ge K\tau_l$, we conclude $\Gamma_u, \Gamma_l \ge 0$. □

**Proposition 5.** *Given samples $\mathbf{x}, \mathbf{y} \in \mathcal{S}^K$, where $\mathcal{S}^K$ is a simplex of $K$ parts, we have*

$$0 \le d_{\mathbf{v}}^2(\mathbf{x}, \mathbf{y}) - d_{\mathcal{S}^K}^2\left(\mathbf{x}\oplus\mathbf{v}_x, \mathbf{y}\oplus\mathbf{v}_y\right) \le \frac{1}{K}(\Delta + K\tau)^2$$

*where* $d_{\mathbf{v}}^2(\mathbf{x}, \mathbf{y}) = \sum_k\left(\log x_k v_{x_k} - \log y_k v_{y_k}\right)^2$, $\tau = \max_k\{\log\frac{x_k}{y_k}\}$, *and* $\Delta = \sum_k\left(\log\frac{v_{x_k}}{v_{y_k}}\right)$.

*Proof.*

$$\begin{aligned} d_{\mathcal{S}^K}^2\left(\mathbf{x}\oplus\mathbf{v}_x, \mathbf{y}\oplus\mathbf{v}_y\right) &= \sum_{k=1}^{K}\left(\log x_k v_{x_k} - \log y_k v_{y_k} - \frac{1}{K}\log\prod_k\frac{x_k v_{x_k}}{y_k v_{y_k}}\right)^2 \\ &= \sum_{k=1}^{K}\left(\log x_k v_{x_k} - \log y_k v_{y_k} - \frac{1}{K}\sum_k\log\frac{x_k v_{x_k}}{y_k v_{y_k}}\right)^2 \\ &= \sum_{k=1}^{K}\left(\log x_k v_{x_k} - \log y_k v_{y_k} - D\right)^2 \end{aligned} \tag{S58}$$

where $D = \frac{1}{K}\sum_k\left(\log x_k v_{x_k} - \log y_k v_{y_k}\right)$. Therefore,

$$\begin{aligned} d_{\mathcal{S}^K}^2\left(\mathbf{x}\oplus\mathbf{v}_x, \mathbf{y}\oplus\mathbf{v}_y\right) &= \sum_{k=1}^{K}\left(\log x_k v_{x_k} - \log y_k v_{x_k}\right)^2 + KD^2 - 2D\sum_{k=1}^{K}\left(\log x_k v_{x_k} - \log y_k v_{y_k}\right) \\ &= d_{\mathbf{v}}^2(\mathbf{x}, \mathbf{y}) - KD^2 \end{aligned}$$

$$d_{\mathbf{v}}^2(\mathbf{x}, \mathbf{y}) = d_{\mathcal{S}^K}^2\left(\mathbf{x}\oplus\mathbf{v}_x, \mathbf{y}\oplus\mathbf{v}_y\right) + KD^2 \tag{S59}$$

Since $KD^2 \geq 0$, $d^2_{\mathbf{v}}(\mathbf{x},\mathbf{y}) \geq d^2_{\mathcal{S}^K}\left(\mathbf{x}\oplus\mathbf{v}_x, \mathbf{y}\oplus\mathbf{v}_y\right)$.

Let $\tau = \max_k\{\log \frac{x_k}{y_k}\}$ and $\Delta = \sum_k \left(\log \frac{v_{x_k}}{v_{y_k}}\right)$. Then,

$$\begin{aligned} d^2_{\mathbf{v}}(\mathbf{x},\mathbf{y}) - d^2_{\mathcal{S}^K}\left(\mathbf{x}\oplus\mathbf{v}_x, \mathbf{y}\oplus\mathbf{v}_y\right) &= \frac{1}{K}\left(\sum_k \left(\log\frac{x_k}{y_k} + \log\frac{v_{x_k}}{v_{y_k}}\right)\right)^2 \\ &= \frac{1}{K}\left(\Delta + \sum_k\left(\log\frac{x_k}{y_k}\right)\right)^2 \\ &\leq \frac{1}{K}(\Delta + K\tau)^2 \ . \end{aligned} \tag{S60}$$

□

#### Consensus in the simplex

**Proposition 6.** *Suppose $\mathbf{c}_a, \mathbf{c}_b \in \mathcal{S}^K$, where $\mathcal{S}^K$ is a simplex of $K > 0$ parts. Let $d_{\mathcal{S}^K}(\mathbf{c}_a,\mathbf{c}_b)$ denote the distance in Aitchison geometry and $d^2_\sigma(\mathbf{c}_a,\mathbf{c}_b) = \sum_k \left(\sigma^{-1}_{a_k}\log c_{a_k} - \sigma^{-1}_{b_k}\log c_{b_k}\right)^2$ denote a perturbed distance. Then,*

$$d^2_{\mathcal{S}^K}(\mathbf{c}_a,\mathbf{c}_b) - \rho_l \leq d^2_\sigma(\mathbf{c}_a,\mathbf{c}_b) \leq d^2_{\mathcal{S}^K}(\mathbf{c}_a,\mathbf{c}_b) + \rho_u$$

*where $\rho_u, \rho_l \geq 0$, $\rho_u = K\left(\tau^2_{\sigma_u} + \tau^2_{\mathbf{c}}\right) + 2\Delta_\sigma\tau_{\mathbf{c}}$, $\rho_l = \frac{\Delta^2_\sigma}{K} - K\tau^2_{\sigma_l}$, $\tau_{\mathbf{c}} = \max_k\{\log c_{a_k} - \log c_{b_k}\}$, $\tau_{\sigma_u} = \max_k\{g_k\}$, $\tau_{\sigma_l} = \max_k\{-g_k\}$, $\Delta_\sigma = \sum_k g_k$, and $g_k = (\sigma^{-1}_{a_k} - 1)\log c_{a_k} - (\sigma^{-1}_{b_k} - 1)\log c_{b_k}$.*

*Proof.* Consider $\mathbf{x} = \mathbf{c}_a$, $\mathbf{y} = \mathbf{c}_b$, $\mathbf{v}_x = \mathbf{v}_a = \left(\frac{c_{a_1}^{(\sigma^{-1}_{a_1}-1)}}{\gamma_a}, \ldots, \frac{c_{a_K}^{(\sigma^{-1}_{a_K}-1)}}{\gamma_a}\right)$, and $\mathbf{v}_y = \mathbf{v}_b = \left(\frac{c_{b_1}^{(\sigma^{-1}_{b_1}-1)}}{\gamma_b}, \ldots, \frac{c_{b_K}^{(\sigma^{-1}_{b_K}-1)}}{\gamma_b}\right)$, where $\gamma_a = \sum_k c_{a_k}^{(\sigma^{-1}_{a_k}-1)}$ and $\gamma_b = \sum_k c_{b_k}^{(\sigma^{-1}_{b_k}-1)}$. We have

$$d^2_{\mathcal{S}^K}(\mathbf{c}_a\oplus\mathbf{v}_a, \mathbf{c}_b\oplus\mathbf{v}_b) = \sum_{k=1}^K \left(\sigma^{-1}_{a_k}\log c_{a_k} - \sigma^{-1}_{b_k}\log c_{b_k} - D\right)^2 \tag{S61}$$

where $D = \frac{1}{K}\sum_k \left(\sigma^{-1}_{a_k}\log c_{a_k} - \sigma^{-1}_{b_k}\log c_{b_k}\right)$. Using Propositions 4 and 5,

$$d^2_{\mathcal{S}^K}(\mathbf{c}_a,\mathbf{c}_b) + K\tau^2_{\sigma_l} - \frac{\Delta^2_\sigma}{K} \leq d^2_{\mathcal{S}^K}(\mathbf{c}_a\oplus\mathbf{v}_a, \mathbf{c}_b\oplus\mathbf{v}_b) \leq d^2_{\mathcal{S}^K}(\mathbf{c}_a,\mathbf{c}_b) + K\tau^2_{\sigma_u} - \frac{\Delta^2_\sigma}{K} \tag{S62}$$

and

$$0 \leq d^2_\sigma(\mathbf{c}_a,\mathbf{c}_b) - d^2_{\mathcal{S}^K}(\mathbf{c}_a\oplus\mathbf{v}_a, \mathbf{c}_b\oplus\mathbf{v}_b) \leq \frac{1}{K}(K\tau_{\mathbf{c}} + \Delta_\sigma)^2 \tag{S63}$$

Therefore,

$$d^2_{\mathcal{S}^K}(\mathbf{c}_a,\mathbf{c}_b) + K\tau^2_{\sigma_l} - \frac{\Delta^2_\sigma}{K} \leq d^2_\sigma(\mathbf{c}_a,\mathbf{c}_b) \leq d^2_{\mathcal{S}^K}(\mathbf{c}_a,\mathbf{c}_b) + \frac{1}{K}\left((K\tau_{\mathbf{c}} + \Delta_\sigma)^2 + K^2\tau^2_{\sigma_u} - \Delta^2_\sigma\right) \tag{S64}$$

$$d^2_{\mathcal{S}^K}(\mathbf{c}_a,\mathbf{c}_b) - \rho_l \leq d^2_\sigma(\mathbf{c}_a,\mathbf{c}_b) \leq d^2_{\mathcal{S}^K}(\mathbf{c}_a,\mathbf{c}_b) + \rho_u \ . \tag{S65}$$

□

### Supplementary Note 5

#### The MNIST benchmark

MNIST is one of the most widely used benchmark datasets in the field of machine learning. It contains a collection of 60K $28 \times 28$ images of handwritten digits that are classified into ten classes, corresponding to the digits 0 to 9. The dataset had been collected from a diverse set of contributors. As a result, the writing style of the digits (rotation, thickness, width, etc.) varies significantly across samples.

There is an extensive body of research on clustering in mixture models [3, 8, 10, 19, 14]. The idea of improving the clustering performance through seeking a consensus and *co-training* and *ensembling* across multiple observations has been explored in both unsupervised [16, 13] and semi-supervised contexts [1]. However, these methods do not consider the underlying continuous variabilities across observations. Moreover, unlike ensemble methods, which pool the results of different trained workers, autoencoding arms seek a consensus at the time of learning in our framework.

To enable comparison with other state-of-the-art approaches that can jointly infer discrete and continuous factors of variability, without any supervision, we train three state-of-the-art unsupervised VAE-based methods for mixture modeling: JointVAE [4], CascadeVAE [9], and MMIDAS (ours) (Table S1). Note that methods that focus only on clustering, image data, and the presence of a handful of clusters can achieve very high clustering accuracy for the MNIST dataset, e.g. ACOL with 98% [12]. However, these are not easily applicable to single cell data (non-image modality) and to joint inference of cell types and continuous factors in single-neuron datasets. Following the convention [4, 9, 2], each arm of MMIDAS uses a 10-dimensional categorical variable representing digits (type), and a 10-dimensional continuous random variable representing the writing style (state).

To study the interpretability of the mixture representations, (i) for the discrete latent factor, we report the accuracy (ACC) of discrete factors (categorical assignments) and the $D_{KL}(q(\mathbf{c})\|p(\mathbf{c}))$, (ii) for the continuous variable, we perform latent traversal analysis by fixing the discrete factor and changing the continuous variable according to $q(\mathbf{s}|\mathbf{c}, \mathbf{x})$. Additionally, we report the computational efficiency (number of iterations per second) to compare the training complexity of the multi-arm framework against earlier methods (Table S1). All reported numbers for MMIDAS models are average accuracies calculated across arms. In MMIDAS, each arm received an augmented copy of the original input generated by the deep generative augmenter (Supplementary Note 6) during training. Details of the network architectures and training settings can be found in Supplementary Note 14.

Table S1 displays the accuracy of the categorical assignment and the discrepancy between $q(\mathbf{c})$ and $p(\mathbf{c})$ for two 1-arm VAE methods (JointVAE and CascadeVAE), and MMIDAS with 2 arms.

| **Method** | **ACC** (%) $\uparrow$ (mean $\pm$ s.d.) | $D_{KL}(\times\ 100) \downarrow$ | **Computation** $\uparrow$ (iterations/sec) |
|---|---|---|---|
| JointVAE | 68.99 $\pm$ 11.8 | 3.12 $\pm$ 1.0 | 74.1 |
| CascadeVAE | 74.83 $\pm$ 06.9 | 1.99 $\pm$ 1.1 | 23.8 |
| MMIDAS | **84.56 $\pm$ 06.47** | **1.48 $\pm$ 1.4** | 17.5 |

Table S1: Test results for mixture VAE models trained on MNIST based on 10 randomly initialized runs. All models trained for $120K$ iterations. MMIDAS uses 2 arms. ACC and $D_{KL}$ denote the accuracy of the categorical assignment, and KL divergence between $q(\mathbf{c}|\mathbf{x})$ and $q(\mathbf{c})$, respectively. "Computation" reports the average number of iterations per second on a GeForce RTX 2080 Ti GPU, using batch size 256. The computation of MMIDAS includes the entire execution time for training one pair of coupled networks, plus data augmentation.

Fig. S1 illustrates the continuous latent traversals for four dimensions of the state variable inferred by MMIDAS, where each row corresponds to a different dimension of the categorical variable, and the continuous variable monotonically changes across columns. Results in Table S1 and Fig. S1 show that MMIDAS achieved an interpretable mixture representation with the highest categorical assignment accuracy. It outperforms earlier methods, without using extraneous optimization or heuristic channel capacities. Beyond performance

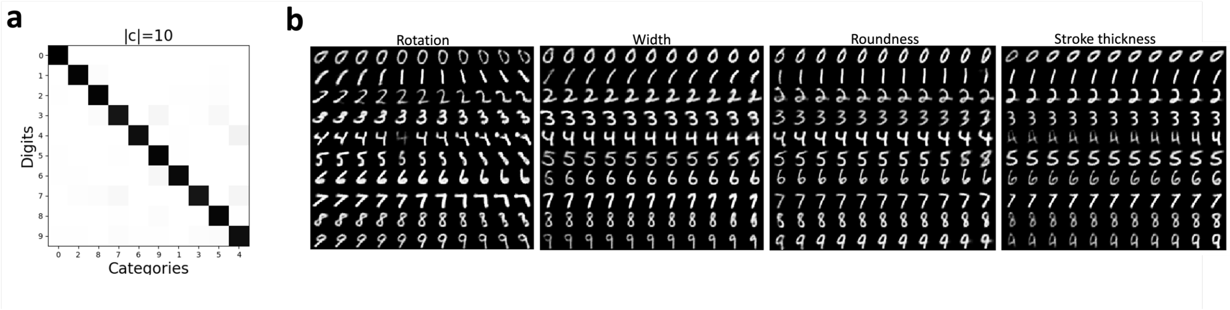


Figure S1: **Characterizing discrete and continuous latent factors in the MNIST dataset using MMIDAS.** Each arm's mixture representation is parameterized with 10-dimensional continuous and 10-dimensional categorical variables. (a) Confusion matrix comparing the true digit class labels against the categorical assignments inferred by MMIDAS. (b) Continuous latent traversals for the 1$^{\text{st}}$ arm of MMIDAS. Each row corresponds to a fixed discrete variable **c**. Each subfigure is associated with a different dimension of the continuous latent factor, in which each of the varied latent dimensions corresponds to an interpretable generative factor, such as rotation angle, width, roundness, and stroke thickness.

and robustness, its computational cost is also comparable to that of the baselines.

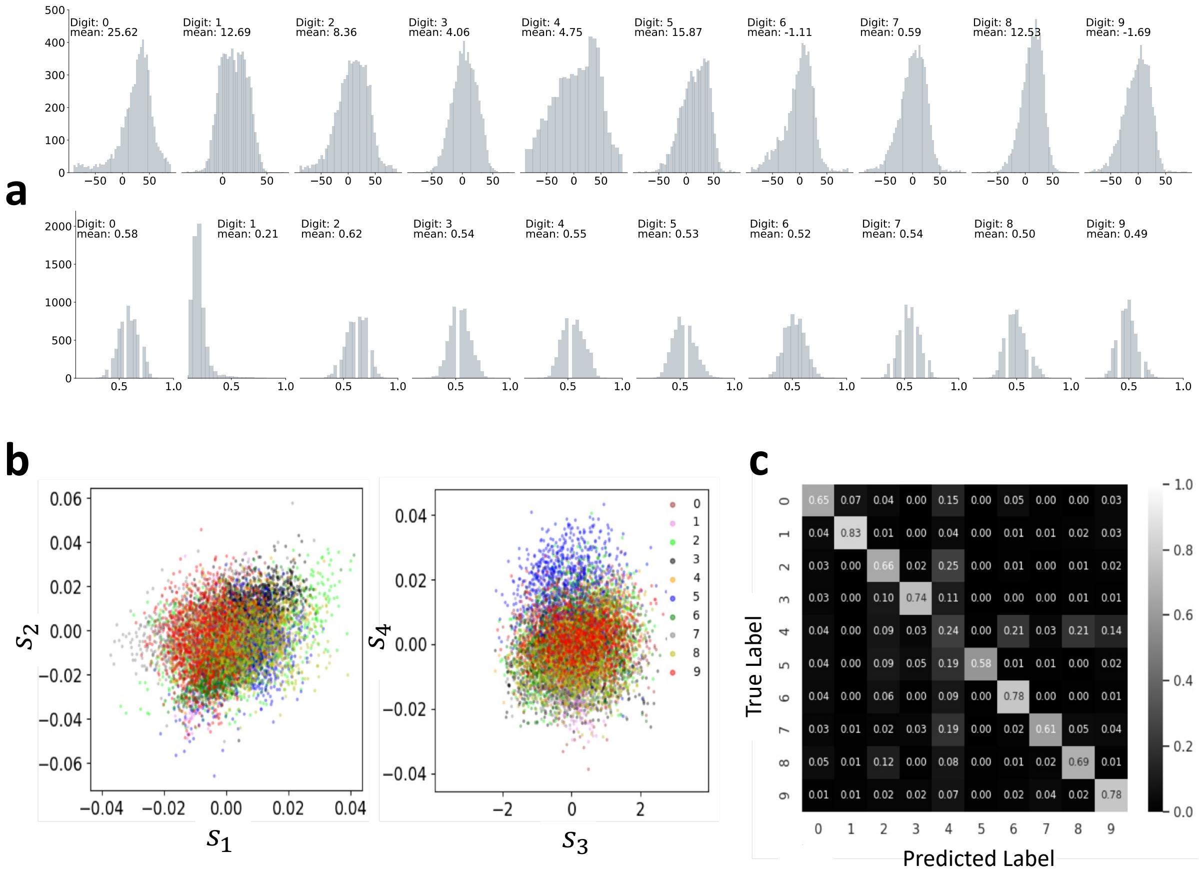


Figure S2: **Dependency of the continuous variable and the categorical type variable in the MNIST dataset.** (a) Histograms of angle and width for all digits in the MNIST dataset. The empirical distributions of rotation angle (top) and character width (bottom) are illustrated. The reported mean values and the shapes of the histograms illustrate the dependency of the state variable on the digit type. (b) 2-dimensional projections of the continuous variable obtained by a single-arm VAE, JointVAE. Each dot represents a sample of the MNIST dataset and colors represent different digits. (c) Confusion matrix for MNIST digit clustering using only the continuous latent variable learned by single-arm VAE.

### Supplementary Note 6

#### Type-preserving data augmentation

##### MNIST

Fig. S3a displays example noisy samples generated by the type-preserving augmentation for MNIST. To quantitatively evaluate the frequency with which the proposed type-preserving data augmentation preserves the underlying category, we use a benchmark classifier for MNIST digits, which achieves 99.54% accuracy over 10,000 test samples[1]. Applying the imported classifier to the augmented test samples yields 96.14% classification accuracy, which demonstrates that the augmenter preserves the label information (type) for **96.58**% of the augmented samples.

##### scRNA-seq

Generating augmented samples without altering the categorical identity can be challenging. In the case of image datasets, e.g. MNIST, intuitions on the identities of the discrete and continuous variational factors help to explicitly define a set of transformations, such as rotation, translation, scaling, flipping, that can be used as type-preserving augmentations. However, for non-image datasets, e.g. the single cell RNA-seq dataset, such intuition may not exist. Moreover, in case of biological datasets, learning an augmentation transformation is rather challenging due to the limited number of samples. Accordingly, in this section, we investigate the extent to which the proposed data augmentation method is successful in realistic generation of scRNA-seq samples.

Fig. S3c illustrates 2-dimensional representations for both the original and augmented samples, which were obtained by training a regular autoencoder: first, the autoencoder was trained on the original cell samples. After learning the latent representation, $\mathbf{z}$, for the original samples (left panel), we used the autoencoder to visualize the augmented samples (right panel). Comparing the visualizations demonstrates that these two representations are qualitatively similar and all groups of cells sharing the same type (same color) are placed in similar locations. Additionally, Fig. S3b shows the expression profiles of a subset of genes for an inhibitory cell, demonstrating qualitatively similar expression profiles. Since scRNA-seq data is heavily unbalanced, we additionally report the data augmenter's performance at the single gene expression level. Fig. S4 illustrates the expression distribution of a subset of known genes for augmented cell samples (colorful histograms) compared with the original expressions (gray histograms).

[1] Digit Recognizer, kaggle competition: https://www.kaggle.com/c/digit-recognizer

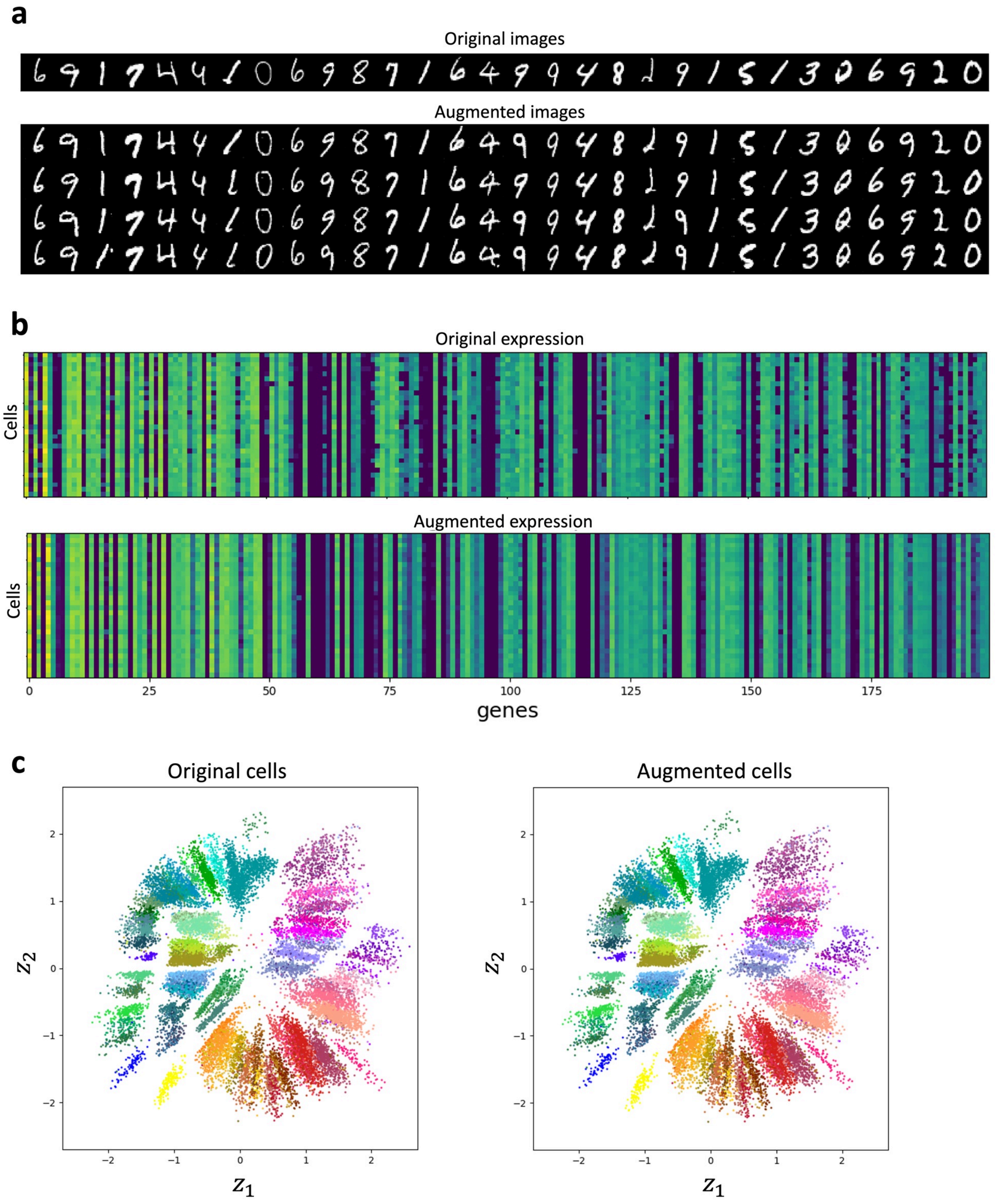


Figure S3: **Type-preserving data augmentation in MMIDAS.** (a) Augmented samples for the MNIST dataset. The samples generated by the type-preserving augmentation conserve the type of the original sample. (b) Qualitative comparison across the original and augmented gene expression profiles for a group of *Sst* neurons in the mouse Smart-seq dataset. (c) Low-dimensional visualization of the original and augmented cells in the mouse Smart-seq dataset. Both visualizations are obtained by a conventional autoencoder (used only for nonlinear dimensionality reduction). Each data point represents a cell in a 2-dimensional space. The left panel showcases original samples, while the right panel displays samples generated by the augmenter, presented within the same coordinate framework. Color coding is according to the proposed taxonomy in [18].

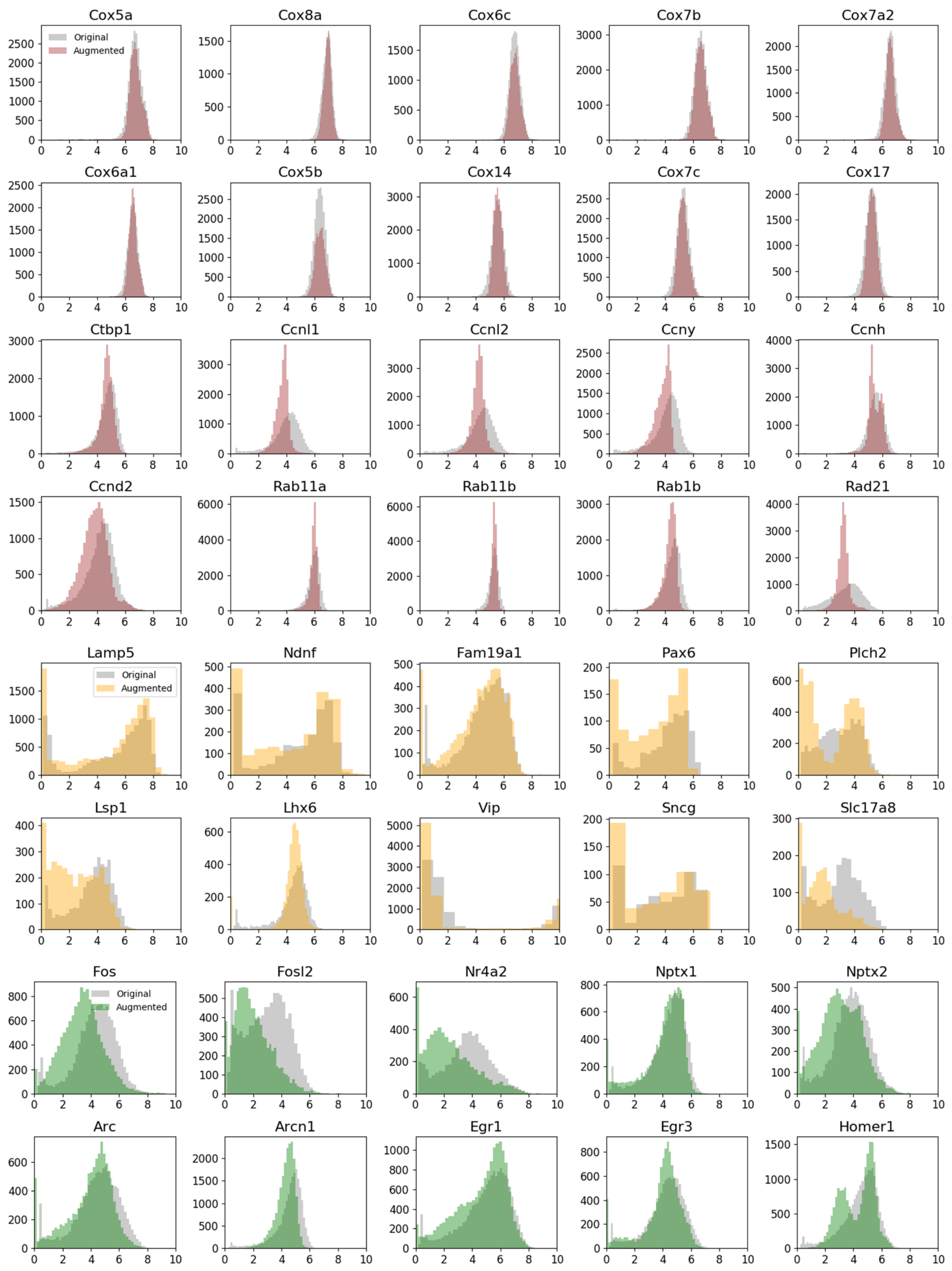


Figure S4: **Type-preserving data augmentation in scRNA-seq.** Comparison between the distribution of gene expression values in the original cell sample (gray color in all plots) and augmented samples for a subset of (a) immediate early genes (green), (b) house keeping genes (brown), and (c) marker genes (yellow).

### Supplementary Note 7

#### Ablation studies

To shed light on the robustness and accuracy of MMIDAS in mixture modeling, we investigate the categorical assignment performance under different training settings. Table S2 shows the performances of 1-arm mixture VAE (referred to as JointVAE [4]), MMIDAS and their variants under different training settings for the MNIST dataset. Here, we consider the reconstruction error ($\mathcal{L}_{rec}$) and the accuracy of the categorical assignments. In Table S2, JointVAE refers to the original JointVAE model that was trained with settings suggested in [4]; JointVAE$^{\dagger}$ is a JointVAE model that was trained with noisy copies of the original MNIST dataset generated by the type-preserving data augmentation method in Supplementary Note 6; JointVAE$^{\ddagger}$ is another JointVAE model that uses the same architecture for the basic encoder/decoder networks as the one used in MMIDAS. These results do not show any improvement in the performance of JointVAE by using data augmentation or altering the network architecture.

Next, we study the performance of the proposed MMIDAS model under three different settings. In Table S2, MMIDAS refers to the proposed 2-arm VAE framework using coupled-autoencoders and the type-preserving data augmentation in Supplementary Note 6; MMIDAS$^{*}$, is a MMIDAS model in which coupled networks are not independent and the network parameters are shared; MMIDAS$^{a}$, is a MMIDAS model that uses random rotations ($[-\pi/9, \pi/9]$) for data augmentation; MMIDAS ($\mathbf{s} \not\perp \mathbf{c}$), is a MMIDAS model in which the state variable is independent of the discrete variable. These results show that proposed model (MMIDAS) obtains the most accurate categorical assignments while the values are robust (i.e., consistently higher than those for the JointVAE variants) across all training settings.

| **Method** | $\mathcal{L}_{\text{rec}} \downarrow$ | **ACC** $\uparrow$ (mean $\pm$ s.d.) |
|---|---|---|
| JointVAE | 0.166 | 68.99 ± 11.76 |
| JointVAE$^{\dagger}$ | 0.166 | 68.21 ± 09.58 |
| JointVAE$^{\ddagger}$ | 0.162 | 62.19 ± 05.73 |
| MMIDAS | 0.145 | **84.56 ± 06.47** |
| MMIDAS$^{*}$ | 0.140 | 80.25 ± 05.37 |
| MMIDAS$^{a}$ | **0.135** | 82.92 ± 04.64 |
| MMIDAS($\mathbf{s} \not\perp \mathbf{c}$) | 0.146 | 79.63 ± 08.32 |

Table S2: Average reconstruction loss ($\mathcal{L}_{\text{rec}}$) and categorical assignment accuracy (ACC) in mixture representation learning under different training settings for 10 randomly initialized runs, for the MNIST dataset.

### Supplementary Note 8

#### Sensitivity to the consensus factor

Clustering is an ill-defined problem. Therefore, all models dealing with unsupervised grouping of samples include hyperparameters and MMIDAS is no exception. The MMIDAS framework has a regularization hyperparameter, $\lambda$, which controls the level of categorical consensus (coupling) between a pair of autoencoder arms. In this section, we conduct a series of experiments to assess the sensitivity of MMIDAS to its coupling factor, in comparison with JointVAE which has four critical hyperparameters, two for the discrete and two for the continuous variables. Fig. S5 shows how the mixture representation performance depends on the hyperparameters for both JointVAE and MMIDAS. For JointVAE, we only consider the channel capacity for the discrete variable, i.e. $\mathcal{C}_c$, which requires adjustment over training iterations.

Fig. S5a shows the categorical assignment accuracy as a function of $\lambda$ (for MMIDAS) and $\mathcal{C}_c$ (for JointVAE). While MMIDAS's performance is relatively insensitive to the coupling factor across a wide range, the performance of JointVAE depends strongly on the exact value of the channel capacity factor. Fig. S5b illustrates the categorical variables learned by JointVAE, when we reduce the maximum capacity from 5 to 1. Likewise, Fig. S5d shows a similar issue for JointVAE, when the maximum capacity is increased to 25. In contrast, MMIDAS's performance is relatively flat as a function of $\lambda$ within the range $[0.1, 10]$. Finally, adjusting the channel capacity throughout the training iterations for each dataset and latent space dimensionality is computationally expensive.

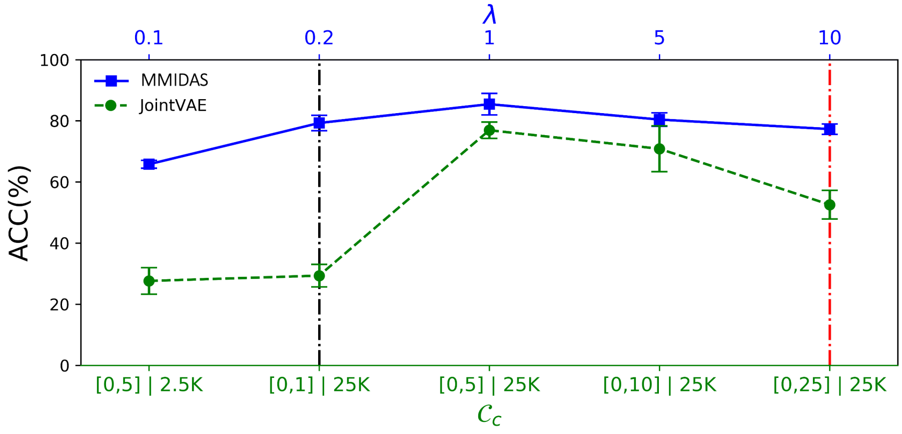


(a)

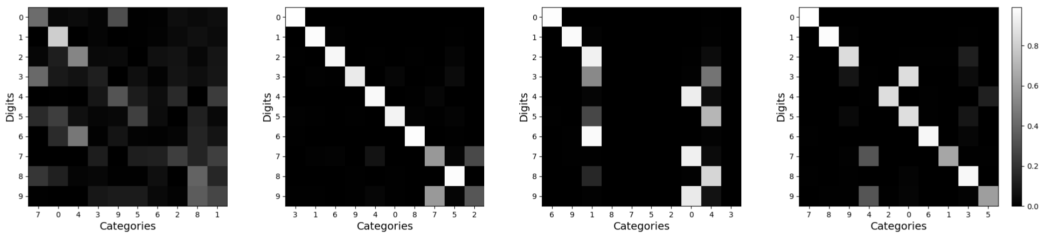


(b) JointVAE($[0, 1]|25K$) (c) MMIDAS(0.2) (d) JointVAE($[0, 25]|25K$) (e) MMIDAS(10)

Figure S5: **Sensitivity analysis.** (a) Effect of the consensus factor ($\lambda$) in MMIDAS and the channel capacity ($\mathcal{C}_s$) in the JointVAE models. Reported values present the average accuracy of categorical assignment for 3 randomly initialized runs, over $15K$ training iterations for the MNIST dataset. (b-c) Confusion matrices for JointVAE and MMIDAS models, respectively corresponding to the hyperparameters marked by the dash-dotted black line. (d-e) Confusion matrices for JointVAE and MMIDAS models, respectively corresponding to the hyperparameters marked by the dash-dotted red line.

### Supplementary Note 9

#### Comparative analysis of clustering approaches for cell type identification

To demonstrate the robustness and effectiveness of MMIDAS in inferring accurate and interpretable categorical variables, we conducted a comparative analysis. In this analysis, we evaluated the performance of MMIDAS in inferring categorical variables (cluster labels) with respect to that of single-cell clustering methods and leading-edge deep mixture models. The study utilized the same gene expression matrix, incorporating $\log(CPM + 1)$ normalized values (Methods), with the exception of scVI training, which required count values. For the community detection-based algorithms, the resolution parameter was fine-tuned to achieve optimal performance (based on the classification accuracy and Silhouette score). Training parameters for deep mixture models also correspond to the best performance that we obtained with those models.

**PCA + Seurat:** Utilizing PCA for dimension reduction with 100 PCs, creating a neighborhood graph with $k = 10$ neighbors for $k$-nearest neighbors (KNN), and subsequently applying the Leiden algorithm to the constructed graph.

**scVI + Seurat:** Utilizing scVI model for dimension reduction with 10 latent variables, building a neighborhood graph with $k = 10$ neighbors for KNN, and then applying the Leiden algorithm to the resulting graph.

**ACTIONet:** Following the tutorial provided with the associated code base, we initially performed kernel reduction on the normalized expression matrix (log-transformation). Subsequently, ACTIONet was executed to construct a structural representation of the cell-state patterns (encoding matrix $H^*$ as described in [15]) and to capture the cell–cell similarity graph. The Leiden algorithm was then applied to the similarity graph to identify clusters. Throughout these steps, we used the suggested default settings. In the archetype pruning part, we set the *minimum number of cells* parameter to 10.

**Deep mixture models:** We also conducted training sessions for JointVAE and CascadeVAE, two closely related unsupervised VAE-based mixture models that can be considered as 1-arm MMIDAS implementations. For the JointVAE and CascadeVAE models, we used the same network architecture used for MMIDAS (see Fig. S21b and Supplementary Note 14). In contrast to MMIDAS, which employs a pruning algorithm to dynamically adjust the dimensionality of the discrete space (Methods), mixture models exclusively learn discrete representations for a predefined set of categorical variables. Consequently, in all training sessions, we followed the suggested number of cell types, as specified in the taxonomy (Figs. S14, S16, and S17), as the dimensionality of the discrete space. Training parameters used for scRNA-seq datasets are listed as follows.

- **JointVAE** (Mouse Smart-seq)
  - Continuous and categorical variational factors: $D_s = 2$, $D_c = 115$
  - Batch size: 5000
  - Training epochs: $10K$
  - Gumbel-Softmax temperature ($\tau$): 1.0
  - Hyperparameters of KL divergence ($\gamma_{\mathbf{s}}$, $\gamma_{\mathbf{c}}$): 100
  - Continuous channel capacities ($C_{\mathbf{s}}$): Increased linearly from 0 to 50 in 10000 iterations
  - Discrete channel capacities ($C_{\mathbf{c}}$): Increased linearly from 0 to 10 in 10000 iterations
  - Optimizer: Adam with learning rate 1e-3
- **JointVAE** (Mouse 10x)
  - Continuous and categorical variational factors: $D_s = 3$, $D_c = 113$ (Glutamatergic), $D_c = 97$ (GABAergic)
  - Batch size: 5000
  - Training epochs: $5K$

  - Gumbel-Softmax temperature ($\tau$): 1.0
  - Hyperparameters of KL divergence ($\gamma_{\mathbf{s}}$, $\gamma_{\mathbf{c}}$): 100
  - Continuous channel capacities ($C_{\mathbf{s}}$): Increased linearly from 0 to 50 in 20000 iterations
  - Discrete channel capacities ($C_{\mathbf{c}}$): Increased linearly from 0 to 15 in 20000 iterations
  - Optimizer: Adam with learning rate 1e-3

- **CascadeVAE** (Mouse Smart-seq)
  - Continuous and categorical variational factors: $D_s = 2$, $D_c = 115$
  - Batch size: 5000
  - Training epochs: $10K$
  - Hyperparameter of $D_{KL}\big(q(\mathbf{c}|\mathbf{x}) \parallel U(|\mathbf{c}|)\big)$ ($\lambda'$): 0.1
  - Hyperparameter of $D_{KL}\big(q(\mathbf{s}|\mathbf{x}) \parallel p(\mathbf{s})\big)$ ($\beta$): Increased linearly from 0 to 10 in 10000 iterations
  - Optimizer: Adam with learning rate 1e-4

- **CascadeVAE** (Mouse 10x)
  - Continuous and categorical variational factors: $D_s = 3$, $D_c = 113$ (Glutamatergic), $D_c = 97$ (GABAergic)
  - Batch size: 5000
  - Training epochs: 1000
  - Hyperparameter of $D_{KL}\big(q(\mathbf{c}|\mathbf{x}) \parallel U(|\mathbf{c}|)\big)$ ($\lambda'$): 0.1
  - Hyperparameter of $D_{KL}\big(q(\mathbf{s}|\mathbf{x}) \parallel p(\mathbf{s})\big)$ ($\beta$): Increased linearly from 0 to 10 in 20000 iterations
  - Optimizer: Adam with learning rate 1e-4

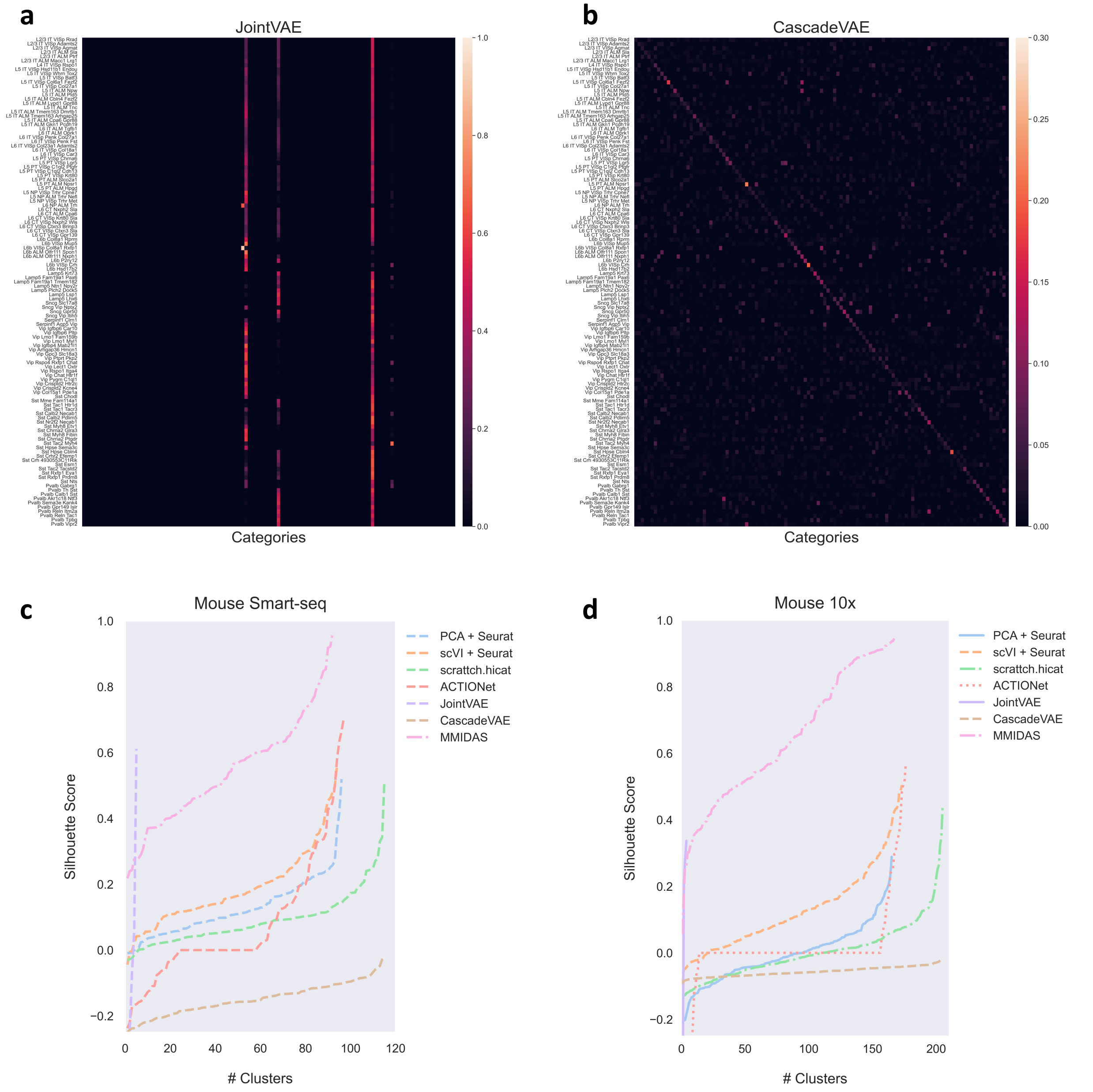


Figure S6: (a, b) Categorical variables derived using deep mixture models for the mouse Smart-seq dataset. (a) Mutual information scores between the 115 t-types identified by scrattch.hicat [18] (y-axis) and the 5 categories inferred by JointVAE (x-axis). (b) Mutual information scores between the same 115 t-types (y-axis) and the 115 categories inferred by CascadeVAE (x-axis). (c, d) Average Silhouette scores per-cluster, for the mouse Smart-seq dataset (c) and the 10x dataset (d).

### Supplementary Note 10

#### Dimensionality of the latent space

Efficient optimization of the dimensionality of the latent space in VAE-based models remains an open problem in model selection. In this manuscript, we introduce a pruning technique aimed at optimizing the dimensionality of the categorical latent factor, denoted as $|\mathbf{c}|$, within an over-parameterized simplex (Methods).

The choice of the dimensionality for the continuous latent space, $|\mathbf{s}|$, presents a more intricate problem. In the case of adult mouse single-cell datasets obtained from mice with homogeneous genetic backgrounds, a low-dimensional latent space, such as 2- or 3-dimensional space, typically suffices to capture both technical and biological variabilities [7, 17, 6]. In contrast, for the human AD dataset, we opt for a higher-dimensional latent space, $|\mathbf{s}| = 15$, owing to increased variability arising from diverse donor conditions (e.g., disease stage, cognitive scores, age, race). While computationally expensive, model selection can be performed for $|\mathbf{s}|$ by gradually increasing this dimensionality and monitoring its impact on the categorical assignment and the clusterability metric.

### Supplementary Note 11

#### Number of arms

Our theoretical results show that the required number of arms is a function of the categorical distribution and the likelihood term (Supplementary Note 3, Eq. S39). In the case of uniformly distributed categories, as exemplified by datasets like MNIST, we demonstrate that just a single pair of coupled arms ($A = 2$) is sufficient for MMIDAS to accurately infer the underlying categorical representation (as indicated in Corollary1). However, in more general scenarios where the distribution of clusters remains unknown, and their abundances differ significantly, the minimum requisite number of arms remains uncertain. In single-cell data analysis, despite this uncertainty, we observe that even with just 2 arms, the model can infer a reproducible and distinguishable discrete representation of the underlying data as quantified by the classification performance and the average Silhouette score. To further investigate the impact of additional arms on the categorical representation, we conduct experiments with different numbers of arms. Table S3 summarizes MMIDAS's clusterability performance on the mouse Smart-seq dataset as a function of the number VAE arms. While increasing the number of arms does indeed enhance categorical performance, the improvements are relatively small. Table S3 also highlights the trade-off between computational cost and performance enhancement when choosing the number of arms.

| $A$ (number of arms) | **ACC** (%) ↑ (mean ± s.d.) | **Silhouette score** ↑ (mean ± s.d.) | **Computation** ↑ (iterations/sec) |
|---|---|---|---|
| 2 | 98.38 ± 0.30 | 0.55 ± 0.16 | 2.5 |
| 3 | 98.79 ± 0.24 | 0.60 ± 0.16 | 1.8 |
| 4 | 99.27 ± 0.27 | 0.71 ± 0.19 | 1.4 |
| 5 | 99.46 ± 0.16 | 0.76 ± 0.10 | 1.2 |

Table S3: Categorical representation learning performance of MMIDAS as a function of the number of VAE arms ($A$) for the mouse Smart-seq dataset. "ACC" denotes the balanced classification accuracy attained on test cells for all 92 consensus transcriptomic categories inferred by MMIDAS. Categorical assignments of each multi-arm configuration is used to train a Random Forest classifier. "Silhouette score" reports the mean and standard deviation of the per-cluster Silhouette score across class labels. "Computation" indicates the average number of iterations per second achieved on a GeForce RTX 2080 Ti GPU with a batch size of 5000.

### Supplementary Note 12

#### Impact of the network depth on mixture modeling performance

We compare the performance of a shallower network with one hidden layer (shallow MMIDAS) to a deeper network with four hidden layers on the Smart-seq dataset to elucidate the impact of deeper processing. To facilitate the comparison, training is performed without the pruning step, allowing both models to have the same dimensionality of discrete and continuous variables. The results in Fig.S7 and Table S4 indicate that deeper processing improves the clustering and reconstruction performances. While both metrics improve statistically significantly (based on Gaussian models of the studied metrics), the bigger impact of deeper processing is on the accuracy of the generative model, rather than clusterability.

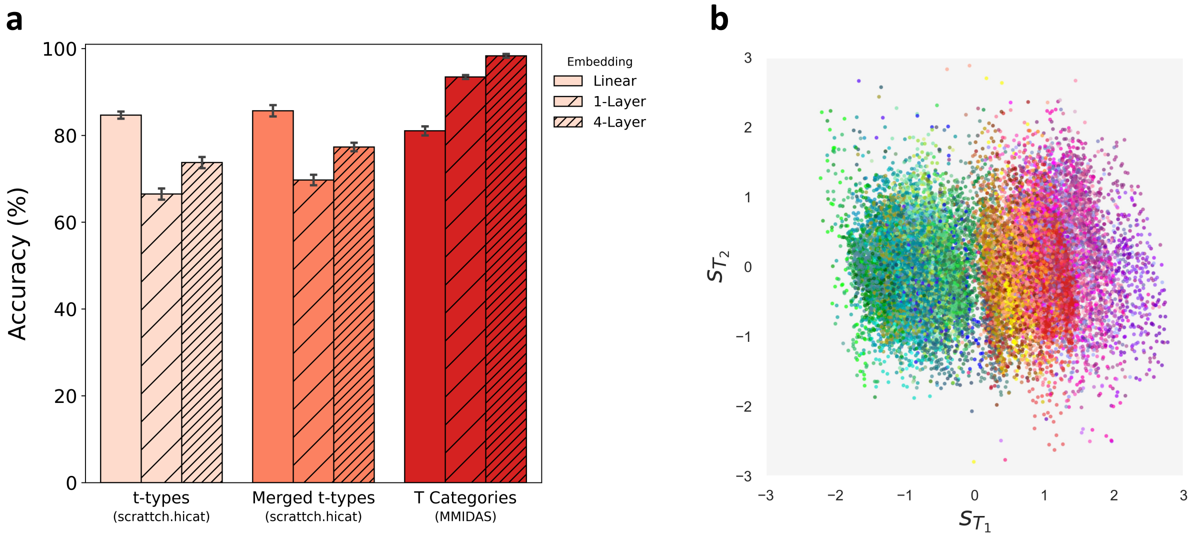


Figure S7: **Impact of network depth on MMIDAS' performance** (a) The discrete T categories obtained by the shallow model (MMIDAS with 1 layer of non-linearity) are compared with those from the deeper model (MMIDAS with 4 layers of non-linearity) for the mouse Smart-seq dataset. The average balanced classification accuracy with standard deviation is reported for 10-fold cross-validation, evaluated across both linear (100-dimensional) and non-linear (10-dimensional $\mathbf{z}$ in the shallow MMIDAS) embeddings. A Random Forest classifier is used for the classification task. (b) The 2-dimensional latent representation of continuous variability, color-coded by the inferred 92 T categories obtained by the shallow network. The representation learned by the shallow network does not properly encode the continuous type-dependent variability, as compared to Fig. 2e.

| **Dataset** | **MMIDAS's Network** | $\mathcal{L}_{\text{rec}} \downarrow$ (mean ± s.d.) | $\rho_{\text{rec}} \uparrow$ (mean ± s.d.) | $\Delta_{\text{rec}} \downarrow$ (mean ± s.d.) | **ACC** (%) ↑ (mean ± s.d.) | **Silhouette Score** ↑ (mean ± s.d.) |
|---|---|---|---|---|---|---|
| Mouse Smart-seq | 1-Layer | 1.89 ± 0.06 | 0.97 ± 0.00 | 0.86 ± 0.09 | 93.45 ± 0.59 | 0.26 ± 0.14 |
| | 4-Layer | 1.57 ± 0.03 | 0.99 ± 0.00 | 0.29 ± 0.05 | 98.38 ± 0.30 | 0.55 ± 0.16 |
| Mouse 10x | 1-Layer | 8.74 ± 1.32 | 0.48 ± 0.07 | 2.53 ± 1.07 | 96.02 ± 1.85 | 0.31 ± 0.25 |
| | 4-Layer | 4.96 ± 0.50 | 0.81 ± 0.02 | 0.95 ± 0.99 | 99.48 ± 0.27 | 0.68 ± 0.14 |

Table S4: The performance of the mixture representation obtained by MMIDAS is evaluated as a function of network depth for both the mouse Smart-seq with 5,032 genes and 10x datasets with 10,000 genes. The average reconstruction errors for the normalized gene expression values are denoted by $\mathcal{L}_{\text{rec}}$, calculated for the test set. $\rho_{\text{rec}}$ and $\Delta_{\text{rec}}$ refer to the correlation and mean-squared error between reconstructed gene expression across arms, respectively. "ACC" refers to the average balanced classification accuracy, including the standard deviation, assessed using a Random Forest classifier across non-linear (10-dimensional $\mathbf{z}$ in each network) embeddings. "Silhouette Score" indicates the mean and standard deviation of the per-cluster Silhouette score. All reported metrics are averaged across arms.

### Supplementary Note 13

#### Additional insights into c and s inferred by MMIDAS

To gain a deeper understanding on the inferred discrete and continuous representations for single-cell measurements, we study three potential scenarios:

1. t-types pairs that MMIDAS merges into a cluster,
2. t-types that MMIDAS splits into multiple clusters,
3. t-types that are essentially preserved by MMIDAS.

We choose the cell types for each scenario based on mutual information scores shown in Fig. 2a, focusing on the glutamatergic types in the mouse Smart-seq dataset.

To assess the biological significance of the grouped cells, we study the significance of the difference between clusters based on the number of genes that are statistically significantly differentially expressed (DE). We used the `scanpy.tl.rank_genes_groups` function with the Wilcoxon method for statistical hypothesis testing. At a $p$-value of $p < 10^{-20}$, we find that, on average, 60 genes are differentially expressed between neighboring glutamatergic t-type pairs, i.e. denoted as $\bar{n}_{DE}$ in Fig. S8a.

- In cases belonging to the first scenario above (t-types merged by MMIDAS), for instance L2/3 IT ALM Sla vs. L2/3 IT ALM Ptrf (Fig. S8a), we find that the number of the DE genes is much fewer than the average. This suggests that, in comparison, MMIDAS merges subpopulations that are not significantly differentiated from each other.
- In cases belonging to the second scenario above (t-types split by MMIDAS), e.g. L2/3 IT VISp Agmat (Fig. S8a), we find that the number of the DE genes between MMIDAS clusters is above the average. This suggests that, in comparison, MMIDAS splits populations with enough differentiation to support a discrete perspective into subpopulations.
- In cases belonging to the third scenario above (t-types preserved by MMIDAS), for instance L6 IT ALM Tgfb1 and L6 IT ALM Oprk1 (Fig. S8a), we find that the number of the DE genes between the t-type and its neighbors are above the average, supporting the discretization.

Thus, MMIDAS demonstrates an ability to identify “sufficiently differentiated” sub-populations, and offers revisions to previously under-, and over-split categorizations.

Next, we explore the continuous variability inferred by MMIDAS within example categories corresponding to the different scenarios mentioned above:

- Fig. S8b displays a unified continuum within the **s** space for the L2/3 IT T_6 category, merging three t-types in Fig.S8a. This visualization underscores that not only is there an absence of enough DE genes between these t-types, but also that the variability across them can be adequately accounted for by the continuous variable. Moreover, there is no clear separation observed among cells derived from different t-types.
- In contrast, T categories L2/3 IT T_3, L2/3 IT T_4, and L2/3 IT T_5, which were considered as a single t-type (L2/3 IT VISp Agmat in Fig.S8a), separate into distinct substructures in the **s** space. Moreover, the inferred continuous variabilities within each T category also appear heterogeneous.
- Fig. S8d shows a single continuum in the **s** space for the T category L5 PT T_27, which aligns well with the L5 PT VISp Chrna6 t-type. This provides further evidence for MMIDAS’ preservation of this t-type: not only are there enough DE genes between the t-type and its neighbors, the variability within this population is also well explained by a continuum.

In addition to examining the latent continuous representation, we also compare the overall gene expression patterns for two merge and split scenarios in Figs. S8b,c.

- Fig. S8c displays the average expression level of all genes for each cell, across three t-types that were grouped by MMIDAS and identified as a single T category, i.e. L2/3 IT T_6. This figure illustrates that the variabilities observed are more akin to a continuum rather than discrete changes among t-types.
- In contrast, Fig. S8d reveals discrete changes (jumps) in gene expression among cells within the L2/3 IT VISp Agmat t-type. Although the reference taxonomy assigns only one type to these cells, MMIDAS appropriately separates them, as indicated by the observed jumps in average gene expression.

Thus, MMIDAS consistently uses the discrete and continuous factors of variability and jointly infers them for a unified and improved understanding of the landscape of neuronal phenotypes.

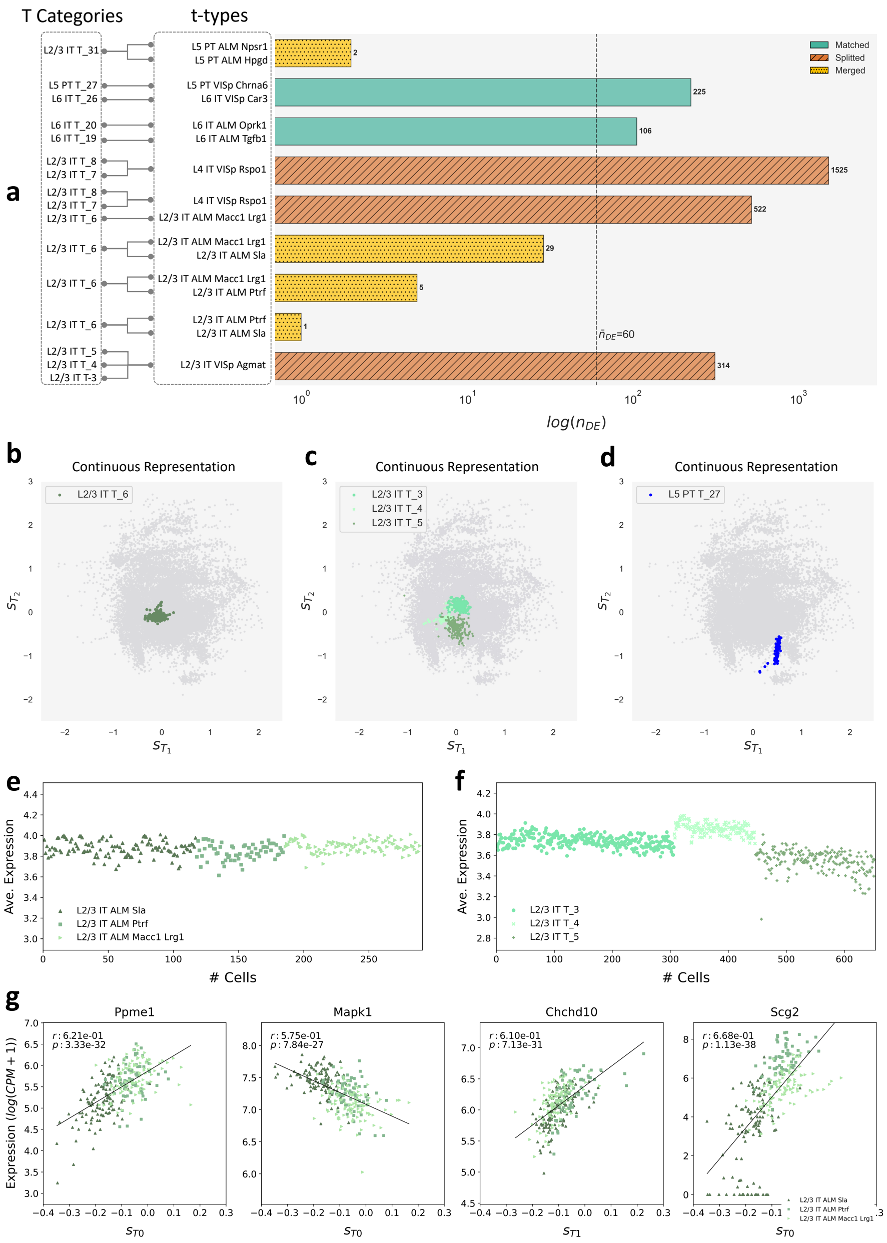

Figure S8: **Exploring the differences between MMIDAS categories and t-types.** (a) Differential expression analysis is performed for glutamatergic cells in the mouse Smart-seq dataset. The bar plot illustrates the number of DE genes across 11 t-types as defined in [18], alongside their corresponding 11 T categories obtained by MMIDAS. Three distinct relationship patterns emerge between t-types and T categories: i) merging pattern, denoted by dots, where multiple t-types are merged into a single T category; ii) splitting pattern, denoted by dashed lines, where a single t-type is split into multiple T categories; and iii) matching pattern, where one t-type corresponds to a single T category. $\bar{n}_{DE}$ denotes the average number of DE genes between neighboring glutamatergic t-type pairs. Statistical significance is determined using the Wilcoxon test with a threshold of $p < 10^{-20}$. Corresponding continuous representations for (b) a T category merging two t-types, (c) three T categories splitting a single t-type, and (d) a T category predominantly aligning with a single t-type. Gray-colored dots represent all cells underlying the analysis. (e) Average gene expression for each cell within three t-types including L2/3 IT ALM Sla, L2/3 IT ALM Ptrf, and L2/3 IT ALM Macc1 Lrg1, which have been merged and treated as a single T category, as illustrated in (a) and (b). (f) Average gene expression per cell across three T categories, which are annotated as a single t-type, as illustrated in (a) and (c). (g) The normalized expression levels of a subset of genes exhibit monotonic changes in relation to continuous type-dependent variables within the L2/3 IT T-6 cluster, which includes three t-types: L2/3 IT ALM Sla, L2/3 IT ALM Ptrf, and L2/3 IT ALM Macc1 Lrg1.

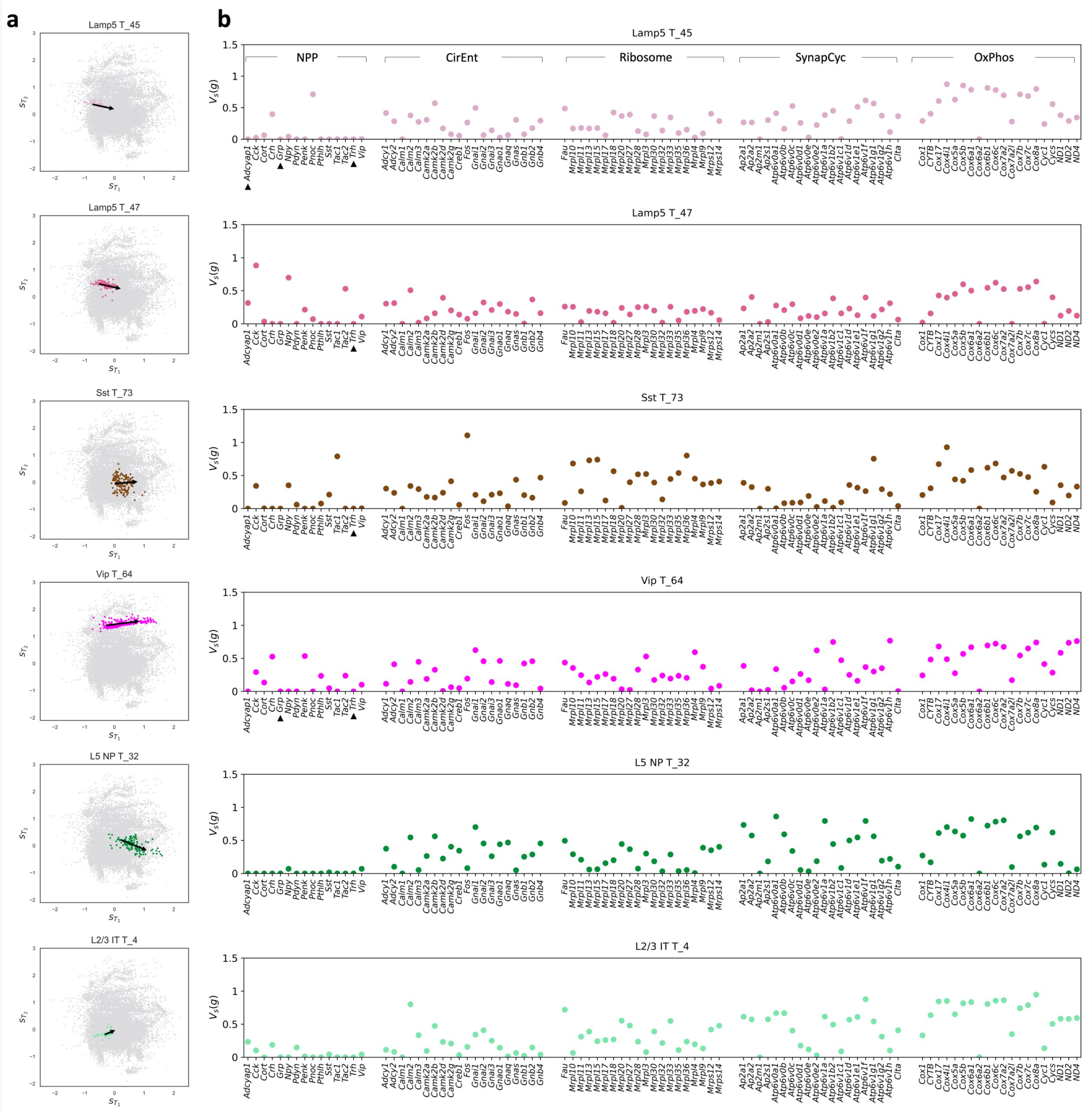


Figure S9: **Continuous traversal analysis in the adult mouse, based on data from the Smart-seq platform.** (a) 2-dimensional representation of the inferred continuous variables for six MMIDAS T-categories. Each point represents a cell. For each category, the black arrow denotes the eigenvector with the largest magnitude eigenvalue capturing the primary axis of continuous variation. (b) Average normalized variation (Eq. 16) over each cell type for a select group of genes involved in five signaling pathways identified in [11]. Among these pathways, The NPP set includes 15 neuropeptide precursor genes involved in neuropeptide signaling pathways [17]. In CirEnt set includes 20 (randomly selected) genes associated with circadian entrainment pathways. The Ribosome set includes 20 (randomly selected) genes associated with neuronal ribosomal protein regulation. The SynapCyc set includes 20 (randomly selected) genes associated with the synaptic vesicle cycle. OxPhos includes 20 (randomly selected) genes associated with the oxidative phosphorylation pathway. Each data point on the plot denotes the variation of the expression value of an individual gene as **s** traverses along the dominant eigenvector within the continuous space. Genes with cell type–specific expression profiles, particularly those in NPP [17], exhibit comparatively smaller variations. Conversely, genes contributing to cellular activity, environmental adaptation, and metabolism, display larger variations. Genes with zero expression for all samples in the category are marked with ▲.

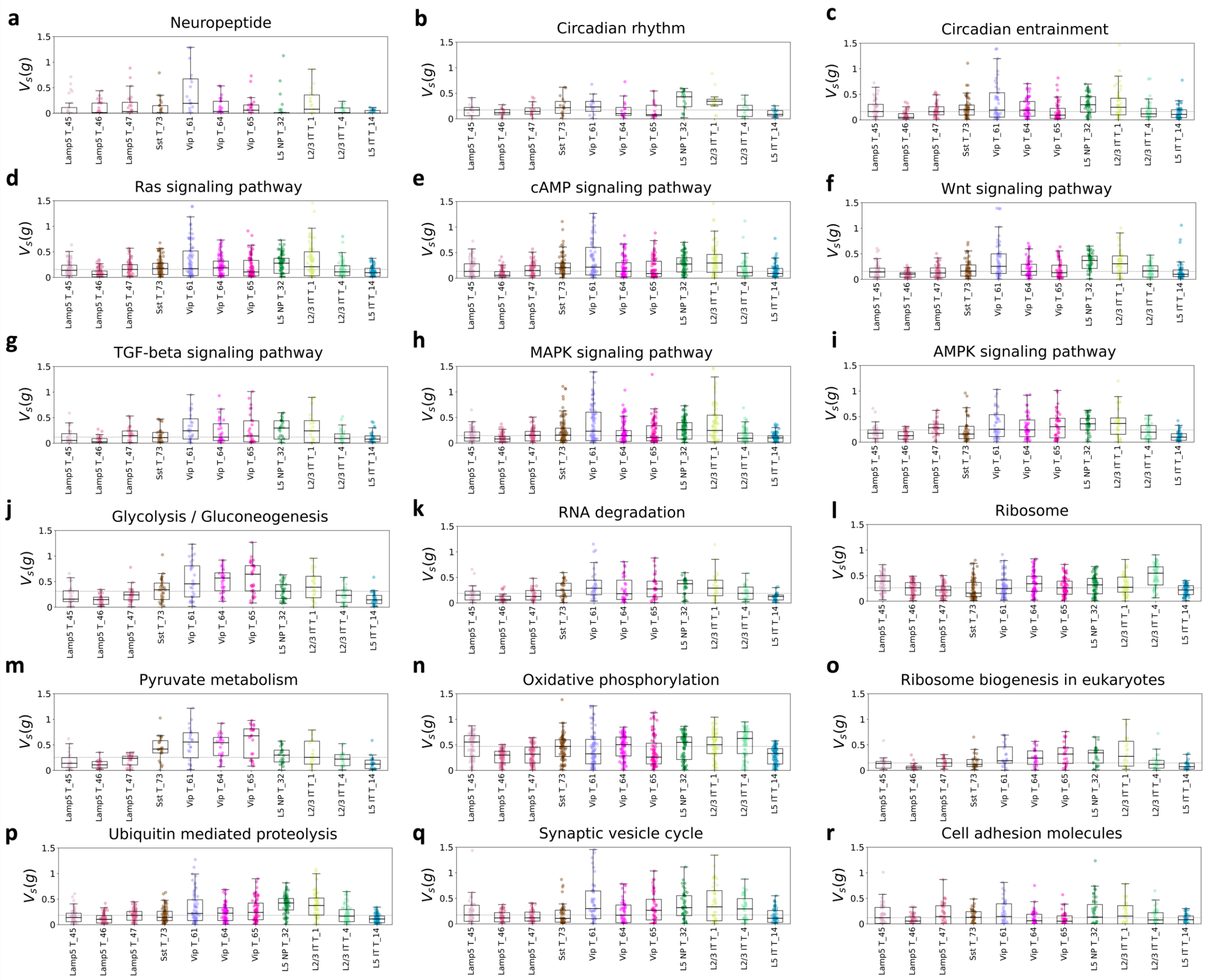


Figure S10: **Exploring the continuous variability across cell types for a set of cellular pathways in the adult mouse Smart-seq dataset.** In each subfigure, the expression variability of gene modules is showcased across 11 MMIDAS T categories. These gene modules are linked to distinct cellular pathways [11].

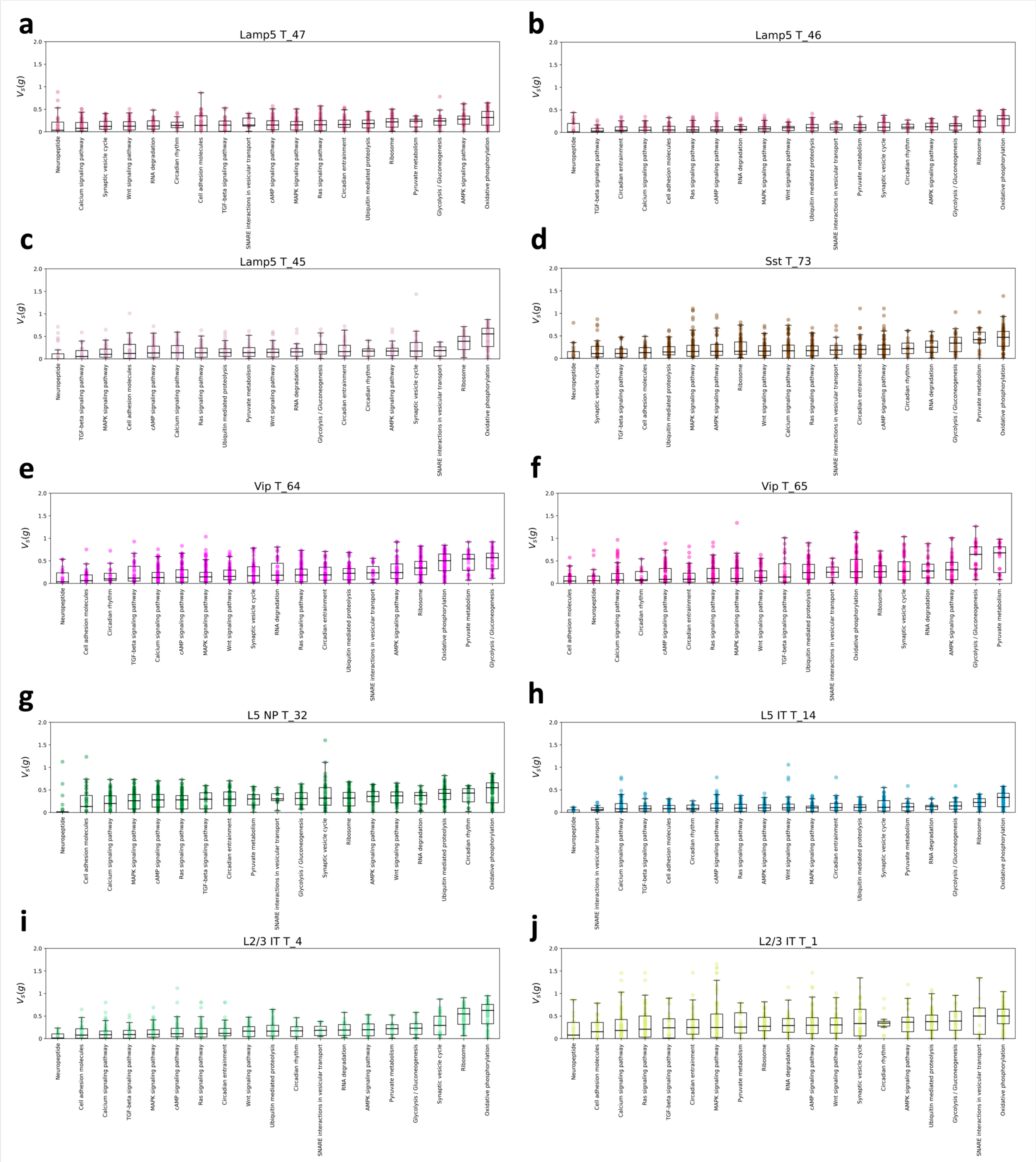


Figure S11: **Exploring the continuous variability across cellular pathways for individual cell types in the adult mouse Smart-seq dataset.** Each subfigure demonstrates the sorted expression variability across gene modules contributing to 19 cellular pathways [11] for a MMIDAS T category. Gene modules involved in neuropeptide signaling always exhibit the lowest variation under the continuous traversal analysis. In contrast, genes that contribute to oxidative phosphorylation pathway exhibit greater variation across all types.

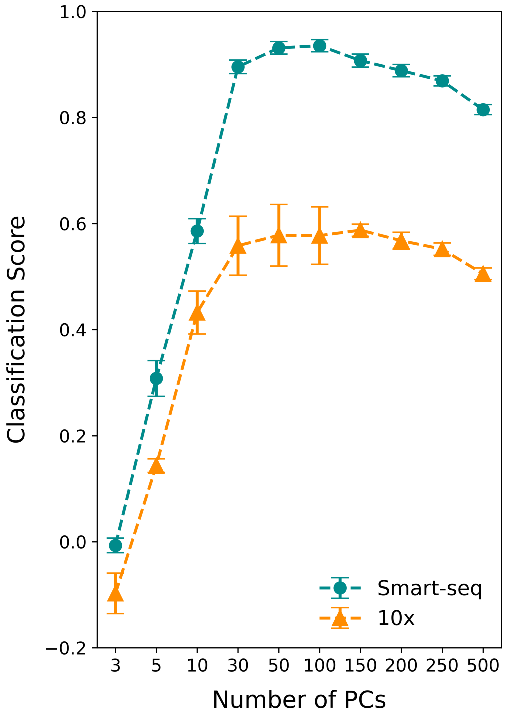


Figure S12: **Interplay between the number of principal components and t-type classification performance.** The figure shows the average classification scores for t-types obtained by the `scrattch.hicat` method, along with their standard deviations, obtained from a 10-fold cross-validation process on both Smart-seq and 10x mouse datasets. The classification score is calculated by adding the balanced classification accuracy and the overall silhouette scores for all t-types. A Random Forest classifier is used for the classification task. For enhanced clarity in visualization of the results, x-axis is not uniformly scaled.

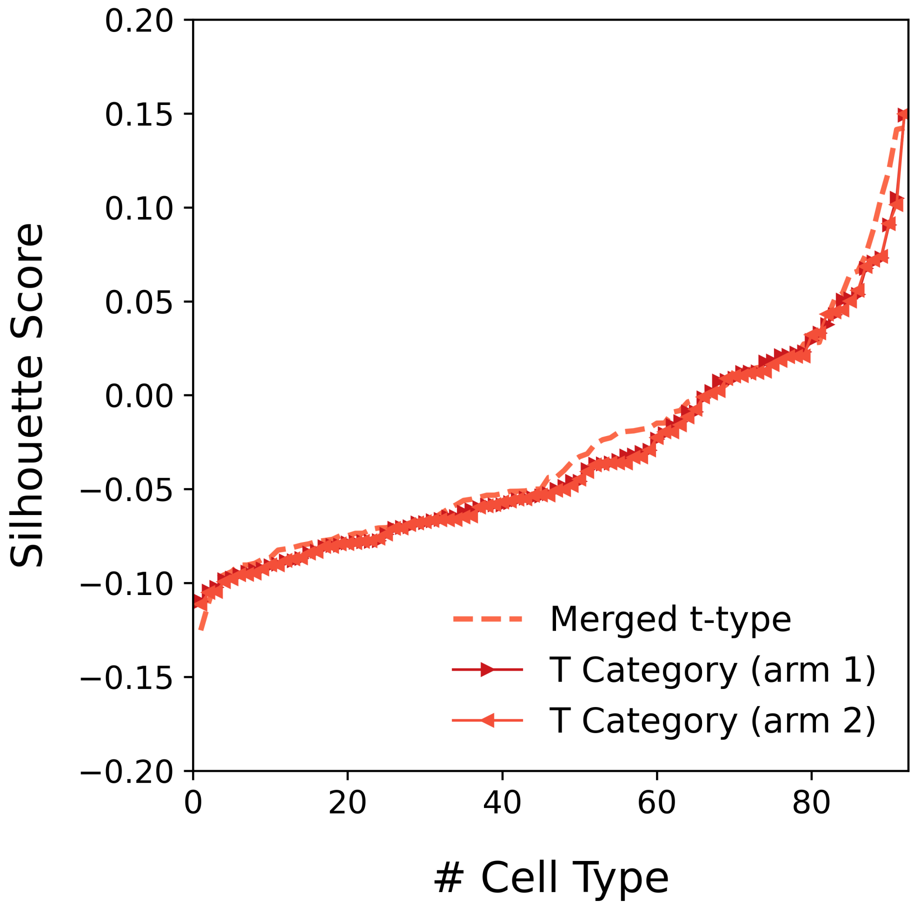


Figure S13: **Silhouette score in the high-dimensional gene space.** Average Silhouette scores per-cluster, calculated from all cells in the mouse Smart-seq dataset. The Silhouette coefficients are computed directly in the high-dimensional gene space, which are much lower (all less than zero on average) than the scores calculated on the lower dimensional embeddings (Fig. 2d, and Table 1), illustrating the "curse of dimensionality".

| Cell types | K | Embedding | | | |
|---|---|---|---|---|---|
| | | $\mathbf{z}_{linear}$ (100 PCs) | $\mathbf{z}_{non-linear}$ ($\|\mathbf{z}\| = 10$) | $\mathbf{z}^*_{linear}$ (100 PCs) | $\mathbf{z}^*_{non-linear}$ ($\|\mathbf{z}\| = 10$) |
| t-types | 115 | 84.7 ± 0.8 | 73.7 ± 1.6 | 84.0 ± 0.7 | 71.4 ± 2.1 |
| Merged t-types | 92 | 85.7 ± 1.2 | 77.3 ± 1.5 | 85.1 ± 1.4 | 75.2 ± 2.6 |
| MMIDAS categories | 92 | 88.5 ± 1.2 | 98.3 ± 0.5 | 87.3 ± 1.0 | 98.2 ± 0.4 |

Table S5: Performance of the Random Forest (RF) classifier for three groups of cell type labels using linear and non-linear embeddings. A 100-dimensional linear embedding, denoted as $\mathbf{z}_{linear}$, is obtained from gene expression data across the entire dataset. A 10-dimensional non-linear embedding, denoted as $\mathbf{z}_{non-linear}$, is obtained using the entire dataset. Embeddings $\mathbf{z}^*_{linear}$ and $\mathbf{z}^*_{non-linear}$ are obtained within each cross validation fold from the training data. Classifier performances are similar between the $\mathbf{z}$ and $\mathbf{z}^*$ variants.

| Datasets | Cell types | | | | | | | | |
|---|---|---|---|---|---|---|---|---|---|
| | t-types | | | Merged t-types | | | MMIDAS categories | | |
| | K | RF | KNN | K | RF | KNN | K | RF | KNN |
| Mouse Smart-seq | 115 | 84.7 ± 0.8 | 83.5 ± 1.0 | 92 | 85.7 ± 1.2 | 85.1 ± 0.9 | 92 | 98.2 ± 0.5 | 98.1 ± 0.5 |
| Mouse 10x | 205 | 55.5 ± 5.6 | 61.0 ± 2.5 | 167 | 59.8 ± 8.9 | 65.2 ± 4.8 | 167 | 99.5 ± 0.3 | 99.4 ± 0.4 |
| Mouse Patch-seq (T) | 92 | 59.2 ± 6.2 | 65.1 ± 8.4 | 60 | 64.5 ± 6.5 | 72.9 ± 8.0 | 60 | 91.2 ± 2.9 | 91.1 ± 2.9 |
| Mouse Patch-seq (E) | 92 | 40.0 ± 3.6 | 38.7 ± 3.5 | 60 | 47.5 ± 3.6 | 46.4 ± 3.6 | 60 | 89.7 ± 1.2 | 89.0 ± 1.5 |
| SEA-AD | 42 | 81.1 ± 1.4 | 75.0 ± 1.1 | 30 | 85.0 ± 1.2 | 82.6 ± 1.0 | 30 | 97.7 ± 0.2 | 92.4 ± 1.0 |

Table S6: Performance comparison of the Random Forest (RF) classifier versus the K-nearest neighbors (KNN) classifier using class labels obtained by MMIDAS and scrattch.hicat (t-types and Merged t-types) across the four reported datasets. $K$ represents the number of identified cell types. Average balanced accuracy with standard deviation on test samples is reported for 10-fold cross-validation. MMIDAS consistently outperforms scrattch.hicat in both metrics and across datasets, with similar performances observed between the classifiers.

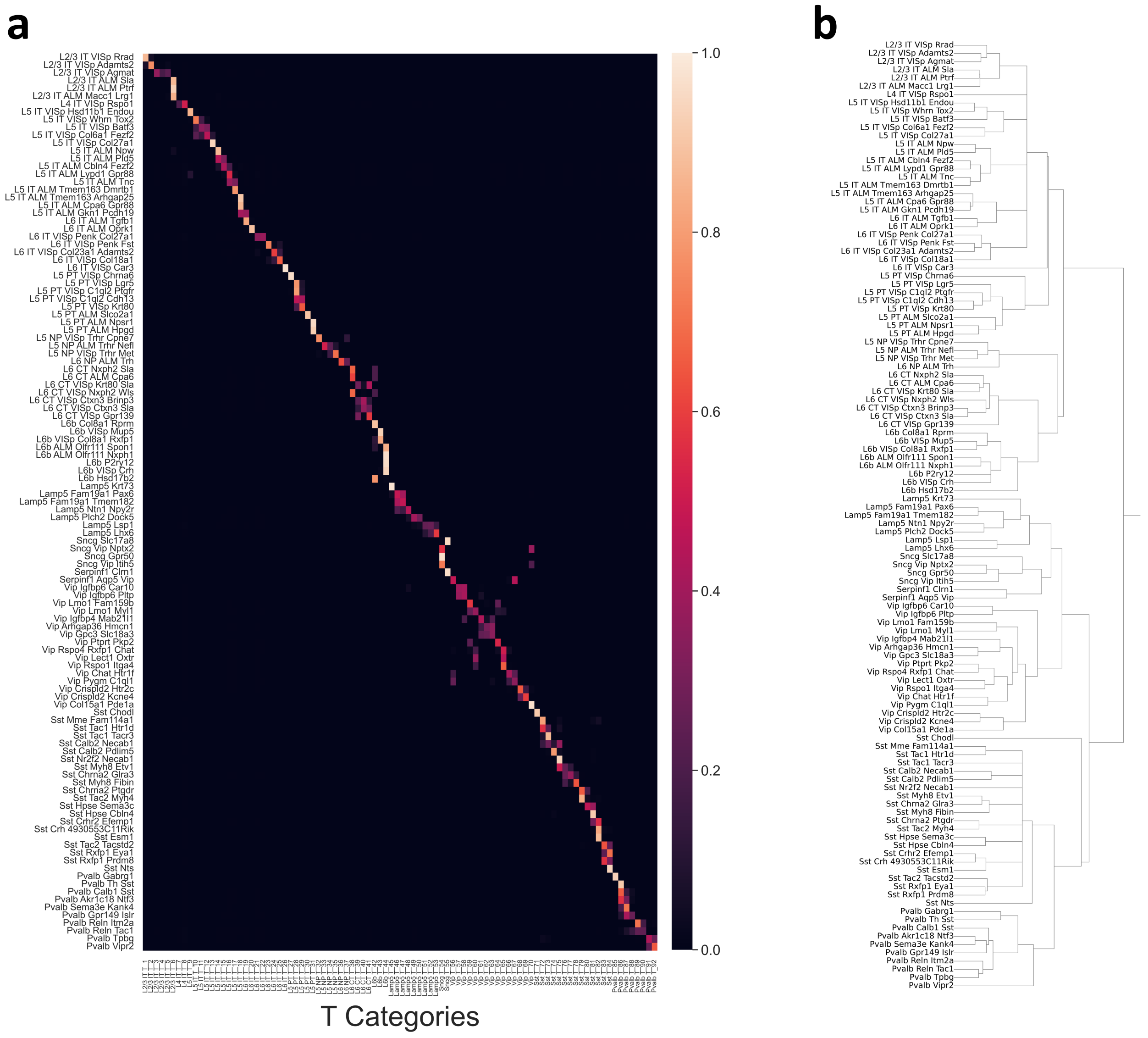


Figure S14: **Matching of inferred MMIDAS categories and reference t-types for the mouse Smart-seq dataset.** (a) Mutual information scores between 115 t-types (y-axis) and 92 MMIDAS categories (x-axis). (b) Reference taxonomies for 115 t-types, including 55 glutamatergic and 60 GABAergic distinct neuronal types reported as a taxonomy in [18] for the mouse Smart-seq dataset. Taxonomy is built by filtering out three categories of t-types: non-neuronal types, t-types with less than 10 cells, and those belonging to "CR" and "Meis2" subclasses.

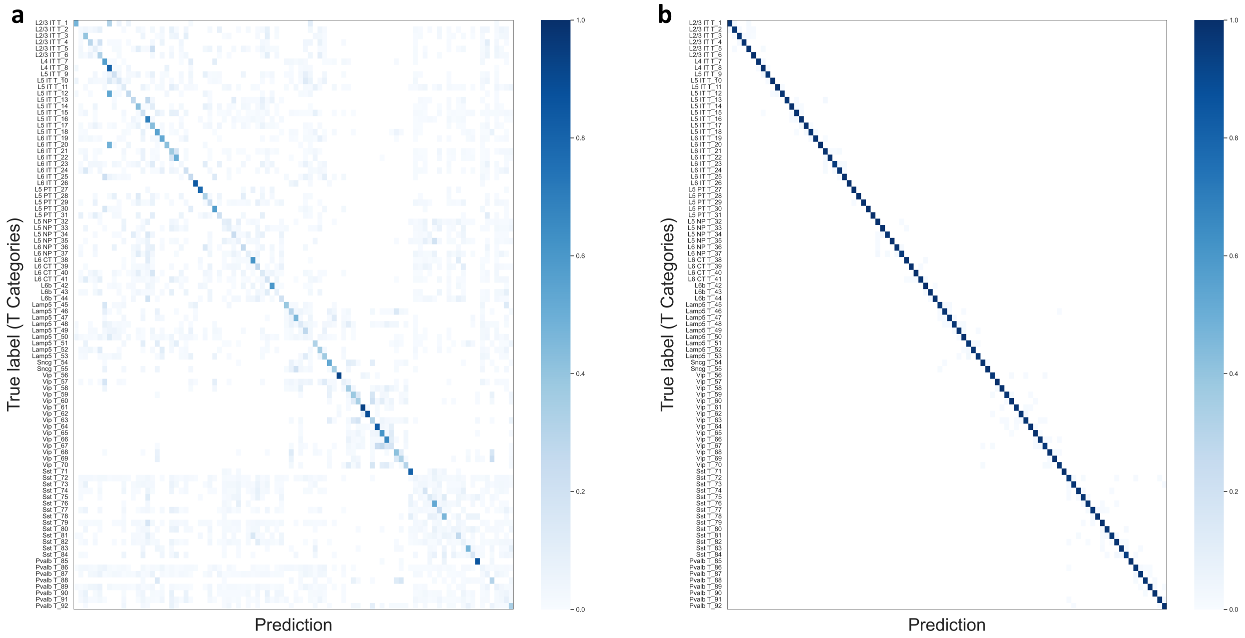


Figure S15: **Predicting discrete cell types using inferred continuous type-dependent variables versus inferred categorical variables.** (a) Confusion matrix from 10-fold cross-validation showing classification performance using the continuous latent variable **s** inferred by MMIDAS to predict 92 transcriptomic categories in the mouse Smart-seq dataset using a Random Forest classifier. (b) Same as (a) for the categorical latent variable **c** instead of **s**. Comparing these matrices reveals that the continuous, within-type factors of variability include some categorical information. They are a weak classifier of cell types. Here, chance level accuracy is 5%.

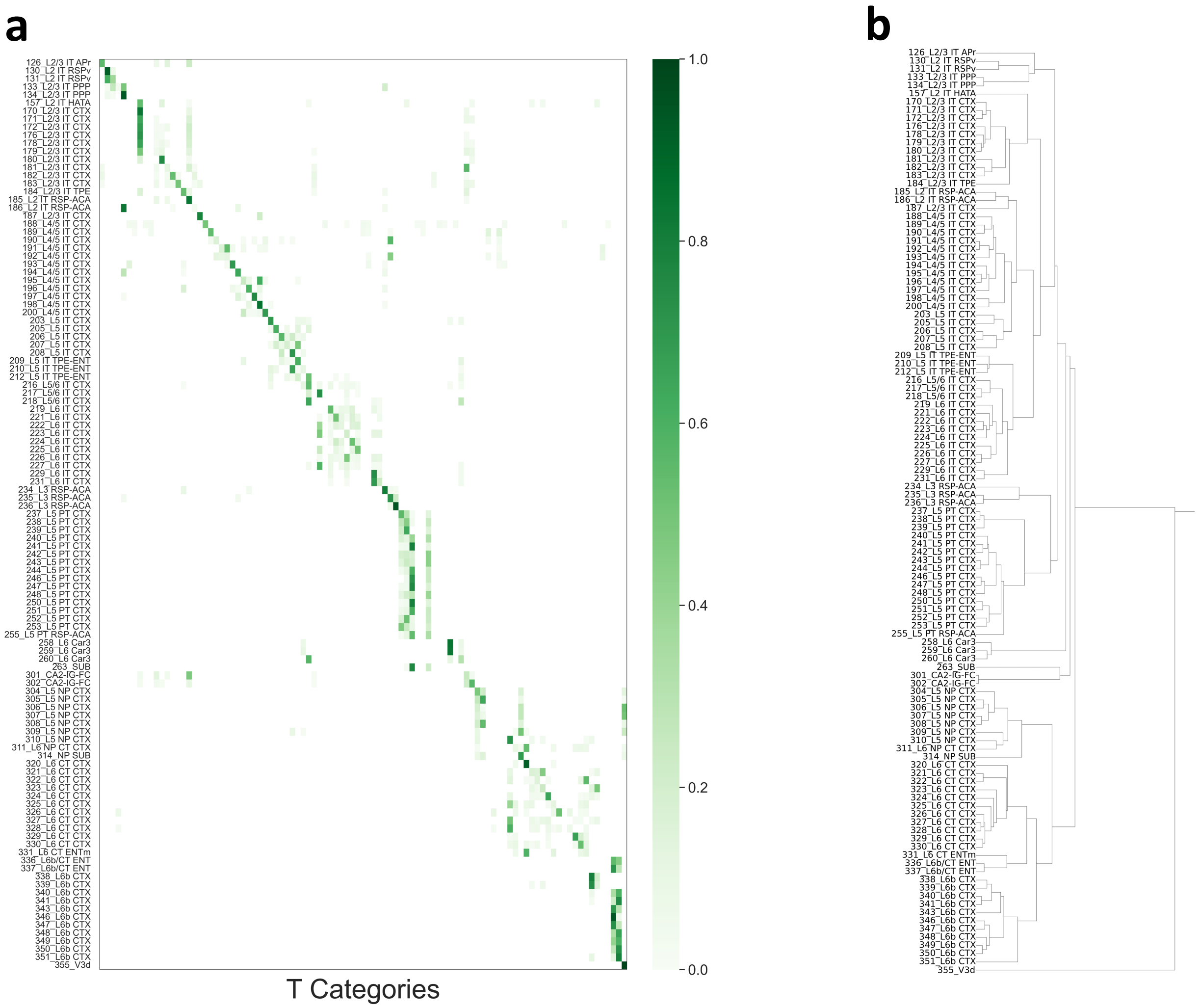


Figure S16: **Matching of inferred MMIDAS categories and reference t-types for glutamatergic cortical cells in the mouse 10x dataset.** (a) Mutual information scores between 97 MMIDAS categories (x-axis) and 113 t-types (y-axis). (b) Reference taxonomy for 113 t-types [20]. Here the taxonomy is built by filtering out two categories of t-types: t-types with less than 10 cells and those belonging to "CR" subclasses.

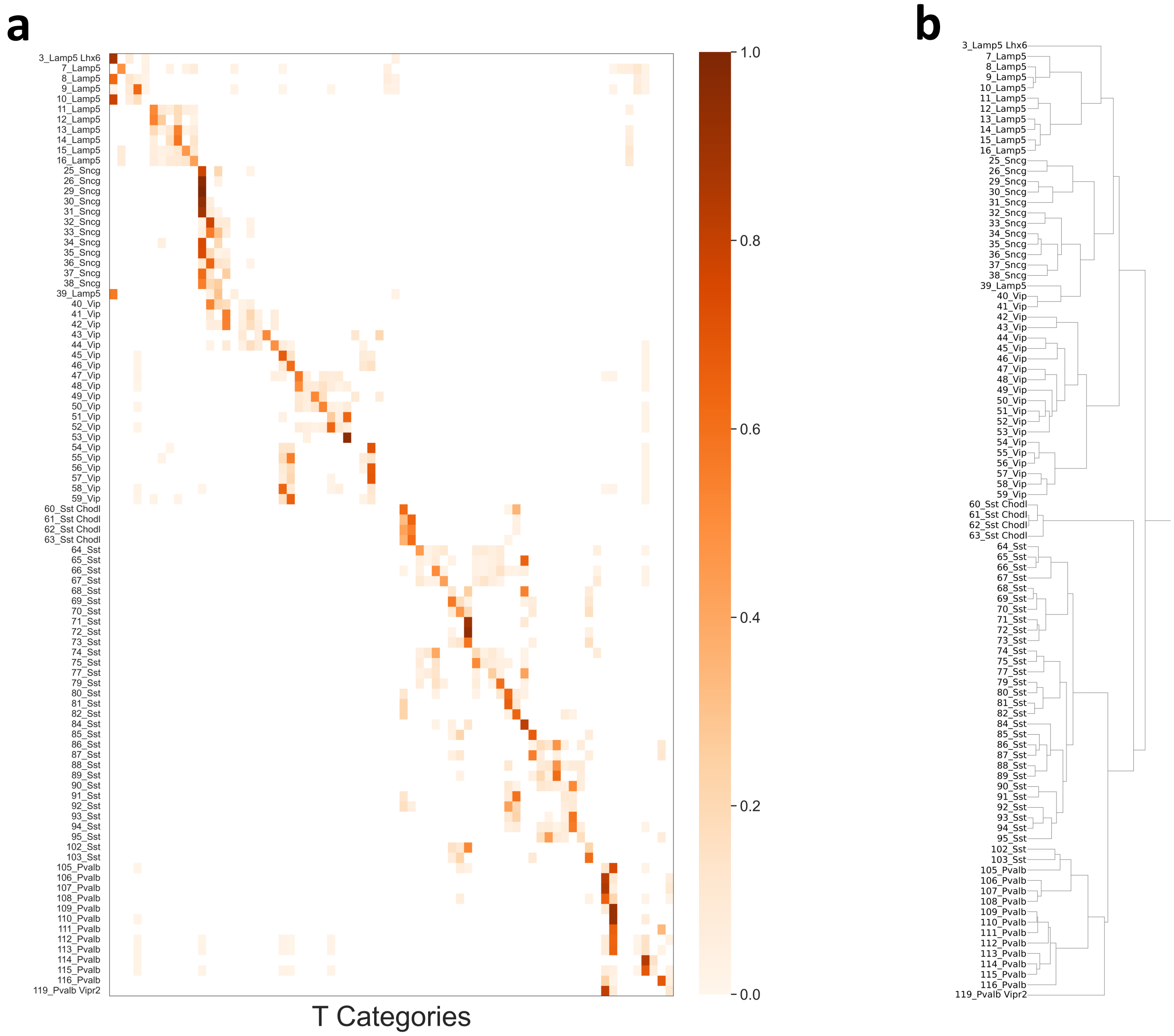


Figure S17: **Matching of inferred MMIDAS categories and reference t-types for GABAergic cortical cells in the mouse 10x dataset.** (a) Mutual information scores between 70 MMIDAS (x-axis) categories and 92 t-types (y-axis). (b) Reference taxonomy for 92 t-types in [20]. Here, the taxonomy is built by filtering out two categories of t-types: t-types with less than 10 cells and those belonging to "NP PPP", "Ndnf", "Meis2", and "Pax" subclasses.

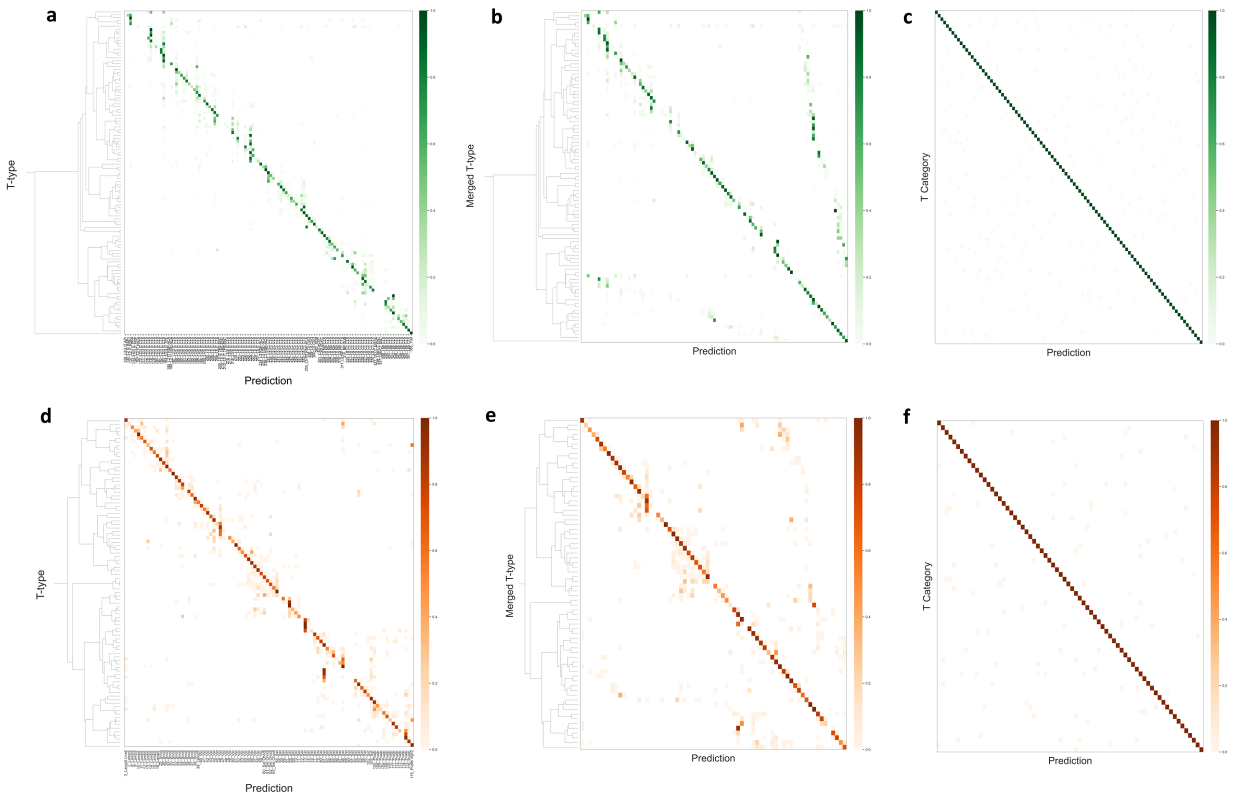


Figure S18: **Classification of transcriptomic cell types in the mouse cortex 10x data.** A Random Forest classifier is used in all plots. Dendrograms on the left sides of the plots illustrate the proposed hierarchical relationships between the cell types in the respective reference classifications. Confusion matrices displaying the classification performance for (a) 113 glutamatergic transcriptomic cell types as suggested in [20], (b) 97 merged glutamatergic transcriptomic cell types, merged according to the reference transcriptomic hierarchy [20], (c) 97 glutamatergic MMIDAS T categories, (d) 92 GABAergic transcriptomic cell types as suggested in [20]. (e) 70 merged GABAergic transcriptomic cell types, merged according to the reference transcriptomic hierarchy [20], (f) 70 GABAergic MMIDAS T categories.

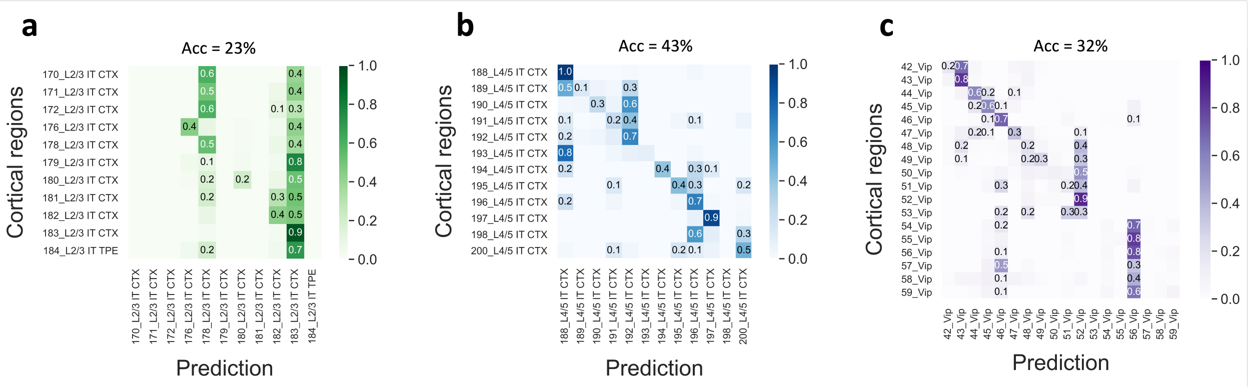


Figure S19: **Classification of transcriptomic cell types using the continuous representation inferred by MMIDAS.** Confusion matrices for predicting (a) L2/3 IT, (b) L4/5 IT, and (c) *Vip* neurons in the mouse 10x isocortex data are shown. A Random Forest classifier is used in all plots. "Acc" denotes the overall balanced accuracy of identifying t-types from the continuous representation. The classification performance indicates that the latent continuous representation does not encode discrete diversity for either of sub-classes.

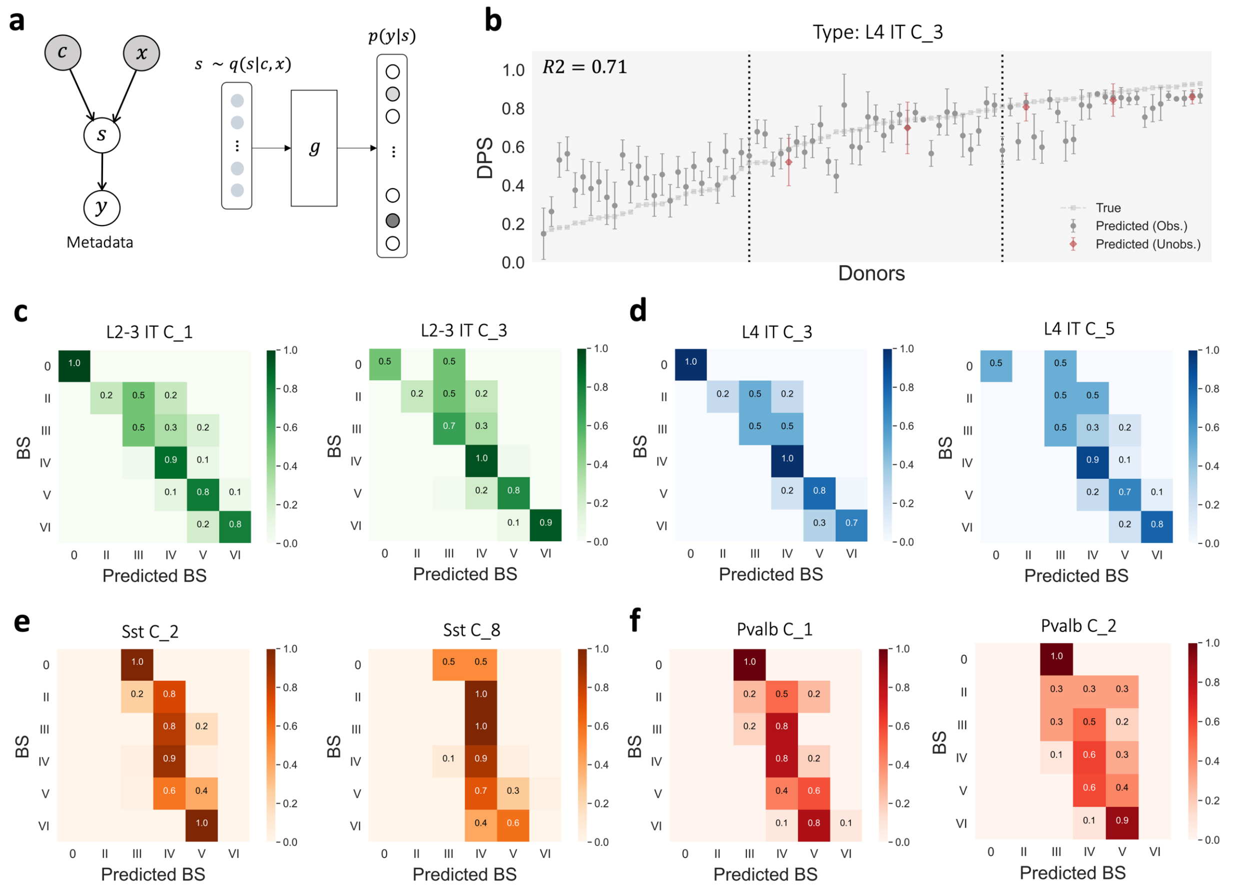


Figure S20: **Role of continuous latent factors in predicting Alzheimer's Disease progression.** (a) The probabilistic graphical model used in the regression analysis of the SEA-AD dataset. Given a cell $\mathbf{x}$, and its identified discrete type, $\mathbf{c}$, MMIDAS generates continuous samples $\mathbf{s}$ using the approximated posterior probability, $q(\mathbf{s}|\mathbf{c},\mathbf{x})$. Through regression, the model learns function $g$, which approximates metadata $y$ based on the input $\mathbf{s}$. (b) Regression analysis for L4 IT C_3 excitatory neurons within the L4 IT subclass. This cell type is among the types that experience significant reduction in abundance during the later stages of the disease (discussed in Results, Fig. 5g). The plot showcases progression of the disease pseudotime (DPS), a measure quantified in [5]. We also present the DPS predicted from the continuous representation inferred by MMIDAS. Dots and bars represent the mean and the standard deviation, respectively, across 10-fold cross-validation. Gray dots correspond to donors whose cells contribute to the training phase of the MMIDAS model. Red dots denote donors entirely excluded from both the training phase of MMIDAS and the regression model, suggesting that both the MMIDAS-derived representation and the regression study exhibit robustness against batch effects. (c-f) Confusion matrices for predicting Braak stages (BS) using the inferred continuous representation for a set of neuronal types in MTG. For this analysis, we train a Random Forest classifier and report the accuracy for each stage. The continuous latent factors inferred for excitatory neurons are influenced by disease progression (Results, Fig. 5h) so that these neurons can effectively predict the Braak stages ((c) and (d)). In contrast, the continuous factors inferred for inhibitory neurons show limited correlation with disease progression and do not reliably indicate Braak stages ((e) and (f)).

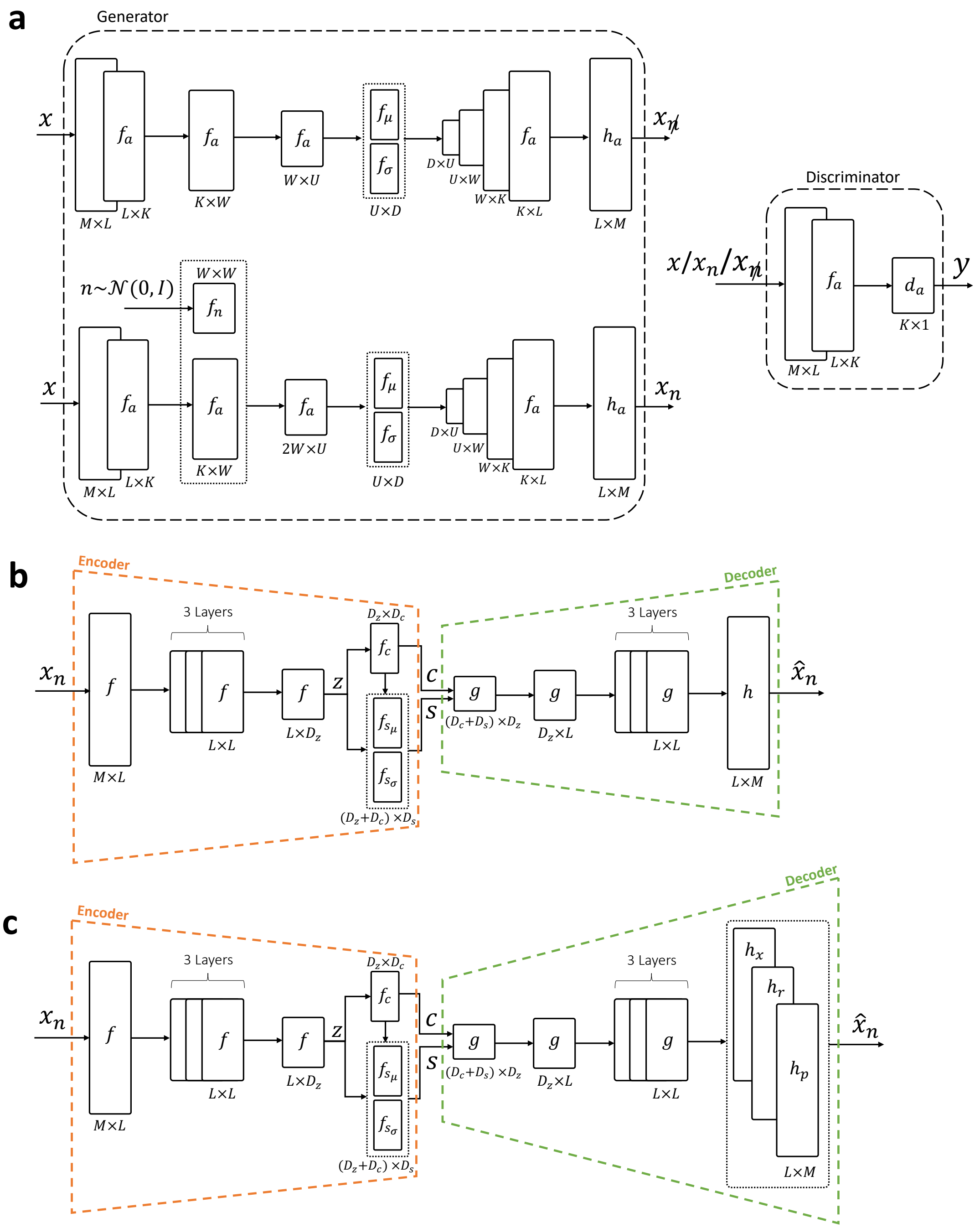


Figure S21: **Architectures of networks employed in MMIDAS.** (a) Type-Preserving Data Augmentation through VAE-GAN Design. Given an input sample $\mathbf{x} \in \mathbb{R}^M$, the generator networks create two samples: $\mathbf{x}_{\not n} \in \mathbb{R}^M$ denotes the reconstructed original sample, while $\mathbf{x_n} \in \mathbb{R}^M$ denotes the augmented sample. In parallel, the discriminator network distinguishes between the original and augmented samples, as outlined in the Methods section. $M$ denotes the dimensionality of data (e.g., number of genes or electrophysiological features. See Methods, Datasets).(b, c) VAE network architectures used for mixture representation learning in MMIDAS for single-cell datasets. In the case of a Gaussian observation noise model, the architecture depicted in (b) is used. When observation noise is modeled by the ZINB distribution, the architecture in (c) is used. The dashed trapezoid in orange represents the encoder unit of the network, and the green dashed trapezoid illustrates the decoder. Here, $z$ represents the low-dimensional embedding obtained in MMIDAS. $c$ and $s$ denote the latent discrete and continuous samples obtained from $q(c|x_n)$ and $q(s|c, x_n)$, respectively (as discussed in Figure 1c). All activation functions and variables are explained in Table S7 and Supplementary Note 14.

| Activation function | Data modality | | |
|---|---|---|---|
| | Smart-seq | 10x | Electrophysiology |
| $f$ | BatchNorm + RELU | BatchNorm + RELU | BatchNorm + ELU |
| $f_c$ | Softmax | Softmax | Softmax |
| $f_{s_\mu}$ | Linear | Linear | Linear |
| $f_{s_\sigma}$ | Sigmoid | Sigmoid | Sigmoid |
| $g$ | RELU | RELU | ELU |
| $h$ | RELU | - | Linear |
| $h_x$ | - | RELU | - |
| $h_r$ | - | Sigmoid | - |
| $h_p$ | - | Sigmoid | - |
| $f_a$ | RELU + BatchNorm | RELU + BatchNorm | - |
| $f_n$ | ELU + BatchNorm | ELU + BatchNorm | - |
| $f_\mu$ | Linear | Linear | - |
| $f_\sigma$ | Sigmoid | Sigmoid | - |
| $h_a$ | RELU | RELU | - |
| $d_a$ | Sigmoid | Sigmoid | - |

Table S7: Activation functions for networks in Fig. S21 across three data modalities. Type-Preserving augmentation is not required for multimodal Patch-seq dataset. Accordingly, data augmenter is not employed for electrophysiology properties.

### Supplementary Note 14

#### Implementation and training parameters

- **Mouse Smart-seq data**
  - Optimizer: Adam with learning rate 1e-3
  - Augmenter network
    * Batch size: 1000
    * Training epochs: $10K$
    * Network parameters: $L = 1000$, $K = 100$, $W = 50$, $U = 20$, $D = 10$
    * Hyperparamters: $\gamma_1 = 0.5$, $\gamma_2 = 0.5$, $\gamma_3 = 0.1$, $\alpha = 0.2$
    * Input layer regularizer: 20% random dropout
  - Mixture VAE network
    * Continuous and categorical variational factors: $D_s = 2$, $D_c = 120^*$
    * Batch size: 5000
    * Training epochs: $53K$
    * Gumbel-Softmax temperature ($\tau$): 1.0
    * Consensus weight ($\lambda$): 1.0
    * Network parameters: $L = 100$, $D_z = 10$
    * Input layer regularizer: 50% random dropout
- **Mouse scRNA-seq 10x data**
  - Optimizer: Adam with learning rate 1e-3
  - Augmenter network
    * Batch size: 1000
    * Training epochs: 1000
    * Network parameters: $L = 1000$, $K = 100$, $W = 50$, $U = 20$, $D = 10$
    * Hyperparamters: $\gamma_1 = 0.5$, $\gamma_2 = 0.5$, $\gamma_3 = 0.1$, $\alpha = 0.2$
    * Input layer regularizer: 20% random dropout
  - Mixture VAE network
    * Continuous and categorical variational factors: $D_s = 3$, $D_c = 110^*$
    * Batch size: 3500
    * Training epochs: $15K$ (Glutamatergic neurons), $40K$ (GABAergic neurons)
    * Gumbel-Softmax temperature ($\tau$): 1.0
    * Consensus weight ($\lambda$): 1.0
    * Network parameters: $L = 100$, $D_z = 10$
    * Input layer regularizer: 50% random dropout
- **Patch-seq**
  - Optimizer: Adam with learning rate 1e-3
  - Batch size: 1000
  - Training epochs: $400K$
  - Gumbel-Softmax temperature ($\tau$): 1.0
  - Consensus weight ($\lambda$): 1.0
  - Mixture VAE network for T modality

    * Continuous and categorical variational factors: $D_s = 2$, $D_c = 100^*$
    * Network parameters: $L = 100$, $D_z = 30$
    * Input layer regularizer: 50% random dropout
  - Mixture VAE network for E modality
    * Continuous and categorical variational factors: $D_s = 3$, $D_c = 100$
    * Network parameters: $L = 100$, $D_z = 30$
    * Input layer regularizer: Additive Gaussian noise, $n \sim \mathcal{N}(0, 0.05)$ and 10% random dropout

- **SEA-AD**
  - Optimizer: Adam with learning rate 1e-3
  - Augmenter network
    * Batch size: 1000
    * Training epochs: 1000
    * Network parameters: $L = 1000$, $K = 100$, $W = 50$, $U = 20$, $D = 10$
    * Hyperparamters: $\gamma_1 = 0.5$, $\gamma_2 = 0.5$, $\gamma_3 = 0.1$, $\alpha = 0.2$
    * Input layer regularizer: 20% random dropout
  - Mixture VAE network
    * Continuous and categorical variational factors: $D_s = 15$, $D_c = 20^*$
    * Batch size: 1000
    * Training epochs: $10K$
    * Gumbel-Softmax temperature ($\tau$): 1.0
    * Consensus weight ($\lambda$): 1.0
    * Network parameters: $L = 100$, $D_z = 30$
    * Input layer regularizer: 20% random dropout

* Here, the initial size of the discrete space, which is the upper bound of the number of clusters in the dataset, is configured to be slightly larger than the suggested number of cell types in the taxonomy (Figs. S14b, S16b, and S17b).In the absence of a taxonomy, one can use an available mouse/human brain atlas to set the maximum expected number of cell types in the dataset.

[2] Diane Bouchacourt, Ryota Tomioka, and Sebastian Nowozin. Multi-level variational autoencoder: Learning disentangled representations from grouped observations. *arXiv preprint arXiv:1705.08841*, 2017.

[3] Nat Dilokthanakul, Pedro AM Mediano, Marta Garnelo, Matthew CH Lee, Hugh Salimbeni, Kai Arulkumaran, and Murray Shanahan. Deep unsupervised clustering with gaussian mixture variational autoencoders. *arXiv preprint arXiv:1611.02648*, 2016.

[4] Emilien Dupont. Learning disentangled joint continuous and discrete representations. In *Advances in Neural Information Processing Systems*, pages 710–720, 2018.

[5] Mariano Gabitto, Kyle Travaglini, Jeannelle Ariza, Eitan Kaplan, Brian Long, Victoria Rachleff, Yi Ding, Joseph Mahoney, Nick Dee, Jeff Goldy, et al. Integrated multimodal cell atlas of alzheimer's disease. *bioRxiv*, 2023.

[6] Rohan Gala, Agata Budzillo, Fahimeh Baftizadeh, Jeremy Miller, Nathan Gouwens, Anton Arkhipov, Gabe Murphy, Bosiljka Tasic, Hongkui Zeng, Michael Hawrylycz, et al. Consistent cross-modal identification of cortical neurons with coupled autoencoders. *Nature computational science*, 1(2):120–127, 2021.

[7] Rohan Gala, Nathan Gouwens, Zizhen Yao, Agata Budzillo, Osnat Penn, Bosiljka Tasic, Gabe Murphy, Hongkui Zeng, and Uygar Sümbül. A coupled autoencoder approach for multi-modal analysis of cell types. In *Advances in Neural Information Processing Systems*, pages 9263–9272, 2019.

[8] Fangjian Guo, Xiangyu Wang, Kai Fan, Tamara Broderick, and David B Dunson. Boosting variational inference. *arXiv preprint arXiv:1611.05559*, 2016.

[9] Yeonwoo Jeong and Hyun Oh Song. Learning discrete and continuous factors of data via alternating disentanglement. In *International Conference on Machine Learning*, pages 3091–3099. PMLR, 2019.

[10] Zhuxi Jiang, Yin Zheng, Huachun Tan, Bangsheng Tang, and Hanning Zhou. Variational deep embedding: an unsupervised and generative approach to clustering. In *Proceedings of the 26th International Joint Conference on Artificial Intelligence*, pages 1965–1972, 2017.

[11] Minoru Kanehisa, Miho Furumichi, Mao Tanabe, Yoko Sato, and Kanae Morishima. Kegg: new perspectives on genomes, pathways, diseases and drugs. *Nucleic acids research*, 45(D1):D353–D361, 2017.

[12] Ozsel Kilinc and Ismail Uysal. Learning latent representations in neural networks for clustering through pseudo supervision and graph-based activity regularization. *arXiv preprint arXiv:1802.03063*, 2018.
